# Loss of *Mycobacterium marinum* ESX-1 genes increase transcription of ESX-6 genes

**DOI:** 10.64898/2026.03.18.712377

**Authors:** Phani Rama Krishna Behra, Malavika Ramesh, B. M. Fredrik Pettersson, Leif A. Kirsebom

## Abstract

Mycobacteria form rough and smooth colonies. The *Mycobacterium marinum* strain 1218S is a smooth colony forming variant isolated from the 1218R strain, which forms rough colonies and is more virulent than 1218S in infecting fish. Genes for the type VII secretion ESX-1 system, which includes mycobacterial virulence genes, have been partially duplicated in *M. marinum* and is refered to as ESX-6. We recently reported that several ESX-1 genes are missing in the 1218S strain. On the basis of the complete genomes of these two and three other *M. marinum* strains we provide insight into strain differences and similarities focusing on 1218R and 1218S, and ESX genes, selected virulence genes, and LOS genes, which are involved in lipo<u>o</u>ligo<u>s</u>accharide synthesis and smooth colony formation. We provide RNA-Seq data for 1218R and 1218S and two other well-characterized *M. marinum* strains suggesting that loss of ESX-1 genes in 1218S results in increased transcript levels of ESX-6 genes. Furthermore, while there is no difference in gene synteny and sequence of LOS genes comparing 1218R and 1218S, with the exception of duplication of *lsr2*, a regulator of LOS genes, in 1218S. Our RNA-Seq data show increased transcript levels of LOS genes in stationary 1218S cells relative to 1218R indicating that transcription and/or RNA degradation of LOS genes influence smooth and rough colony formation. We finally provide data suggesting that Ms1 RNA affect the transcription of LOS genes (and ESX-1 genes), and that loss of ESX-1 genes influence biofilm formation.

## Introduction

Mycobacteria occupy various ecological niches and can be isolated from soil, tap water and ground water and they are divided into <u>s</u>low (SGM) and <u>r</u>apid (RGM) <u>g</u>rowing <u>m</u>ycobacteria. Several mycobacteria cause disease both in humans and animals (land and aquatic), *e.g*. *Mycobacterium tuberculosis* (*Mtb*) the causative agent of tuberculosis (TB). The SGM *Mycobacterium marinum* (*Mmar*) infects a wide array of different fish species, in particular in warm water systems, causing the TB-like disease mycobacteriosis but it can also infect humans causing skin lesions^1–3^. Given that *Mtb* and *Mmar* are phylogenetically close, and the similarities between TB and mycobacteriosis, *Mmar* is used as a model system to study mycobacterial pathogenicity^3–6^.

Mycobacteria (*e.g*. *Mtb* and *Mmar*) carry genes for type VII secretion systems refered to as ESX-1, −2, −3, −4 and −5 where ESX-1, ESX-3 and ESX-5 genes include major virulence factors^7–17^. In *Mmar* the ESX-1 region is partly duplicated, and is refered to as ESX-6^7,18^. We recently reported the genome sequences for several *Mmar* strains including the complete and draft genomes of the *Mmar* strains 1218R and DE4381 (also refered to as 1218S), respectively^18^. The *Mmar* strain 1218S (*Mmar*^1218S^) is a smooth colony forming variant isolated from 1218R (*Mmar*^1218R^), and it is less virulent than 1218R in infecting Japanese medaka^18, see^ ^also^ ^19^. Compartive genomics revealed that 1218S lacks several ESX-1 genes, *espF*_2, *espG*1_2, *espH*, *eccA*1_3, *eccB*1_5, *eccC*a_1__4, *eccC*b_1_, PE35 (gene id 1218R_05483), PPE68 (gene id 1218R_05484) and *esxB*_3, which encodes for CFP-10 (following *Mtb*^H37Rv^ gene annotation). Among these genes, the 1218S ESX-6 region encompasses homologs to *eccB*1, PE35, PPE68 and *esxB*_3 in addition to other ESX-1 homologs^7,18^. Given these differences between 1218R and 1218S, and the draft genome statues of the *Mmar*^1218S^ genome, we were interested in to investigate this further. Here we provide the complete 1218S genome showing that parts of *espF*_2 and *esxB*_3 is present while the other ESX-1 genes mentioned above are missing as previously reported^18^. Because ESX-1 genes are duplicated in *Mmar* (see above), where the ESX-1 and ESX-6 *esxA* and *esxB* homologs are well conserved, we were also interested in to get insight into the expression of the ESX-1 and ESX-6 genes in 1218R and 1218S. In particular, whether lack of ESX-1 genes in 1218S affect ESX-1 (for the genes that are still present) and ESX-6 mRNA levels. Hence, we generated and analyzed transcription (RNA-Seq) profiles of 1218R and 1218S in exponential growing and stationary cells focusing on ESX-1 and ESX-6 gene transcripts. In this context, we also analysed transcript levels of ESX-3, ESX-4, ESX-5 and selected virulence genes. For comparison, we include the transcription profiles for two other *Mmar* strains, the type strain strain CCUG 20998 (*Mmar*^CCUG^)^18,20^ and the M strain (*Mmar*^M^).

As stated above the *Mmar*^1218S^ strain forms smooth (S) colonies while 1218R forms rough (R) colonies. It is well documented that mycobacteria form R and S colony morphotypes due to differences in the structure of the outer membrane/cell wall for a review ^see^ ^e.g. 21^. This difference influence virulence, where R morphotypes appear to be more virulent^see^ ^e.g. 18,22–24^ and Refs therein. For mycobacteria, genes related to the production of glycopeptidolipids (GPL) and lipooligosaccharides (LOS) have been shown to be responsible for generating S and R morphotypes^21,23–31^. We were therefore also interested in to understand whether in particular LOS genes, and/or the transcription of these genes, differ in the 1218R and 1218S *Mmar* strains. Finally, because ESX-1 and LOS might influence sliding motility and biofilm formation^25,32^ provided the incentive to study 1218R and 1218S biofilm formation with and without a plasmid carrying *espF*_2, *espG*1_2 and *espH*, *i.e*. genes not present in the 1218S strain.

Our data show that the absence of ESX-1 genes in 1218S resulted in higher ESX-6 mRNA levels. Hence, an increased transcription level of ESX-6 genes appears to compensate for the loss of ESX-1 genes in 1218S. Moreover, our RNA-Seq data for four different *Mmar* strains show that in particular ESX-1 and ESX-5 mRNA levels are higher in stationary phase compared to exponentially growing *Mmar* cells, while ESX-3 gene transcripts are lower. With respect to the LOS gene locus, we detected no difference in the number of genes, or in their sequences. However, in 1218S the *lsr2* gene, which is involved in the regulation of the expression of LOS genes (for references see above), is duplicated. Furthermore, the transcript levels of several LOS genes were increased in stationary 1218S cells relative to 1218R. We discuss these findings, and that the transcription of LOS genes (and ESX-1 genes) is influenced by Ms1 RNA and plausibly also by other non-coding RNAs^33^, in relation to the formation of smooth *Mmar* colonies. We also present data for other virulence genes of interest that are present/absent comparing 1218R and 1218S, and that their mRNA levels differ, and that 1218S form less robust biofilm than 1218R. Together our data provide insight into, why 1218S is less virulent than 1218R and formation of smooth colonies in *Mmar*.

## Results

We previously reported a genomic analysis of different *M. marinum* strains, which included the R and S colony forming (Fig 1a) *Mmar* strains 1218R (*Mmar*^1218R^) and DE4381, a derivative of 1218S, and refered to as 1218S or *Mmar*^1218S^ ^18^. To get deeper insight into the differences between 1218R and 1218S we generated the complete genome sequence for 1218S (see Materials and Methods). The 1218R and 1218S complete genomes were subsequently used in a comparative genomic analysis along with the complete genomes of three *Mmar* strains, CCUG 20998 (*Mmar*^CCUG^; type strain), M (*Mmar*^M^) and ATCC 927^T^ (*Mmar*^ATCC^)^7,28,34^. Firstly, we discuss the genomic differences comparing 1218R and 1218S and then we focus on virulence genes, ESX genes (specifically ESX-1 genes), and LOS genes. Following this we examine the transcript levels of selected virulence genes (discussed in Supplementary information), ESX genes, in particular ESX-1 and ESX-6 genes, LOS genes and genes encoding relevant transcriptional regulators in exponentially growing and stationary cells using RNA-Seq data focusing on 1218R and 1218S (see Materials and Methods).

**Figure 1.**
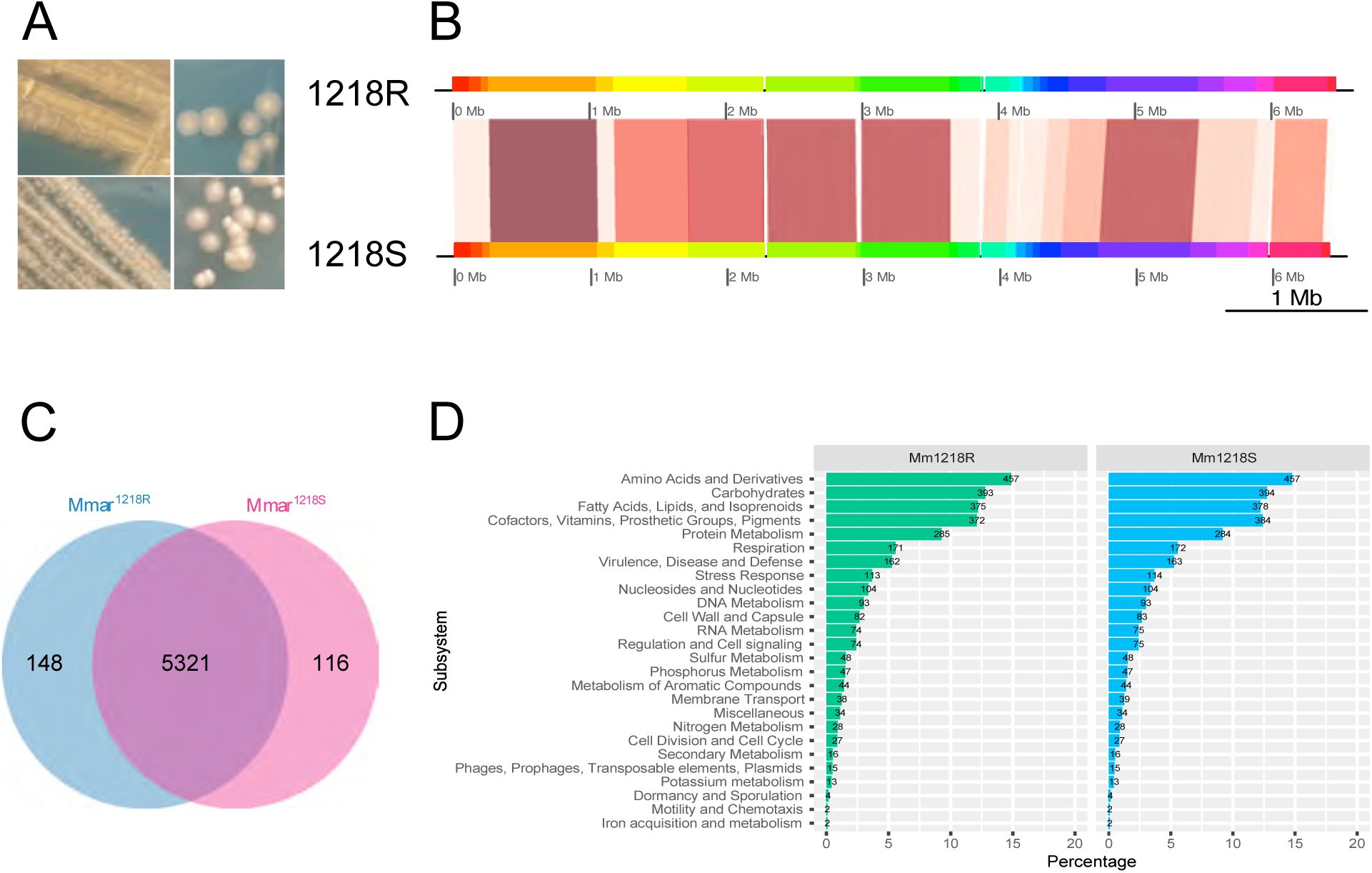
Colony morphologies, overview of the 1218R and 1218S genomes, and analysis of CDS and functional classifications. (a) Colony morphologies of 20 days old cultures of 1218R (top row) and 1218S (bottom row) grown on 7H10 at 30°C. (b) Whole-genome alignment for 1218R and 1218S where horizontal blocks represent the genomes as marked while vertical “blocks” correspond to homologous regions. White gaps represent insertions/deletions. For details see the main text. (c) Venn diagram showing the presence of common and unique annotated genes for 1218R and 1218S. (d) Functional classification of 3073 and 3095 core genes in 1218R and 1218S, respectively. For comparison of 1218R, 1218S, *Mmar*^CCUG^, *Mmar*^ATCC^ and *Mmar*^M^, see Figure S1a-c.

### Comparative genomics of five M. marinum strains - Mmar^1218R^, Mmar^1218S^, Mmar^CCUG^, Mmar^ATCC^ and Mmar^M^

The sizes of the complete 1218R and draft 1218S genomes were reported to be 6467358 and 6324388 bp, respectively^18^. *Mmar*^1218S^ DNA was prepared and PacBio sequencing technology generated a single scaffold representing the complete 1218S genome encompassing 6410821 bp, which is larger than the previously reported draft genome (Fig 1b and Table S1a-d for description of genes; see also Refs 18 and 35). Hence, the 1218R genome is 56537 bp larger than that of 1218S, disclosing a reduction in the size of the 1218S genome.

Alignment of the complete 1218R and 1218S genomes revealed differences in a few regions corresponding to *e.g*., locations near positions 2.28, 2.98, 3.85 and 5.96 Mbps (Fig 1b, white marked regions; see Table S1d and below for rearrangements of individual genes). Comparison with the complete genomes of the *Mmar*^CCUG^, *Mmar*^M^ and *Mmar*^ATCC^ suggested that both 1218R and 1218S are more similar to *Mmar*^CCUG^ and *Mmar* ^ATCC^ than to *Mmar*^M^. This is in agreement with that the M strain is a *Mmar* subspecies separated from the “ATCC” lineage/Aronson type (Fig S1a, see also Table S1e-g)^18^. The *Mmar*^CCUG^ and *Mmar* ^ATCC^ are similar given their common origin^36^. Moreover, comparing 1218R and 1218S with *Mmar*^CCUG^ and *Mmar*^M^ revealed rearrangement/inversion of the 1.71-1.76 Mbp-region in 1218R and 1218S, which corresponds to the 4.81-4.86 and 4.62-4.67 Mbp-regions in *Mmar*^CCUG^ and *Mmar*^M^, respectively (Fig S1a; for 1218R and 1218S the regions are marked with * blue boxes).

We identified 5321 <u>c</u>oding <u>d</u>etermining <u>s</u>equences, CDS, that are common in 1218R and 1218S (Fig 1c; see also Tables S1a, b), 49 correspond to tRNA and 47 non-coding (nc) RNA genes in both strains. Including *Mmar*^CCUG^, *Mmar*^ATCC^ and *Mmar*^M^ in the analysis revealed that 4557 CDS are common among these five strains (Fig S1b; see also Tables S1a, b, e-g). *Mmar*^CCUG^ encodes for 47 tRNAs and 48 ncRNAs while *Mmar*^M^ and *Mmar*^ATCC^ both have 46 tRNA and 49 ncRNA genes (Rfam annotated ncRNAs^18,35^).

The RAST tool predicted functional classification of >3000 CDS in 1218R and 1218S (Fig S1c), and did not suggest any significant difference comparing the CDS (but see below). The RAST functional classification can overlook genes due to mutations such as single nucleotide changes and deletions. Given that we were primarily interested in comparing 1218R and 1218S, we manually edited and updated the functional classification for these two strains. As expected, the classification profiles were very similar given the close relation between these two strains (Figs 1d).

Considering unique genes (after editing for <u>s</u>ingle <u>n</u>ucleotide <u>p</u>olymorphisms, SNP, duplications, shift of gene location and missing annotations), we identified 148 and 116 unique genes in 1218R and 1218S, respectively (Fig 1c; Table S1d). When we compared the functional classification profiles including the other three *Mmar* strains we detected roughly a two-fold higher number of genes belonging to the “Cell division and Cell cycle” in *Mmar*^CCUG^ and higher numbers in the “RNA metabolism” category in *Mmar*^M^ relative to the other strains (Fig S1c; note, numbers shown for the different strains represent unedited functional classification).

### Virulence genes and ESX genes comparing 1218R, 1218S, Mmar^CCUG^, Mmar^ATCC^ and Mmar^M^

The smooth variant 1218S showed a four-fold reduction in virulence relative to the rough 1218R strain in infecting Japanese medaka^18^. Hence, using the complete genomes we mapped and classified virulence-related genes/factors for functions in these two strains. As shown in Fig 2a, 1218R encodes for 333 virulence factors (using the VFanalyzer tool, VFDB database), which is eight more than in 1218S. This was expected since the ESX-1 region encodes for several known virulence-related genes, which are not present in 1218S (see also below). The RAST tool, however, predicted 160 and 166 virulence-related genes in 1218R and 1218S, respectively, distributed in subsystem catagories (Fig 2b). The difference relates to six additional genes in the “Cofactors, Vitamins, Prosthetic Groups, Pigments” subsystem category in 1218S. The predicted virulence-related genes are listed in Table S2.

**Figure 2.**
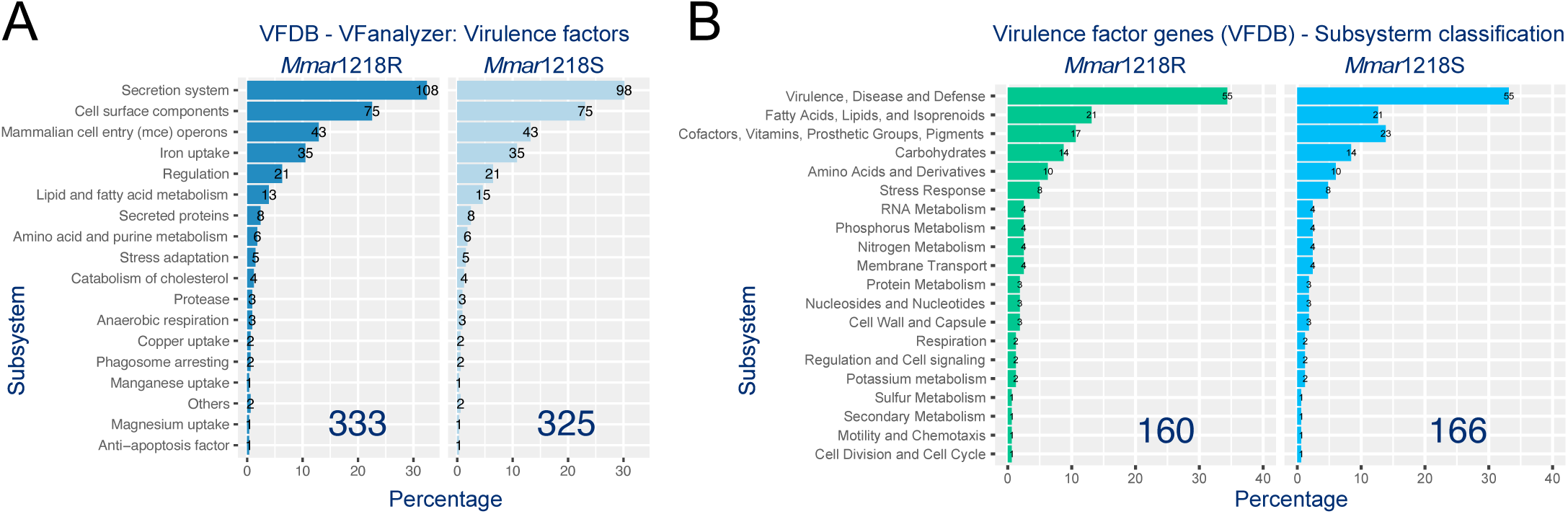
Prediction of virulence-related genes in 1218R and 1218S. (a) Identification and classification of virulence-related genes in 1218R and 1218S using the VFanalyzer tool (the VF data base, VFDB). (b) Subsystem classification, RAST subsystem categories, of genes identified in the VFDB analysis in (a).

One difference between 1218R and 1218S is a consequence of the deletion of 10 genes in the ESX-1 region, encompassing *espF*_2 and *esxB*_3^18^. However, the complete 1218S genome uncovered that *espF*_2 and *esxB*_3 (corresponding to the CFP-10 gene) are truncated (Fig 3a, b). The most likely reason to this is that for the draft 1218S genome *esxB*_3 is positioned close to a scaffold that affected the assembly. Nevertheless, the deletion in the 1218S ESX-1 region covers nucleotides corresponding to nucleotides 6408265-6418995 in the 1218R genome generating truncated *espF*_2 and *esxB*_3 genes in 1218S (Figs 3a, b and S2a). For *espF*_2, the Shine-Dalgarno (SD) sequence and the translational start codon AUG is still present and in frame with a UGA stop codon that overlaps the *esxA*_3 (the ESAT-6 gene) AUG translation start codon. Translation of the corresponding mRNA would result in a 98 amino acids long peptide with a different sequence compared to the wild-type EsxB_3 protein (CFP-10; Figs 3b and S2a; no potential translational stop codon could be detected between the EspF start codon and the identified stop codon; see also below). We also detected an insertion of a C in the gene downstream of *esxA*_3 in 1218S (1218S gene id: 05455), which would change the translational reading frame (Fig S2a). However, this insertion is likely due to a sequencing error.

**Figure 3.**
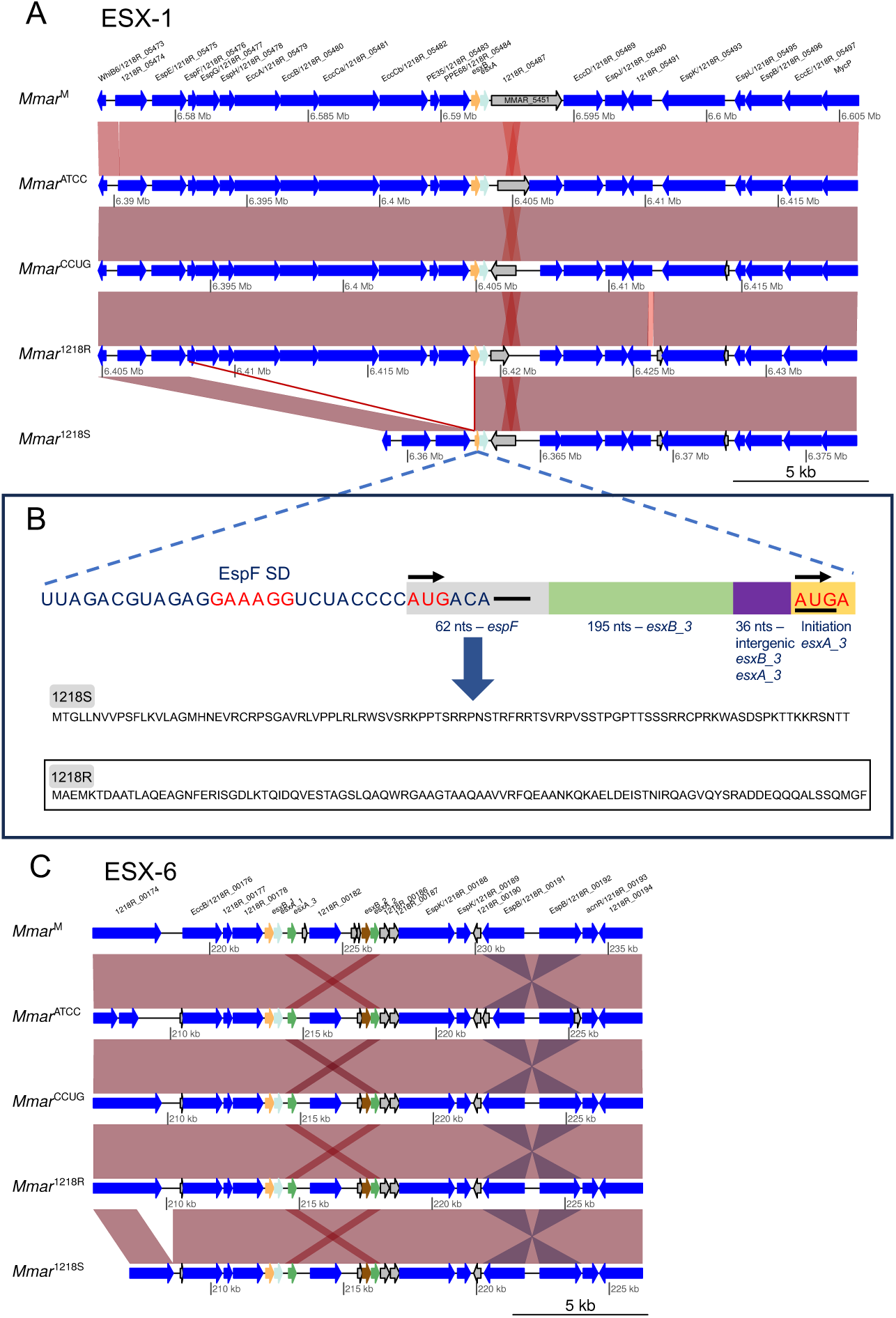
Gene synteny for ESX-1 and the partially duplicated ESX-6 gene clusters in five different *M. marinum* strains. (a) Gene synteny for the ESX-1 gene cluster in five different *M. marinum* strains as indicated. Arrows represent genes where blue colours mark genes with known function, grey hypothetical genes, and orange and light blue *esxB*_3 (*esxB*) and *esxA*_3 (*esxA*), respectively. The vertical brown boxes highlight homologous genes. The red lines mark the deleted ESX-1 genes in 1218S, see also panel (b). (b) Highlight of the predicted “EspF-EsxB” mRNA in 1218S where nucleotides marked red represent the predicted EspF Shine-Dalgarno (SD) sequence and translation initiation codon AUG, and EsxA AUG codon, which overlaps a predicted translational termination codon UGA. The grey box refers to EspF mRNA residues, green box “EsxB mRNA” residues and purple residues originating from the *esxB*_3 and *esxA*_3 intergenic region. Below the predicted peptide sequence as a result of translation of the “EspF-EsxB” mRNA in 1218S. The boxed peptide sequence corresponds to the wild-type 1218R EsxB_3 peptide. (c) Gene synteny for the ESX-6 gene cluster in five different *M. marinum* strains as indicated. As in (a) the arrows represent genes and the colour code as in (a) with the addition that the orthologous *esxA*_3 and *esxA*_2 are marked in green and *esxB*_2 in brown.

Moreover, relative to 1218R the 1218S ESX-6 region carries a deletion of 1371 nucleotides, beginning at the genomic position 208875 in a PE family protein gene (1218R/1218S gene id: 00174) and ending at 210246 in the “00174/00175-intergenic region” (positions refer to 1218R). This gives a C-terminal truncated PE family protein in 1218S (Fig 3c and Fig S2b). We also noted nucleotide deletions in the ESX-6 region, one in 1218S and one in 1218R (*espB*_1 gene id: 00190 1218S and *espB*_2 gene id: 00192 in 1218R; Fig S2b), which might be related to sequencing errors. For the other three ESX regions (ESX-3, ESX-4 and ESX-5), the gene sequences were identical comparing the 1218R and 1218S genomes (Figs S2c-f).

In addition to the deletion of genes in the ESX-1 region (and nucleotides in ESX-6) in 1218S we detect differences in other regions where some encode for virulence factors (Tables S1d). For example, *panC* and *panD* are duplicated in 1218S (relative to 1218R; Fig 4a, see also below) and these two virulence genes are involved in the synthesis of pantothenate (vitamin B5; 37 Suresh et al., 2020). The *lsr2* and *lysS* genes are located close to *panC* and *panD*. These four genes, together with other neighboring genes are duplicated in 1218S (Fig 4a; see also below). Interestingly, a region in the middle of a putative ribonuclease J gene [Fig S3a; 1218R/1218S gene id: 01974/01981(82)] is deleted in 1218S. Deletion of genes in 1218S were also identified in other regions such as the ATP-dependent RNA helicase RhlE gene (Fig S3b), and in several dimodular nonribosomal peptide synthase, *dhbF*, genes (Table S1d). We also detected re-allocations of genes, *e.g*. *rbmH* (flavin reductase), which is postioned close to *rhlE* in 1218S (Fig S3b), and gene duplications in 1218R (Table S1d). Moreover, in 1218S *ppsB*_3 is annotated as one gene, while in 1218R it is annotated as two genes (*ppsA*_1 and *ppsB*_3) due to SNP (Fig S3c). For further differences and details, see Tables S1d and S2.

**Figure 4.**
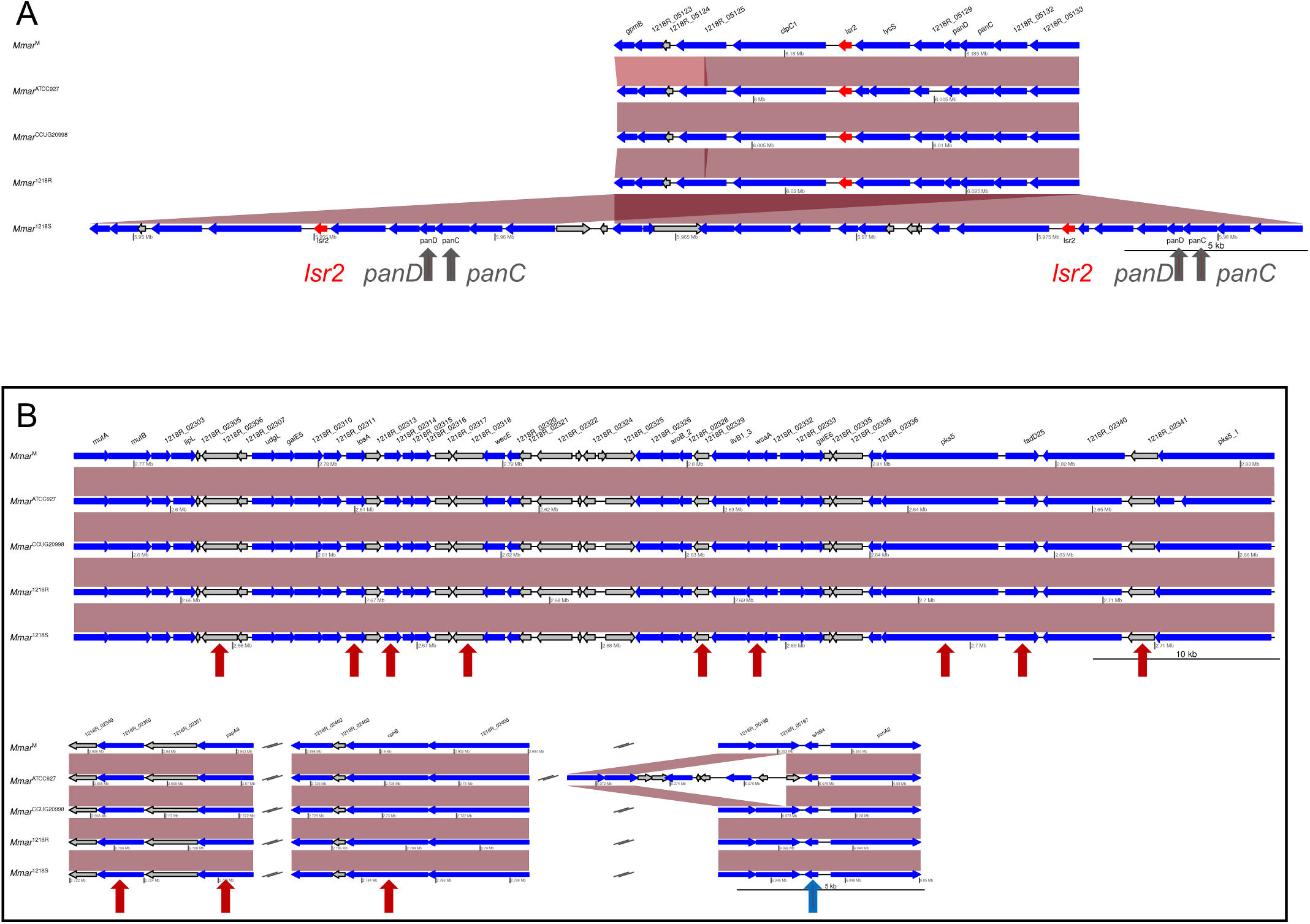
Gene synteny for the *lsr2* region and the LOS genes in five different *M. marinum* strains. (a) Gene synteny of *lsr2* region where *lsr2* is marked in red. The duplication of *panD* and *panC* in 1218S are marked with black arrows. The blue and grey arrows represent genes. For colour code, see Figure legend 3a. (b) Gene synteny of LOS genes and the *whiB*4 region. The blue and grey arrows represent genes, for colour code see Figure legend 3a. The red arrows mark genes, when mutated result in formation of rough colonies in *Mmar*^M^, see main text for details. The blue veritical arrow whiB4. For details see main text and Figure S5.

Examination of the predicted number of virulence-related genes in the five *M. marinum* strains (*Mmar*^1218R^/1218R, *Mmar*^1218S^/1218S, *Mmar*^CCUG^, *Mmar*^ATCC^ and *Mmar*^M^) suggests that *Mmar*^ATCC^ has 250 virulence genes, while *Mmar*^M^ encodes for the highest number, 345 virulence genes, relative to the other four strains (Fig S4; using the VFanalyzer tool, VFDB database; see also Table S2). The difference between the five strains is mainly due to the number of genes in the subsystem categories “Secretion system” and “Cell surface components”. For *Mmar*^CCUG^, the lower number of virulence classified genes compared to 1218R appear to be related to that 1218R has extra genes in the category “Secretion system”, *e.g*. genes annotated as *ppe68*, *ppe4*, *pe5*, *esxG* (ESX-3), and that *Mmar*^CCUG^ lacks the ESX-3 *esxH* gene (Tables S1e and S2). With respect to *Mmar*^ATCC^, the lower number of annotated virulence genes (Fig S4) pertains to for example lower numbers of ESX genes relative to the other *Mmar* strains. For other differences, see Tables S1f and S2a. The higher number of virulence-related genes in *Mmar*^M^ relates to an increased number of genes in the “Secretion system” and “Cell surface components” categories. Also, *Mmar*^M^ has a higher number of “Copper uptake” genes compared to the other strains (Fig S4, Tables S1g and S2a).

### LOS genes - comparing 1218R and 1218S

When grown on solid media 1218R and 1218S form rough (R) and smooth (S) colony morphotypes (see above; Fig 1a). This shift in *M. marinum* colony appearance is related to genes associated with the biosynthesis of lipooligosaccharides, LOS (for Refs see introduction). The genomes of 1218R and 1218S both contained the same number of LOS genes in the LOS locus and we detected no differences in gene synteny and gene sequences comparing these two genomes (Figs 4a, b and S5a). When comparing genes encoding for relevant regulators in 1218R and 1218S suggested to affect S and R colony formation, no differences in the coding or in the regulatory regions of *whiB*4, *lsr2* and *pknL* (gene ids: 1218R_03215 and 1218S_03210), except that *lsr2* is duplicated in 1218S (see above), were detected (Figs 4a and S5b-d; the gene sequences of the two *lsr2* genes (1218S) and the 1218R *lsr2* were the same). Hence, no apparent rational could be acertained at the LOS gene level, or in the examined regulatory genes (except *lsr2*), to be the reason to the observed difference in the 1218R and 1218S colony morphotypes. We also noted that no difference was detected comparing the number of LOS genes and the gene synteny in the five *M. marinum* strains (Fig 4a; note, *Mmar*^ATCC^ has a few additional genes upstream of *whiB*4).

## Transcript levels of ESX genes, selected virulence genes and LOS genes under different growth conditions

To get deeper insight into the differences between 1218R and 1218S we isolated total RNA from exponentially growing and stationary cell cultures. The isolated RNA was subjected to RNA-Seq (see Materials and Methods). Firstly, we were interested in to understand whether deletion of ESX-1 genes influenced the transcription of ESX-6 genes as well as to compare the transcription levels of ESX-1, ESX-3, ESX-4 and ESX-5 genes in 1218R and 1218S where ESX-1, ESX-3 and ESX-5 are associated with virulence. We also included *espACD* since *espA* and *espC* encode for two proteins that are secreted through ESX-1, where EspC forms a filamentous structure on the cell surface^e.g. 16,17,38,39^. RNA-Seq data for other virulence genes, see Supplementary information (Fig S6a-g and Table S3a, b). Futhermore, the transcription regulators WhiB6, EspR and EspM as well as the two-component system PhoPR influence the transcription of ESX-1 genes and *espACD*. We therefore included *whiB*6, *espR*, *espM*, and *phoPR* in our analysis. Secondly, given that we did not detect any difference comparing LOS gene sequences and synteny in 1218R and 1218S (except *lsr2*, see above) we also asked whether the difference in their colony morphologies is related to the transcript levels of the LOS genes in exponential growing and stationary cells.

The transcript reads were mapped to the respective genome and we used the 1218R genome as reference to compare mRNA levels in *Mmar*^1218S^, *Mmar*^CCUG^ and *Mmar*^M^ relative to *Mmar*^1218R^ (see Materials and Methods). Irrespective of strain, the mRNA levels were calculated such that the distribution in exponential and stationary growth phase as percentage, which corresponds to, *e.g*. individual ESX mRNA levels relative to the sum of the levels for all ESX gene transcripts (including *espACD*, *espR* and *whiB*6) or expressed as TPM values, transcripts <u>p</u>er <u>m</u>illion^40^. While change corresponds to mRNA levels comparing exponential and stationary growth phase for the different strains^20,41^. For comparison, we also analyzed the mRNA levels for the genes discussed above and virulence genes in exponentially growing and stationary *Mmar*^CCUG^ and *Mmar*^M^ cells (Figs 5, S6 and S7; Table S3a, b; Supplementary information).

**Figure 5.**
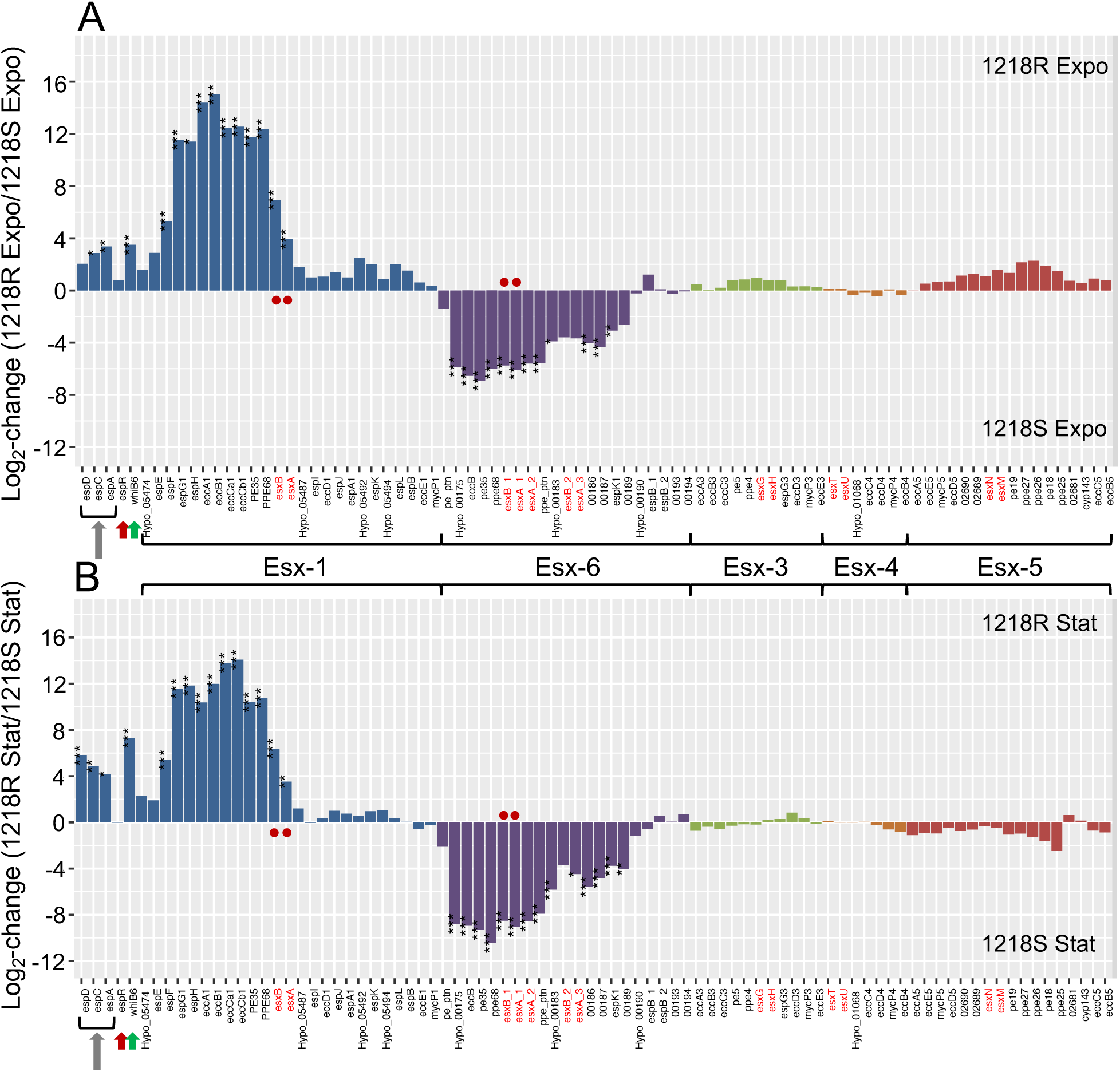
Transcript levels of ESX genes comparing 1218R and 1218S. (a) Transcript levels in 1218R and 1218S exponentially growing cells as indicated. (b) Transcript levels in 1218R and 1218S stationary cells as indicated. The *esxBA* paralogous are indicated in red while the *esxBA* homologous in ESX-1 and ESX-6 are marked with red dots. The arrows mark espACD (grey), espR (red) and whiB6 (green). Statistical significance, see Materials and Methods; *p-value < 0.05; **p-value < 0.01; ***p-value < 0.001. For the distribution and log_2_-fold change in 1218R, 1218S, *Mmar*^CCUG^ and *Mmar*^M^, respectively, see Figure S7a-l.

### Analysis of ESX mRNA levels comparing 1218R and 1218S

The transcript levels of the ESX-1 and ESX-6 genes (and ESX-3, ESX-4 and ESX-5 genes) in 1218R and 1218S exponential and stationary cells are shown in Fig 5 (1218R and 1218S; see also Fig S7a-f). The *Mmar*^CCUG^ and *Mmar*^M^ data are shown in Fig S7g-l (see also Table S3a, b).

Overall, transcript levels for the genes belonging to the different ESX systems were detected both in exponentially growing and stationary 1218R and 1218S cells albeit for ESX-4 genes the levels were low (Figs 5 and S7a-f). We detected significant differences comparing ESX-1 and ESX-6 mRNA levels in 1218R and 1218S: irrespective of strain and growth phase ESX-1 mRNA levels (note, genes present in 1218S) were high and levels originating from ESX-6 were low in 1218R while the opposite was observed for 1218S (Figs 5 and S7a-f). Comparing ESX-1 and ESX-6 transcript levels in exponential and stationary 1218R and 1218S cells revealed that the levels are higher in 1218S cells irrespective of growth phase (Fig 5). We conclude that the deletion of the ESX-1 genes in 1218S (*espF*_2 to *esxB*_3; see above), results in a reduction of ESX-1 transcripts from the remaining genes while ESX-6 mRNA levels increase.

Similar levels and changes for the ESX-1 transcripts were also detected in exponential and stationary *Mmar*^CCUG^ and *Mmar*^M^ cells, *i.e*. increased levels in stationary cells (Fig S7g-l). Albeit that ESX-6 transcripts were also low in *Mmar*^CCUG^ and *Mmar*^M^ cells irrespective of growth phase, the levels increased in stationary cells for both these strains, which is different to what we observed for 1218R (cf. Figs S7c, i and l).

The ESX-1 locus encompasses the major virulence genes, *esxB*_3 (CFP-10; protein name in *Mtb*) and *esxA*_3 (ESAT-6; protein name in *Mtb*). In *M. marinum*, ESX-1 genes have been duplicated generating ESX-6 where the *esxB*_ 3/*esxB*_ 1 and *esxA*_3/*esxA*_1 coding-sequences are identical^6^ (Figs 3a, c and S2 a, b). In 1218R and 1218S, the mRNA levels of these genes were the most abundant transcripts originating from the ESX-1 and ESX-6 regions, respectively (marked in red in Figs 5 and S7a, b; note, *esxB* and *esxA* correspond to *esxB*_3 and *esxA*_3 in ESX-1 and are named *esxB*_1 and *esxA*_1 in ESX-6, respectively). The *esxB*_3 and *esxA*_3 mRNA levels were also the highest in *Mmar*^CCUG^ and *Mmar*^M^ (Fig S7g-l). We emphasize that deletion of ESX-1 genes in 1218S result in a truncated *esxB*_3 (see above) and we did detect reads (transcript) overlapping *esxB*_3 in 1218S. This transcript started upstream of the truncated *espF*_2 (see above; Figs 3a, b and S8) and the resulting sequence is suggested to be different compared to the 1218R *esxB*_3 transcript (Fig 3b). If the “EspF_2-EsxB_3” mRNA in 1218S is translated this would result in a peptide with a different amino acid composition relative to wild type EsxB_3 (Fig 3b; see above and the discussion).

Our data also showed that the transcript levels of the *esxB*_3 and *esxA*_3 (ESX-1) paralogs *esxG* and *esxH* (ESX-3), and *esxN* and *esxM* (ESX-5) were the most abundant among the transcripts from these ESX regions (Fig S7a-f; for the ESX-3 and ESX-5 paralogs in *Mmar*^CCUG^ and *Mmar*^M^ see Fig S7g-l). We also note that ESX-3 mRNA levels are lower while ESX-5 mRNA levels are higher in stationary cells compared to exponential growing cells irrespective of strain (Figs S7; Table S3a, b).

The mRNA levels for the virulence genes *espACD* were higher in 1218R than in 1218S irrespective of growth phase and the levels increase in stationary cells relative to exponential growing cells in both strains but less increase was detected in 1218S (Figs 5 and S7a-f; Table S3a, b). Together this is consistent with that the absence of the ESX-1 secretion system influence transcription of *espACD*.

Comparing *Mmar*^CCUG^ and *Mmar*^M^ *espACD* mRNA levels with the 1218R levels, revealed that the levels were higher in exponential growing 1218R cells relative *Mmar*^CCUG^, while in stationary cells the difference was modest, with a higher level only for the *espD* transcript in 1218R (Table S3b). For *Mmar*^M^, the *espACD* mRNA levels were higher in exponential growing cells relative to 1218R. This was also the case comparing stationary cells with the exception of *espD* mRNA, which is higher in 1218R (Table S3b). Higher *espACD* mRNA levels are also detected in stationary *Mmar*^CCUG^ and *Mmar*^M^ cells relative to exponentially growing cells (Fig S7g-l; Table S3a).

### Transcript levels of regulators of ESX-1 – WhiB6, EspM, EspR and PhoPR

The genes *whiB*6, *espM* (*whiB*6 is located immediately upstream of *espM* in ESX-1; Fig 3) and *espR* encode for transcription factors, which directly or indirectly regulate the expression of ESX-1 genes and *espACD*^16,38,42,43^. The two-component regulator PhoPR also influence the expression of ESX-1 genes and *espACD* through its regulation of *espR* and *whiB*6^38^. Data also suggest that the recently identified transcription factor EspM binds to the intergenic region between *whiB*6 and *espM* and regulates the expression of *whiB*6^43^. Together with the finding that linked *whiB*6 expression and the presence of ESX-1 in the membrane^15,44^ guided us to analyse transcript levels of these regulators in the four *Mmar* strains, focusing on 1218R and 1218S. In addition, recent data indicated that WhiB4 binds in the *whiB*6-*espM* intergenic region in *M. tuberculosis*^45^. The levels of *whiB*4 mRNA will therefore also be discussed, see below in the context of transcription of the LOS genes. The data are shown in Figs 5 and S7, and Table S3.

The *whiB*6 mRNA levels were higher in 1218R than in 1218S, both in exponentially growing and stationary cells (Fig 5; Tables S3b and S4). This difference is partly due to a decrease in the *whiB*6 transcript level in 1218S stationary cells while a slight increase (if any) was detected in 1218R (Fig S7a-f; Tables S3a and S4). Higher transcript levels were also detected for *espM* in 1218R relative to 1218S irrespective of growth phase. However, for both strains the EspM mRNA levels increased in stationary cells relative to exponentially growing cells with less increase in 1218S. Together, it appears that the deletion of “*espF*_2 – *espB*_3” (see above) in 1218S influences the transcription of both *whiB*6 and *espM*.

When we compared WhiB6 and EspM mRNA levels in *Mmar*^CCUG^ and *Mmar*^M^ relative to 1218R we detected higher mRNA levels (WhiB6) in exponentally growing *Mmar*^CCUG^ cells and no significant difference in WhiB6 levels in stationary cells. No change in EspM mRNA levels was detected in *Mmar*^CCUG^ cells relative to 1218R irrespective of growth phase. Comparing 1218R and *Mmar*^M^, both WhiB6 and EspM mRNA levels were higher in 1218R stationary cells (Tables S3b and S4). With respect to WhiB6 mRNA, the level was higher in *Mmar*^M^ in exponential growing cells relative to stationary cells, *i.e*. similar to 1218S (cf. Fig S7; Tables S3a and S4). Furthermore, as for 1218R the EspM mRNA levels was higher in stationary cells relative to exponential growing *Mmar*^CCUG^ cells while no difference was detected for *Mmar*^M^ (cf. Fig S7; Table S3a). Thus, the relative levels of *whiB*6 and *espM* transcripts in exponential and stationary cells depends on *Mmar* strain. As discussed above, *Mmar*^M^ represents a different lineage than the other three strains^18^; unpublished data and this might be related to the observed differences.

For *espR*, the transcript level is modestly higher in exponentially growing 1218R cells relative to 1218S, while no difference was detected in stationary cells (Fig 5; Tables S3b and S4). For both strains, *espR* transcript levels were higher in stationary cells, which is consistent with higher *espACD* levels in stationary cells (Table S3a). Furthermore, small differences were observed comparing *espR* transcript levels for 1218R with *Mmar*^CCUG^ and *Mmar*^M^ except comparing stationary 1218R and *Mmar*^M^ cells where the levels are higher in 1218R cells (Tables S3b and S4). This might be related to that no change in the *espR* transcript level was detected comparing exponentially growing and stationary *Mmar*^M^ cells, while for the other three strains we detected increased levels (Fig S7; Tables S3a and S4a).

For PhoPR mRNA levels, we detected only small differences (if any) comparing 1218R with 1218S or *Mmar*^CCUG^, except for stationary *Mmar*^CCUG^ cells where a modestly higher level was detected (Table S3b). In contrast, the levels in *Mmar*^M^ are higher than in 1218R irrespective of growth phase. Also, the levels are higher in exponentially growing *Mmar*^M^ cells while less differences (if any) were detected for the other three strains (Table S4). These findings might reflect that the regulation of the expression of *phoPR* varies and depends on the strain where 1218R, 1218S and *Mmar*^CCUG^ belong to the same and the M strain to a different *Mmar* lineage^18^, unpublised data.

### Transcript levels of LOS genes involved in formation of rough and smooth colony morphology

Focusing on 1218R and 1218S, and as discussed above, 1218S was isolated as a smooth variant of 1218R (Fig 1a). Based on the complete genomes of these two strains we could not detect any change in the genes responsible for LOS (lipooligopolysaccharide) synthesis that would promote formation of smooth colonies (Fig 4b; note, in 1218S *lsr2* is duplicated, Fig 4a; see also Fig S5a, b). We therefore analyzed the transcript levels of the LOS genes. Also, we analyzed the mRNA levels for the global regulator Lsr2 and the transcription factor WhiB4, which both have been implicated to act as regulators of LOS genes. Mutations in several of these genes have been reported to affect LOS synthesis and rough colony morphotype (Fig 4b, marked with red arrows)^see^ ^e.g. 21,29,42,46,47^. The LOS genes are transcribed in both 1218R and 1218S and the levels of the individual genes vary. The distribution pattern appears to be similar both in exponential growing and stationary 1218R and 1218S cells (Fig S9a, c).

For 1218R, comparing the mRNA levels in exponential and stationary phase suggest that the majority of the LOS gene transcripts are higher in exponentially growing cells than in stationary cells (Fig S9b; see also Table S5a). This also applies to the *lsr2* transcript (Lsr2, suggested to repress LOS genes; for references see above), while the *whiB*4 transcript level increases in stationary cells (see also Table S5). For 1218S, the mRNA levels for many of the LOS genes are higher in exponential 1218S cells but we detected increased levels for some in stationary cells (Fig S9d; Table S5), *e.g*. the SL659 acyltransferase *papA*1 gene, *papA*1_2, and its inactivation result in a rough phenotype while a mutation in *pimF* (*losA*) gives a smooth/wrinkled phenotype^29^.

Comparing “LOS” mRNA levels in 1218R *vs* 1218S in exponential growing and stationary cells revealed that the levels for the majority of the LOS genes are higher in exponential 1218R cells. Conversely, higher transcript levels were detected in stationary 1218S cells relative to 1218R (Fig 6; Table S5b).

**Figure 6.**
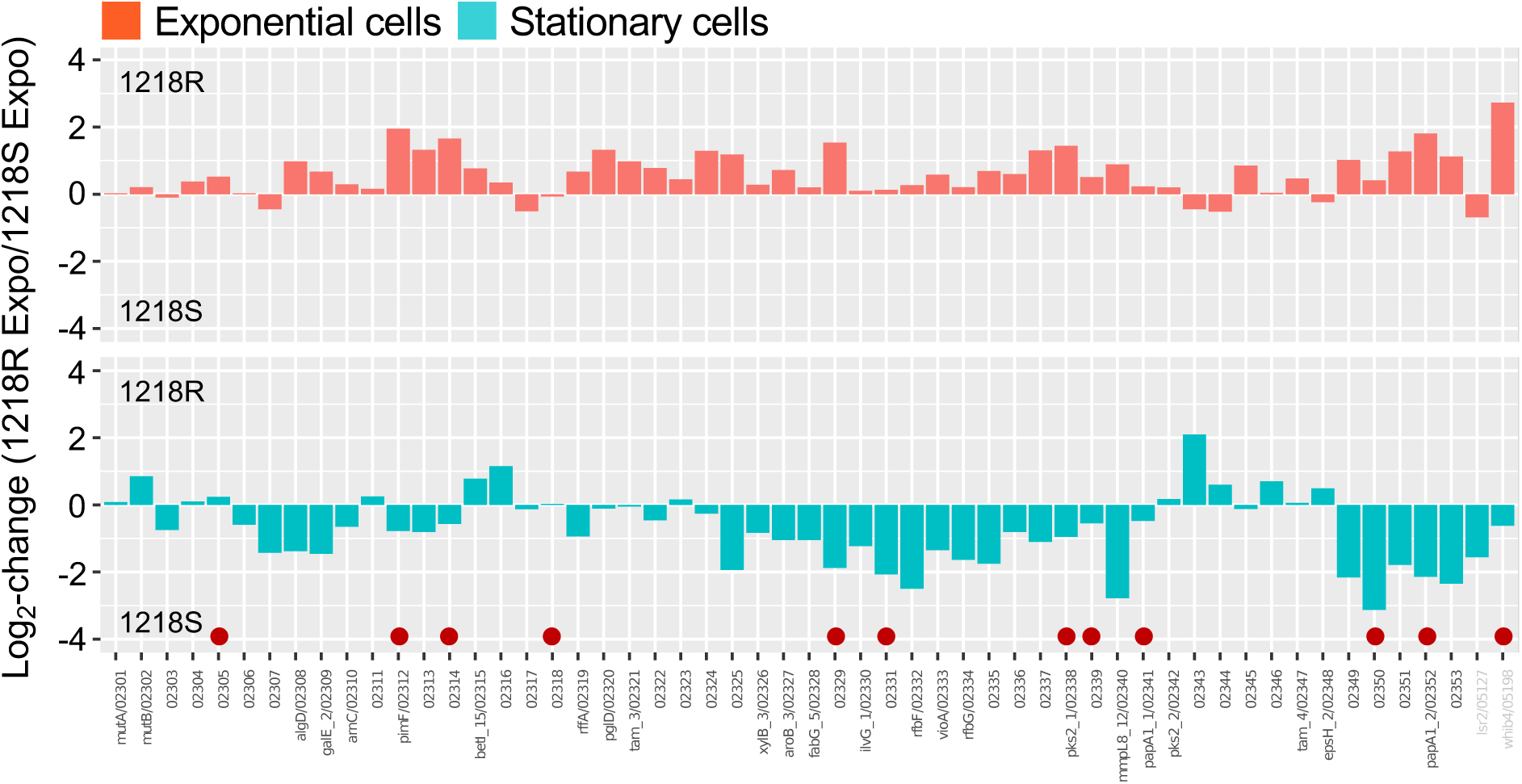
Transcript levels of LOS related genes in 1218R and 1218S. Log_2_-fold change comparing transcript levels in exponentially growing (in red) and stationary cells (in turquoise) 1218R and 1218S. Levels with negative values correspond to higher levels in 1218S while positive values refer to higher levels in 1218R. Naming of the genes refer to 1218R. The *lsr2* and *whiB*4 genes are marked in grey and the red dots mark genes, when mutated result in formation of rough colonies in *Mmar*^M^, see main text for details. For the distribution and log_2_-fold change in 1218R and 1218S, respectively, see Figure S9a-d.

For Lsr2, the mRNA levels are higher both in 1218R and 1218S exponential cells relative to stationary cells. Comparing 1218R vs. 1218S indicate that the Lsr2 mRNA level is higher in 1218S, in particular in stationary cells. This difference might be attributed to the duplication of *lsr2* in 1218S (Fig 6; Tables S4a and S5a). But, note that we could not determine from which copy the transcripts orginated in 1218S because the two copies are identical (see above).

With respect to WhiB4, the mRNA levels increase in stationary cells in both 1218R and 1218S. The level is, however, higher in exponentially growing 1218R than in 1218S cells while the level is marginally higher in stationary 1218S cells (Fig 6; Table S5).

Taken together, conceivably the higher levels of some of the LOS gene transcripts in stationary 1218S cells result in increased protein levels that subsequently influence LOS synthesis (see also the discussion). However, this analysis did not give a clear answer that rationalize the difference in 1218R and 1218S colony morphotypes suggesting that additional factors influence the formation of rough and smooth colonies. Considering the *Mmar*^CCUG^ and *Mmar*^M^ we detected roughly similar responses with some differences, as in the 1218R strain, when comparing LOS mRNA levels in exponentially growing and stationary cells. Relative to 1218R, however, we note several differences (for details see Table S5).

### Non-coding RNA and LOS gene mRNA levels

Non-coding RNAs have been demonstrated to have key roles in the regulation of gene expression^see^ ^e.g. 48,49^. Available data suggest that Ms1 RNA act as a global regulator in mycobacteria with a role in adaptation to growth in stationary phase and its level increase in stationary cells^50–52^. Hence, we analyzed the levels of Ms1 RNA in 1218R, 1218S, *Mmar*^CCUG^ and *Mmar*^M^ to understand whether the Ms1 RNA level correlates with the levels of LOS gene transcripts.

The Ms1 RNA levels increased in stationary phase relative to exponential growing *Mmar* cells irrespective of strain (Figs 7a and S10; Table S4). This in keeping with previous data and its proposed role during the transition from exponential to stationary phase^50–52^. When we compared the Ms1 RNA levels in the different strains we detected higher levels in 1218S, *Mmar*^CCUG^ and *Mmar*^M^ relative to 1218R in both exponentially growing and in stationary cells (Fig 7b; Table S4). Following this we cloned the Ms1 RNA gene behind the tetracycline inducible promoter TetRO in pBS401 (see Materials and Methods) generating pBS401^Ms1RNA^. This construct was introduced into *Mmar*^CCUG^ to investigate which genes are affected in response to overexpressing Ms1 RNA. As expected, Ms1 RNA increased upon introducing pBS401^Ms1RNA^ and addition of tetracycline into the media (Fig 7c; Table S6). Monitoring the impact of Ms1 RNA on the transcription of the LOS region revealed that several LOS gene transcripts increase in *Mmar*^CCUG^. For example, *pimF*/*losA* (glycosyltransferase) and *papA*1_2 (SL659 acyltransferase), *rffA* (dTDP-4-amino-4,6-dideoxygalactose transaminase) and *pglD*/*wecE* (UDP-N-acetylbacillosamine N-acetyltransferase) (Fig 7c, see also Table S6). Mutations in these genes in *Mmar*^M^ result in a rough phenotype and/or affect LOS synthesis (marked with red dots in Fig 7c)^see^ ^e.g. 28^. We also detected a modest increase in the *lsr2* transcript and higher *whiB*4 transcript levels in response to overexpressing Ms1 RNA (Tables S4 and S5).

**Figure 7.**
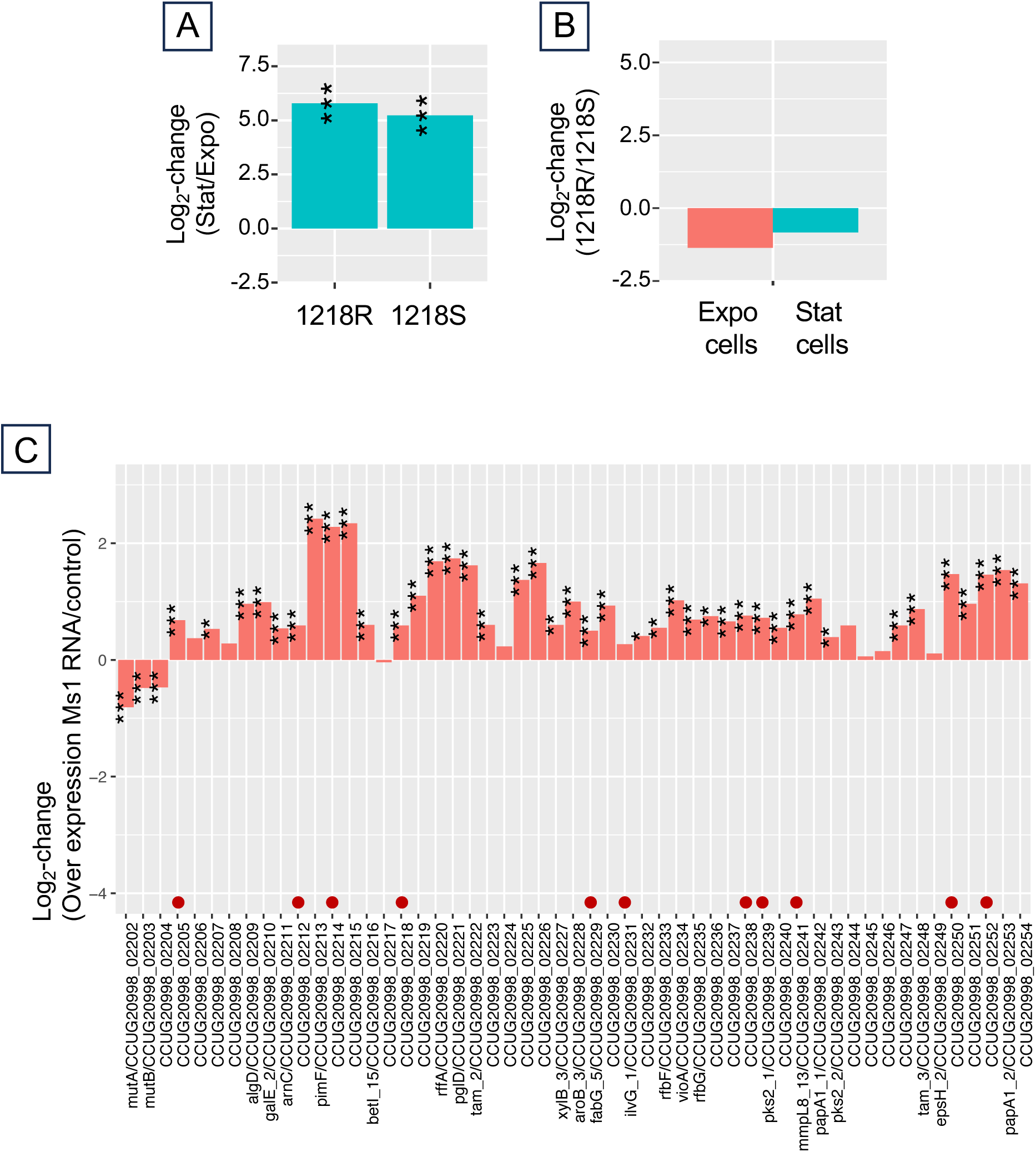
Ms1 RNA and levels of LOS gene transcripts. (a) Ms1 RNA levels in exponentially growing and stationary 1218R and 1218S cells expressed as log_2_-fold change. For Ms1 RNA levels in *Mmar*^CCUG^ and *Mmar*^M^, see Figure S10. (b) Log_2_-fold change comparing transcript levels in exponentially growing (in red) and stationary cells (in turquoise) 1218R and 1218S. Levels with negative values correspond to higher levels in 1218S while positive values refer to higher levels in 1218R. (c) Change in LOS gene transcripts in response to overexpression of Ms1 RNA in exponentially growing *Mmar*^CCUG^ cells expressed as log_2_-fold change. Statistical significance, see Materials and Methods; *p-value < 0.05; **p-value < 0.01; ***p- value < 0.001.

Taken together, these data suggest that Ms1 RNA influence the levels of LOS gene transcripts albeit we cannot distinguish if this is direct or indirect via, *e.g*. WhiB4. Nevertheless, given that the level of Ms1 RNA is higher in 1218S than in 1218R (Table S5b) might be one reason to that LOS gene transcripts are more abundant in 1218S in stationary cells compared to 1218R. In addition, several of the ESX-1 gene transcripts, including *esxB*_3 and *esxA*_4, increased indicating a role of Ms1 RNA also in the expression of ESX-1 genes, while the mRNA levels for the majority of *Mmar*^CCUG^ ESX-6 genes were slightly reduced (if any) upon overexpression of Ms1 RNA (Table S5c).

### Biofilm formation in 1218R and 1218S

A previous report suggested that deleting the ESX-1 genes *espE*, *espF*, *espG* and *espH* in a clinical *Mmar* isolate attenuated the generation of biofilm and sliding motility^32^. We therefore decided to study biofilm formation in 1218R and 1218S cultures to understand if biofilm formation differs comparing these two *Mmar* strains. In keeping with that ESX-1 genes influence the generation of biofilm, 1218R forms a more robust biofilm relative to 1218S, which would be in agreement with that the missing genes “*espF*_2 – *esxB*_3” influence biofilm formation (not shown, but see Fig 8). Hence, we cloned the *espF*_2*-espH* region from 1218R in the plasmid pBS401 generating pBS401*^espF-espH^* (see Materials and Methods), to understand whether these genes contribute to biofilm formation.

**Figure 8.**
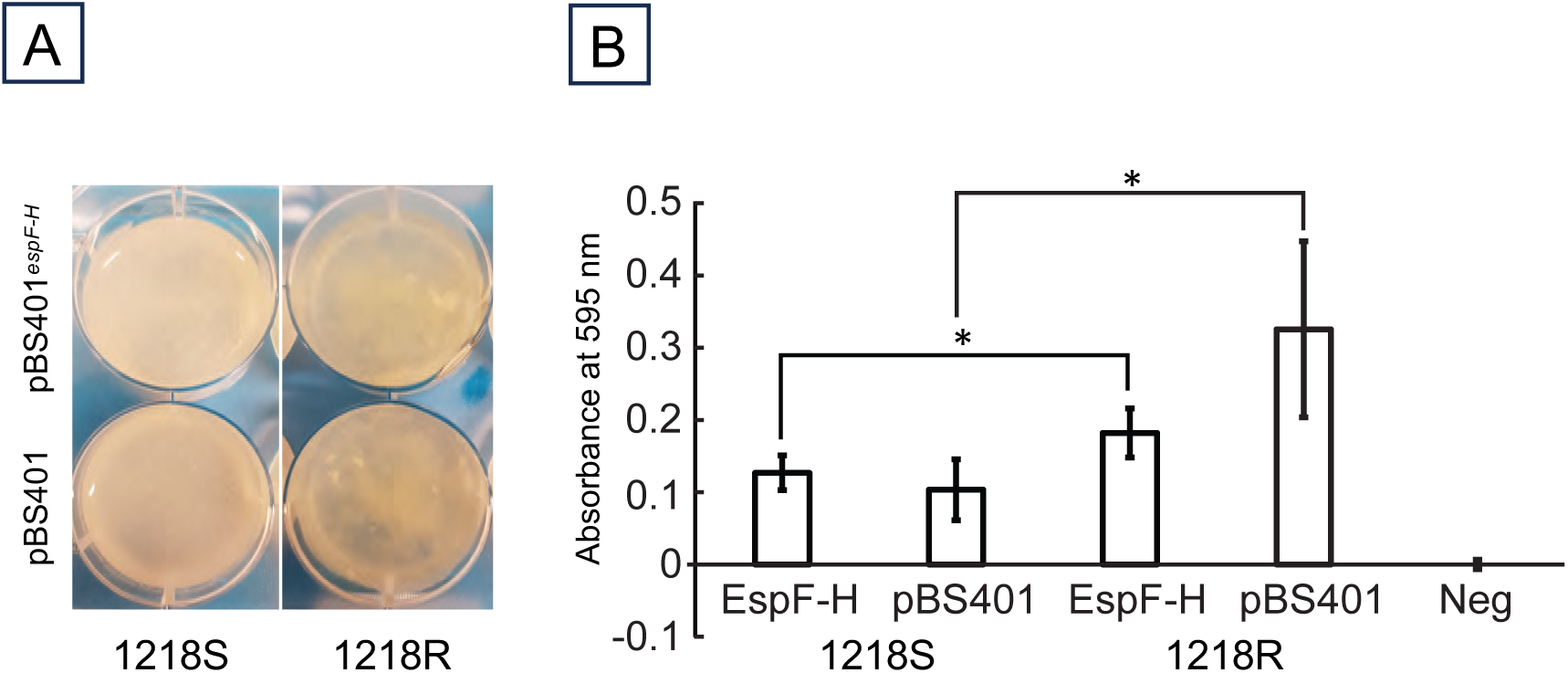
Biofilm formation. The 1218R and 1218S *Mmar* strains were transformed with the empty control plasmid (pBS401) or with pBS401*^espF-H^*and cultivated as outlined in Materials and Methods. (a) Visualization of biofilm formation in 6-well micro titer plates after 26 days of growth at 30°C in 7H9 media without Tween 80 and Hygromycin B. (b) Quantification of biofilm formation after cultivation as in (a) in 96-well micro titer plates as in (a) according Syal^100^, for details see Materials and Methods. Average values of the four measurements, two biological replicates and two technical replicates per biological replicate, with standard deviations are shown and significant differences are indicated with connecting horizontal square brackets where * indicates a p-value < 0.05).

Introduction of the vector pBS401 into 1218R and 1218S verified our initial observation that 1218R produces more biofilm than 1218S (Fig 8; see also Fig S11). The presence of pBS401*^espF-espH^* did not change biofilm formation in 1218S cultures while a reduction was detected for the 1218R strain. These data indicate that an increased number of the ESX-1 genes, *espF*_2, *espG*1_2 and *espH*, influence biofilm formation negatively in 1218R. But, their presence on pBS401*^espF-espH^* did not complement the 1218S biofilm formation phenotype. Therefore, the difference in biofilm formation comparing these two strains is not simply related to the deletion of *espF*_2*-espH*.

## Discussion

As for other bacteria the composition and structure of the mycobacterial cell wall play a critical role for the interaction with the environment and the immune system. In this context, systems involved in secretion such as the type VII secretion systems are essential. The *Mmar* 1218S strain was originally isolated as a smooth variant of 1218R, which forms rough colonies and is more virulent than 1218S. Based on the 1218S draft genome we reported that it lacks part of the ESX-1 secretion system (for reference cf. Table 1 in Ref 18). The complete 1218S genome sequence confirms this but showed that part of the ESX-1 *esxB*_3 gene is still present. With respect to the 1218S draft genome this is most likely due to that *esxB*_3 is positioned close to a scaffold that affected the assembly of the draft genome. Comparing five *Mmar* strains for which the complete genomes are available revealed that the ESX-1 regions in 1218R, *Mmar*^CCUG^, *Mmar*^ATCC^ and *Mmar*^M^ are very similar. Extending this comparison by including other *M. ulcerans* clade members, *M. ulcerans* Agy99, *M. liflandii* 128FXT and *M. pseudoshottsii*^35,53–55^ show that the majority of the ESX-1 genes are absent in the two former mycobacteria, but present in the latter. Moreover, ESX-1 is partly duplicated in *M. marinum* and is refered to as ESX-6^7^. ESX-6 is missing in *M. ulcerans* Agy99, *M. liflandii* 128FXT and *M. pseudoshottsii* (note, however, that some of the ESX-6 genes are present in the two latter mycobacteria^35,53–55^. As will be discussed below, we also note variations in the presence of LOS genes in members of *M. ulcerans* clade. Taken together, this is in agreement with that *M. ulcerans* clade members are exposed to evolutionary pressure and suggest that the ESX-1 and LOS (see below) regions include genes that are exposed to changes^18,35,56^.

### Consequences of the deletion of “espF_2 – esxB_3” on ESX-1 and ESX-6 transcription

The “*espF*_2 – *esxB*_3” region in the ESX-1 locus is missing in 1218S^18^, this study. The complete 1218S genome, however, revealed that parts of the *espF*_2 “N-terminal sequences” and *esxB*_3 “C-terminal sequences” are still present. The RNA-Seq data suggested that the resulting truncated region is transcribed (see below). Among the deleted genes *eccA*_1_, *eccB*_1_, *eccCa*_1_ and *eccCb*_1_ encode for proteins that are part of the ESX-1 secretion machinery and mutations in these genes abolish/reduce ESX-1 specific secretion^16,17^. Furthermore, the “*espF*_2 – *esxB*_3” region encompasses (following 1218R gene annotation): *espF*_2 and *esxB*_3, encode for proteins that are secreted using the ESX-1 machinary, *espG*1_2 encodes for a chaperone involved in secretion of PE/PPE substrates by delivering these substrates to EccA, and EspH, which is needed for secretion of EspF and also EspE (*espE* is still present in 1218S). Mutating or deleting these genes influence the virulence of *M. marinum* as well as *M. tuberculosis*^16,57,58^. Hence, absence of these gene (“Δ*espF*_2 – *esxB*_3”) products in 1218S is one plausible reason to the roughly four-fold reduction in virulence comparing 1218R and 1218S infection of Japanese medaka^18^.

The “Δ*espF*_2 – *esxB*_3” deletion in 1218S begin at position 6408294 and ends at position 6418981 (1218R numbering; Fig S2a). Hence, codons for 21 amino acids in the EspF N-terminus are still present while codons corresponding to 36 amino acids of the “EsxB_3 N-terminus” are missing in 1218S. But, the remaining part of the “EsxB_3 mRNA” is out of frame with respect to the native EsxB_3 translational stop codon, UGA (Figs 3b and S2a). Recently it was suggested that the ESX-1 genes are transcribed as one unit starting 120 bps upstream of *espM* in the *Mmar* strain NTUH-M6885^32^. This is in contrast to *Mtb* where several transcriptional units were identified^59^ and the sequences near the transcription start sites upstream of *espE* and *espF*_2 are conserved comparing *Mmar* and *Mtb* H37Rv. Hence, it is not excluded that genes within the ESX-1 region in *M. marinum* are transcribed as separate units, which would be similar to *Mtb* H37Rv. Also, in the *Msmeg* ESX-1 locus promoters have been identified, one upstream of the PE35 gene and one between PPE68 and *esxB*^60^ supporting transcription of separate units also in *Mmar*. Nevertheless, our RNA-Seq data suggested that in 1218S *espM*, parts of *espF*_2 and *esxB*_3, and *esxA*_3 might be transcribed together [Fig S8; the 1218R RNA-Seq data suggest that this region might also be transcribed together (not shown)]. We speculate that the Shine-Dalgarno sequence upstream of *espF*_2 binds to the ribosome and initiating translation, which would generate a 98 amino acids long polypeptide. The amino acid sequence of this polypeptide would be different compared to wild-type EsxB_3 and encompasses 21 EspF amino acids followed by amino acids originating from the truncated *esxB*_3 gene. Translation terminates when the ribosome encounters a UGA stop codon, which overlaps the EsxA_3 start codon AUG (see Fig 3b). Presence of overlapping “…UG<u>A</u>UG…”- regions among mycobacteria is supported by mass spectrometry data of *Mtb* H37Rv peptides, representing the N-terminus of Rv2537c and Rv1295, which indicated that translational start sites overlap with the stop codon of the annotated upstream gene (see Supplementary material)^61^. Also, our unpublished peptide analysis of mass spectrometry *Mmar*^CCUG^ data (to be publish elsewhere) revealed overlapping translational start (*pspA*_3; gene id, CCUG20998_03005) and termination sites (*uppP*; gene id, CCUG20998_03006). The altered “EsxB_3” peptide is unlikely to interact with EsxA to form the heterodimer that is recognized by EccCb^116,58,62^. This is supported based on structural prediction with AlphaFold3^63^ suggesting that the structure of the altered “EsxB_3” peptide is significantly different compared to the native EsxB_3 protein (not shown). If the altered “EsxB_3” peptide is expressed and functional, does it affect the physiology of the cell, and if so, how? This warrants for further studies. We can not exclude, however, the possibility that an internal *esxB*_3 GUG codon is used to initiate translation, which then would terminate at the normal UGA stop codon. The resulting EsxB_3 peptide would be truncated but it will still have the native C-terminus, which has a role in the interaction with EccCb_1_ after formation of the “EsxB_3-EsxA_3” complex^16,58,62^. Either way, given that EccCb_1_ and the ESX-1 secretion machinery is missing in 1218S raises the question whether EsxA_3 interact with the ESX-6 EsxB_1 (or EsxB_2) protein (see below) and is secreted using another ESX secretion pathway.

In *M. marinum* ESX-6 is a partial duplication of ESX-1^7^ and our RNA-Seq data show that the ESX-6 gene transcripts were significantly increased in the strain lacking ESX-1 secretion machinery genes (1218S). For example, in 1218S the ESX-6 *esxB*_1, *esxA*_1 and *esxA*_2 gene transcripts were among the most abundant relative to other ESX (−1, −3, −4 and −5) gene transcripts. As for ESX-1 gene transcripts in 1218R, the ESX-6 gene transcritps increased in stationary 1218S cells. Furthermore, for 1218R (also *Mmar*^CCUG^ and *Mmar*^M^), the ESX-6 transcript levels were low in both exponentially growing and stationary cells, which is in contrast to 1218S in which the levels are significantly higher relative to 1218R in stationary as well as in exponentially growing cells. These findings are in agreement with a previous report, which suggested that ESX-6 *esxB*_1 and *esxA*_1 in *Mmar*^M^ is expressed albeit at a low level^64^. In this study, the transcript was detected but the ESX-6 EsxB protein was too low to be detected by Western blot analysis.

The sequence of the duplicated ESX-1 region in ESX-6 is conserved^7^, in particular pertaining to *esxB*_1 and *esxA*_1, which are almost identical [except that in 1218S *esxB*_3 (ESX-1) is truncated (Figs 3 and S3a)]. In 1218R, the presumed ESX-1 *esxB*_3 Shine-Dalgarno sequence (SD; 5’AGAAAG) is located 9 bp upstream of the translational start codon AUG (Fig S3b). In relation to ESX-6, and as reported previously for *Mmar*^M 64^, *esxB*_1 has an insert of 14 bp in the 5’UTR (<u>u</u>ntranslated <u>r</u>egion), which appears to relocate the SD sequence and extend the distance to the translation start codon, AUG. Or alternatively, reducing the number of SD-residues to 5’GAG (located 6 residues upstream of the AUG start codon) resulting in a weak SD sequence (Fig S3b). Regardless, this might therefore be part of the reason to why the EsxB_1 protein was produced at a low level also in *Mmar*^M^ lacking the ESX-1 *esxB*_3 gene in spite that the ESX-6 *esxB*_1 gene was transcribed^62^. For 1218S, perhaps the higher transcript levels of ESX-6 genes result in higher protein levels, *e.g*. EsxB_1. However, whether the higher levels of *esxB*_1, *esxA*_1 and *esxA*_2 transcripts also lead to increased protein levels remains to be investigated.

Apart from that no ESX-1 mRNA for the deleted genes in 1218S were detected (but see above), transcripts for ESX-1 substrates, *e.g*. *esxA*_3, and associated ESX-1 genes *espACD* were detected in 1218S, both in exponential growing and stationary cells, albeit the levels are lower than in 1218R. This also pertains to the regulatory genes *whiB*6 and *espM*, which both are involved in the regulation of the transcription of ESX-1 genes^15,16,58^. Our findings are in keeping with previous published data where disruption of genes in the “*espF* – *esxB*_3” region affect transcription of *whiB*6 and *espM* and that there is a link between the status of the ESX-1 secretion machinery and expression of ESX-1 substrates^16,44,58^. Here we note that besides *whiB*6 and *espM*, Chirakos *et al*.^16,58^ discussed the involvement of additional transcription factors (to be identified, but see below) in the regulation of ESX-1 genes as well as that posttranscriptional regulation play’s a role. Also, PhoP is involved in the regulation of *whiB*6, *espACD* and *espR*, where the transcription factor EspR regulates *espACD* expression^38^. For PhoRP, we did not detect any significant difference in the mRNA levels comparing 1218R and 1218S, while the EspR mRNA level is higher in both 1218R and 1218S stationary cells than in exponential cells (Fig S7a, b; Table S4), and the level was modestly higher in exponential 1218R cells compared to 1218S exponential cells (Fig 5; Table S4). Moreover, overexpression of the non-coding RNA Ms1 RNA in *Mmar*^CCUG^ increased “ESX-1 mRNA” levels, but not “ESX-6 mRNAs”, in keeping with that ESX-1 transcripts increased in stationary *M. marinum* cells. Our data also indicated that the WhiB6 and EspM mRNA levels were not affected to any significant extent in response to increasing Ms1 RNA. Together these findings add to the complexity of the regulation of the transcription of the ESX-1 (and associated) genes and the involvement of the non-coding RNA Ms1 RNA (see also below).

Transcripts are subjected to processing and degradation and show variation in their stabilities. Different ribonucleases and RNA helicases participates in these processes^see^ ^e.g^ ^65–68^. Therefore, the higher levels of *esxB*_3 and *esxA*_3 (ESX-1) [and *esxB*_1 and *esxA*_1 (ESX-6) transcripts] might be related to their stabilities due to the interplay between mRNA degradation and processing. We presented data showing that part of the middle of the RNase J gene is deleted in 1218S (Figs S3a; Table S1a, b), which might affect its function and thereby RNA processing and/or degradation. Absence of *M. tuberculosis* RNase J resulted in accumulation of structured mRNA fragments^69^. It is therefore conceivable that the RNA processing and/or degradation differs in 1218S relative to 1218R. In this context, we also note that in *Escherichia coli* the endoribonuclease P (RNase P) is involved in the processing of tRNA as well as mRNA^70–71^. Our unpublised data suggest that higher levels of the RNase P RNA component in *Mmar*^CCUG^ influence the levels ESX-1 gene (and associated gene) transcripts, which might be related to the processing/degradation of these transcripts.

Among the deleted ESX-1 genes in 1218S *espF*_2, *espG*_1__2 and *espH* encode for proteins that are: secreted (EspF), functioning as a chaperone (EspG) and needed for secretion of EspF and EspE (EspH)^16,57,58,72^ and refs therein. Recently, it was reported that EspE, EspF, EspG and EspH affect sliding motility and biofilm formation in the *M. marinum* strain NTUH-M6885^32^. Our data indicated that 1218R forms more biofilm compared to 1218S consisting with these data. Expressing *espF*, *espG*_1_ and *espD* carried on a plasmid resulted in no effect on biofilm formation in the case of 1218S, while a modest reduction was detected in 1218R cultures. Together this suggests that *espF*, *espG*_1_ and *espH* have a modest role in biofilm formation in *M. marinum* cultures. In this context, studies of aerosol *M. tuberculosis* infected animals suggested that the persistent lesion necrosis in partially calcified primary lesions, and that a cellular layer that lines the interior surface of cavitary lesions, resembles the structure of biofilms^73^. Thus, the capacity to form more robust biofilms might also be part of the reason why 1218R showed a higher virulence than 1218S in infecting Japanese medaka^18^.

To conclude, to understand the regulation and expression of ESX-1 vs. ESX-6 genes warrants for further analysis but our current findings pave the way for future experiments.

### LOS and related genes and colony morphology

In the past the tubercle bacillus cell morphology was studied and many studies report variation in colony morphology such as formation of rough and smooth colonies. Early chemical studies of *Mycobacterium bovis* BCG indicated that the difference in cell morphology appeared to be related to variation in lipid content^74^. Also, *Mtb* strains showing colony morphology variations influenced virulence^see^ ^e.g. 75^ and refs therein. More recent studies suggested that the TB complex member *Mycobacterium canetti*, which forms smooth colonies, is less virulent than *Mtb*^22,76,77^ and refs therein. Other mycobacteria have also been reported to form rough and smooth colonies. For instance, *Mycobacterium kansasii* strains showing rough colony morphotype are more virulent^78^. As reported elsewhere and in this study, *Mmar* also forms rough and smooth colonies where the rough phenotype is associated with higher virulence^18,29,30^. For *M. kansasii* and *M. canetti* and in particular *Mmar*, recent genetic and chemical studies have suggested that variation in the amount of lipooligosaccharide (LOS) on the surface of these mycobacteria affect the colony morphotype^26,29,79–81^. Comparative genomic analysis revealed that several LOS genes are present in *M. kansasii*, *M. canetti* and *Mmar* while several of these are missing in *Mtb* H37Rv^23,77^. Albeit LOS genes are mainly present among slow growing mycobacteria, LOS genes are also present in some rapid growing mycobacteria such as *Mycobacterium smegmatis*^21,82^ (note that smooth colony mycobacteria variants are also linked to the presence of glycopeptidolipids (C-type) on the cell surface^24^). In *Mmar* the majority of the LOS genes (approx. 40 genes) are located in the LOS region (Fig 4). Mutations in several of the genes affect LOS synthesis and as a conseqeunce results in rough colony morphotypes (for Refs see above). *M. marinum* belongs to the *Mycobacterium ulcerans* clade^35^ and LOS genes including the two polyketide synthase genes (*pks2*_1/*pks5* and *pks2*_2/*pks5.1*), the polyketide synthase associated protein gene (*papA1*_1/*papA4*) and the transmembrane protein MmpL8_12 are not present in *M. ulcerans* Agy99 (not shown)^7^. Other *M. ulcerans* clade members (*Mycobacterium liflandii* and *Mycobacterium pseudoshottsii*) also lack genes in the LOS-region but one of the *pks2* genes is still present in these two species^35,54,55^. Absence of LOS genes is in keeping with that these three mycobacteria form rough colonies^83,84^. Together these data suggest that the LOS region (as the ESX-1 region; see above) is subjected to evolutionary pressure, which would be expected given the location of LOS on the cell surface. Interestingly, a transposase gene is annotated in this region in *M. ulcerans* Agy99^7^, which might have played a role in the deletion of LOS genes. We detected no difference in the sequence of the LOS region in 1218R and 1218S comparing their complete genomes that could explain the difference in colony morphotypes. However, our RNA-Seq data of RNA isolated from exponentially growing and stationary 1218R and 1218S cells suggest differences in the transcription levels of the LOS genes. It is plausible that this difference, where the majority of the of LOS gene transcripts appeared to be higher in stationary 1218S cells compared to 1218R (Fig 6), is part of the reason to the different colony morphologies. However, the levels for the regulators Lsr2 and WhiB4 were not in keeping with this difference (Figs 6 and S9a-d; Table S4). Lsr2 has been reported to act mainly as a repressor of LOS genes and deletion of *lsr2* in *Mycobacterium smegmatis* result in a rough colony morphotype^47,85^. *Mmar* 1218S has two identical *lsr2* genes and we note that the mucoid *Mycobacterium mucogenicum* also has two genes annotated as *lsr2*^20^. Thus, higher Lsr2 levels might result in promoting smooth or mucoid colonies. Currently, we do not have an explanation to why the *lsr2* transcript level is higher (apart from that *lsr2* is duplicated in 1218S) in 1218S than in 1218R in relation to that 1218S forms smooth colonies. This requires further and deeper analysis *e.g*., does the higher Lsr2 mRNA levels lead to an increase in Lsr2 protein (see also below). In this context, however, our data suggested that additional genes/factors might play a role in changing the *Mmar* colony morphotype. Hence, one reason to the higher levels of LOS gene transcripts in 1218S relative to 1218R stationary cells might be related to, at least partly, the mutated RNase J gene in 1218S. RNase J is, as discussed above, involved in mRNA processing/degradation and an altered RNase J might influence this process thereby affecting mRNA stability (and/or processing) in 1218S vs. 1218R. This might also relate to RNase P, see below.

Another player to consider is the protein kinase PknB, which phosphorylates Lsr2 and thereby regulates its activity^86^. Recent studies suggest that the protein kinase PknL and a protein similar to PKSS (MSMEG_4727; involved in regulating lipid components of the outer cell envelope) in *M. smegmatis* also affect the activity of Lsr2^46^. This raises the question whether phosphorylation of Lsr2 differs in 1218R vs. 1218S and thereby is one factor to why 1218S forms smooth colonies in spite higher levels of *lsr2* transcripts. Furthermore, Nguyen *et al*.^87^ also showed that a FbpA (involved in generating the mycobacterial cord factor, α, α’-trehalose dimycolate) *M. smegmatis* mutant displayed a smooth phenotype indicating a role of this protein. However, there is no variation in FbpA transcript levels comparing 1218R and 1218S (Table S5a, b).

Following the discussion above, an additional candidate involved in regulating the expression of stationary genes is the global regulator Ms1 RNA. Ms1 RNA is highly abundant in stationary phase and binds to RNA polymerase and influence gene transcription^50–52^, this study. Overexpression of Ms1 RNA in *Mmar*^CCUG^ increase transcripts of several LOS genes, *e.g*. *pimF*, *pglD* and *papA*1_2, and mutations in these leads to loss of LOS and changed colony morphologies^29^. Deleting the Ms1 RNA gene in *M. smegmatis* did not change the levels of LOS gene transcripts in exponential or in stationary phase^52^. Therefore, the impact of Ms1 RNA on transcription appears to differ comparing the SGM *Mmar* and the RGM *M. smegmatis*. In this context, our unpublished data suggest that overexpression of the RNA component of the RNase P also affects the mRNA levels of several LOS genes (see also above). Hence, the roles of these players as well as other sRNAs (to be identified) with respect to LOS gene regulation need to be studied. For example, recent data suggest that 6C RNA is involved in the regulation of LOS genes^33^. Finally, given that components of the ESX-1 secretion machinery are missing in 1218S might influence secretion of factors that relate to the formation of LOS and thereby indirectly link LOS and ESX-1, and/or ESX-6.

To conclude, we are just at the doorstep to understand the regulation of LOS genes. Hence, this warrants for further studies to understand their regulation and to identify missing factors that affect LOS synthesis and mycobacterial colony morphotypes. This knowledge would eventually be important and useful to treat mycobacterial infections.

## Materials and Methods

### Strains, cultivation, DNA isolation, DNA sequencing and assembly

Information about and cultivation of the *Mmar* strains 1218R, 1218S, *Mmar*^CCUG^ and *Mmar*^M^ have been described elsewhere^18,41^. Isolation and preparation of 1218S genomic DNA according to Das et al.^18^. The DNA was sequenced using the Pacific Biosciences (PacBio) platform at the NGI-Uppsala Genome Center at Uppsala University. Sequencing reads with an average length of 11686 bp and coverage approximately 118 were assembled with the SMRT-analysis HGAP3 assembly pipeline (Chin) and polished using Quiver (Pacific Biosciences, Menlo Park, CA, USA) generated a single scaffold representing the complete 1218S genome. The complete genomes for 1218R, *Mmar*^CCUG^, *Mmar*^M^ and *Mmar*^ATCC927^ have been reported elsewhere^7,18,55^.

### Genome annotation, functional classification, and identification and analysis of core genes

The PROKKA software (version 1.0.9)^88^ was used for gene annotation and prediction of coding sequences (CDS). For functional classification, the predicted (PROKKA and NCBI annotated) CDS were subjected to BLASTp against the RAST predicted CDS followed by mapping the RAST subsystem database (http://rast.nmpdr.org/, last accessed April 15, 2020)^89^ using the BLAST approach.

Identification of core genes were performed as described elsewhere^20,51^. To identify core and unique genes, predicted CDS from the genomes were used for “all-*vs*-all” BLAST search. Based on the BLAST results orthologous genes were identified using PanOCT with minimum 45% identity and 65% query coverage^90,see^ ^also^ ^18,91,92^.

### Prediction of virulence factor genes (VFDB)

We used the VFanalyzer tool, available at the Virulence Factor Data Base (VFDB)^93,94^, to predict virulence factor genes. As input to the VFanalyzer we initially we used protein CDS and *Mtb*^H37Rv^, *Msmeg*^MC2-155^, *Mmar*^M^, *M. ulcerans* strain Agy99, and *M. avium paratuberculosis* strain K10 as references (available at the VFDB database). The single XLS data files were combined and the resulting data files were used to present the data using the R interface (ggplot2 package)^95^.

### RNA extraction, RNA sequencing and analysis

The RNA-Seq analysis were done as reported previously^20,41^. Briefly, 1218R, 1218S, *Mmar*^CCUG^ and *Mmar*^M^ were grown in 7H9 media at 30°C. Cells (biological duplicates) were harvested in exponential and stationary growth phase. Total RNA was extracted with Trizol using a bead beater, treated with DNase and submitted for RNA sequencing at the SNP@SEQ Technology Platform at Uppsala University (HiSeq 2000 Illumina platform).

The RNA-Seq data sets (number of reads) were mapped to the complete genomes by building the index with bowtie2 v2-2-4^96^ and alignment using the Tophat v2.0.13 tool^97^. Read-counts were generated from the aligned BAM files with the HTseq v0.9.1^98^. For normalization and differential analysis, we used the Deseq.2 package, which gives p + adj values (statistic significance)^99^. With respect to the RNA-Seq data for, 1218R and 1218S see Pettersson et al.^41^ and for *Mmar*^CCUG^ see Pettersson et al.^20,41^.

### Biofilm analysis

The *espF*_2, *espG*_1__2 and *espH* genes from 1218R were cloned into the pBS401 plasmid behind an anhydrotetracycline inducible promoter^100^ and screened using plasmid specific primers (see Supplementary information). This construct is refered to as pBS401*^espF-H^*. The *Mmar* 1218R and 1218S strains were transformed with an empty control plasmid (pBS401) or with pBS401*^espF-H^*. The cells were cultivated in microtiter plates for 26 days in 7H9 medium without Tween 80 or hygromycin B. The cultures grown in the 96-well plates were used for the quantification of biofilm formation^101^. Each strain and plasmid combination were measured in two biological replicates, with two technical replicates per biological replicate; in total four measurements per combination.

## Ethics Statement

All methods were carried out in accordance with relevant guidelines and regulations.

## Acknowledgements

In memory, this paper is dedicated to Dr D. G. Ennis University of Louisiana, Lafayette, USA, a friend and great scientist. We thank our colleagues for discussions. RNA-Seq was performed by the SNP&SEQ Technology Platform in Uppsala, which is part of the Science for Life Laboratory at Uppsala University and supported as a national infrastructure by the Swedish Research Council. The computations were performed on resources provided by SNIC through Uppsala Multidisciplinary Center for Advanced Computational Science (UPPMAX) under Project b2011072. This work was funded by the Swedish Research Council (M and N/T), the Swedish Research Council for Environment, Agricultural Sciences, and Spatial Planning (FORMAS), and Uppsala RNA Research Center (Swedish Research Council Linneus support) to LAK.

## Authors contributions

LAK conceived the study. PRKB performed the bioinformatics computations. MR and BMFP cultivated, prepared DNA and RNA from the different *Mmar* strains. BMFP performed the biofilm experiments. PRKB, BMFP, and LAK analyzed and interpreted the data. LAK wrote the manuscript and all authors read and approved the final version of the manuscript.

## Data availability

The complete genome sequence of *Mycobacterium marinum* strain DE4381 (1218S) is available at NCBI under BioProject ID PRJNA1428566, with the GenBank accession number JBVOUM000000000. The complete transcriptomic datasets used in this study are available at NCBI under BioProject ID PRJNA1423637. Additional information on genome annotation or datasets is available from the corresponding author upon reasonable request.

Supplementary material available at: https://drive.google.com/drive/folders/1SVfrb_wvYiBP6CehuLgXznmQ0H3OVfx1?usp=sharing

