## Supplementary material for "Loss of *Mycobacterium marinum* ESX-1 genes increase transcription of ESX-6 genes": SUPPL TEXT AND FIGS S1-11

[1]Department of Cell and Molecular Biology
Box 596, Biomedical Centre
SE-751 24 Uppsala, Sweden

[2]Current address:

[3]Current address:

*Corresponding author


Running title: *Mycobacterium marinum* strain variation

Key words: *M. marinum* ESX genes and genomics, transcription of ESX genes, Smooth and Rough colony morphology, LOS genes, transcription of LOS genes

*Analysis of transcription of selected virulence genes under different growth conditions*

The 1218R strain is more virulent than 1218S[18], and we identified the presence of virulence genes in these two and three other *M. marinum* strains (see main text). In addition to the analysis of ESX and associated gene transcripts we also examined the transcript levels of selected virulence genes to get deeper insight into the differences between 1218R and 1218S. As discussed in the main text, we analysed the transcript levels of these genes detected in exponentially growing and stationary cells and we also included *Mmar*$^{\mathrm{CCUG}}$ and *Mmar*$^{\mathrm{M}}$. The analysis was performed as outlined in the main text and the results are shown Figure S6 and Table S3.

As indicated the main text (Figs 2 and S4, Tables S1 and S3), relative to 1218R we detected some variation in the number of virulence genes in the four strains, *Mmar*$^{\mathrm{1218S}}$ (in addition to the missing ESX-1 genes), *Mmar*$^{\mathrm{CCUG}}$, *Mmar*$^{\mathrm{ATCC}}$ and *Mmar*$^{\mathrm{M}}$. For *Mmar*$^{\mathrm{1218R}}$, *Mmar*$^{\mathrm{1218S}}$, *Mmar*$^{\mathrm{CCUG}}$ and *Mmar*$^{\mathrm{M}}$ the overall transcript patterns of virulence genes were similar with some differences (Fig S6a-g), where the differences were more apparent comparing 1218R with in particular *Mmar*$^{\mathrm{M}}$ (Fig S6g).

The mRNA levels for the well described virulence genes, *devR* (*dosR*) and *devS* (1218R gene id: 01428 and 01429) involved in regulating genes important to establish mycobacterial dormancy[102,103], were higher in 1218R than in 1218S, in particular in stationary cells. Higher levels were also detected for *devR* and *devS* in 1218R relative to *Mmar*$^{\mathrm{CCUG}}$ (particularly in stationary cells; Fig 6e). Comparing 1218R and *Mmar*$^{\mathrm{M}}$ revealed that the *devS* mRNA levels was higher in stationary cells while both *devR* and *devS* were higher in exponential *Mmar*$^{\mathrm{M}}$ cells (Fig S6g). Moreover, the 04801-gene transcript (1218R gene id; Figs S6c, e and g; Table S3), which encodes a PE family protein[104,105], was higher in 1218R compared to 1218S (and *Mmar*$^{\mathrm{CCUG}}$) while only small differences were detected compared to *Mmar*$^{\mathrm{M}}$

(noteworthy, we did detect strain dependent variations in mRNA levels for several PE/PPE genes; not shown).

For *mce*6 genes (mammalian cell entry; 1218R gene ids 00168-00173)[106], the mRNA levels were modestly higher in stationary 1218S cells, albeit the differences were small (Fig S6c; Table S3), while higher levels were detected in 1218R than in $Mmar^{CCUG}$ and $Mmar^{M}$ (both in exponential and stationary cells; Fig S6e and g).

For *lppX* (1218R gene id 01775; putative lipoprotein, LppX, precursor)[21], no significant differences were detected comparing 1218R and 1218S or $Mmar^{CCUG}$ (except that in stationary $Mmar^{CCUG}$ cells the mRNA level was higher; Fig S6e). In contrast, we detected >12 $\log_2$-fold higher mRNA levels in 1218R relative to $Mmar^{M}$. Higher levels were also seen for a putative ESAT-6-like protein (≈10 $\log_2$-fold; 1218R gene id 03733), polyketide synthase type I Pks15/1 (>3 $\log_2$-fold; 1218R gene id 03856) and for several other genes comparing 1218R and the M strain (Fig 6g; Table S3).

Furthermore, relative to 1218R the transcript levels for the virulence-related genes *icl* (isocitrate lyase; 1218R gene id 00746), *hspX_2* (alpha-crystallin; 1218R gene id 03552), and *eis*_2 (enhanced intracellular survival protein; 1218R gene id 03812; note that the mRNA level for this gene is higher in $Mmar^{CCUG}$) were lower in 1218S. By contrast, we noted higher mRNA levels for these genes in $Mmar^{M}$. Transcript levels of ESX-3 genes and for some phospholipase genes, *e.g.*, *plcA*_2 (1218R gene id 01384), were also higher in $Mmar^{M}$. Interestingly, for the M strain we noted higher levels for the transcription factors *sigD* (1218R gene id 01099) and *whiB*3 [in exponential growing cells (1218R gene id 01097)]. WhiB3 is suggested to be involved in the regulation of *whiB*6 expression[15] and transcript levels of *whiB*6 is discussed in the main text. For other differences see Figs S6 and Table S3).

Taken together, the transcript levels for several virulence-related genes differ depend on *Mmar* strain. Given that we did detect differential mRNA levels for several genes involved in the formation of the outer boundaries might suggest variation in the structure of the outer membrane and thereby virulence for the different strains. For example, the four-fold difference in virulence reported for 1218R relative to 1218S[18]. Moreover, the differences observed for the M strain relative to 1218R is consistent with that these strains represent two different *M. marinum* subspecies/lineages[18,unpubl data]. As discussed in the main text, the levels of the major virulence ESX-1 *esxB*_3 (CFP-10) and *esxA*_3 (ESAT-6) transcripts are among the most abundant virulence-related gene transcripts irrespective of strain with the exception of 1218S. In 1218S the ESX-6 genes *esxB*_1 and *esxA*_1 (homologs to *esxB*_3 and *esxA*_3) are among the most abundant transcripts (see main text; Fig S7a, b, d and f).

Mmar *strains transformed with the empty control plasmid (pBS401) or with pBS401$^{espF\text{-}H}$*

The 1218R and 1218S *Mmar* strains were transformed with the empty control plasmid (pBS401) or with pBS401$^{espF\text{-}H}$ and cultivated as outlined in Materials and Methods. The *espF*_2, *espG*$_1$_2 and *espH* genes from 1218R were cloned into the pBS401 plasmid behind an anhydrotetracycline inducible promoter[100] and screened using plasmid specific primers (see Supplementary information). This construct is refered to as pBS401$^{espF\text{-}H}$. The *Mmar* 1218R and 1218S strains were transformed with an empty control plasmid (pBS401) or with pBS401$^{espF\text{-}H}$. The cells were cultivated in microtiter plates for 26 days in 7H9 medium without Tween 80 or hygromycin B. The cultures grown in the 96-well plates were used for the quantification of biofilm formation[101]. Each strain and plasmid combination were measured in two biological replicates, with two technical replicates per biological replicate; in total four measurements per combination.

**Supplementary Information - References**

**Figure legends Supplementary information**

**Figure S1** Genome aligment, analysis of CDS and functional classifications for 1218R, 1218S, *Mmar*$^{CCUG}$, *Mmar*$^{ATCC927}$ and *Mmar*$^{M}$.

(a) Whole-genome alignment for the complete genomes for five *Mmar* strains. The horizontal blocks represent the genomes as marked while vertical "blocks" correspond to homologous regions. White gaps represent insertions/deletions. For details see the main text.

(b) Venn diagram showing the presence of common and unique annotated genes for the five *Mmar* strains.

(c) RAST functional classification of core genes in the five different *Mmar* strains as indicated (3138, 3071, 3075, 3105 and 3149 correspond to classification of the total number of annotated genes in the respective strain). Note that he functionally classified genes were not manually edited, see main text for details.

### Figure S1

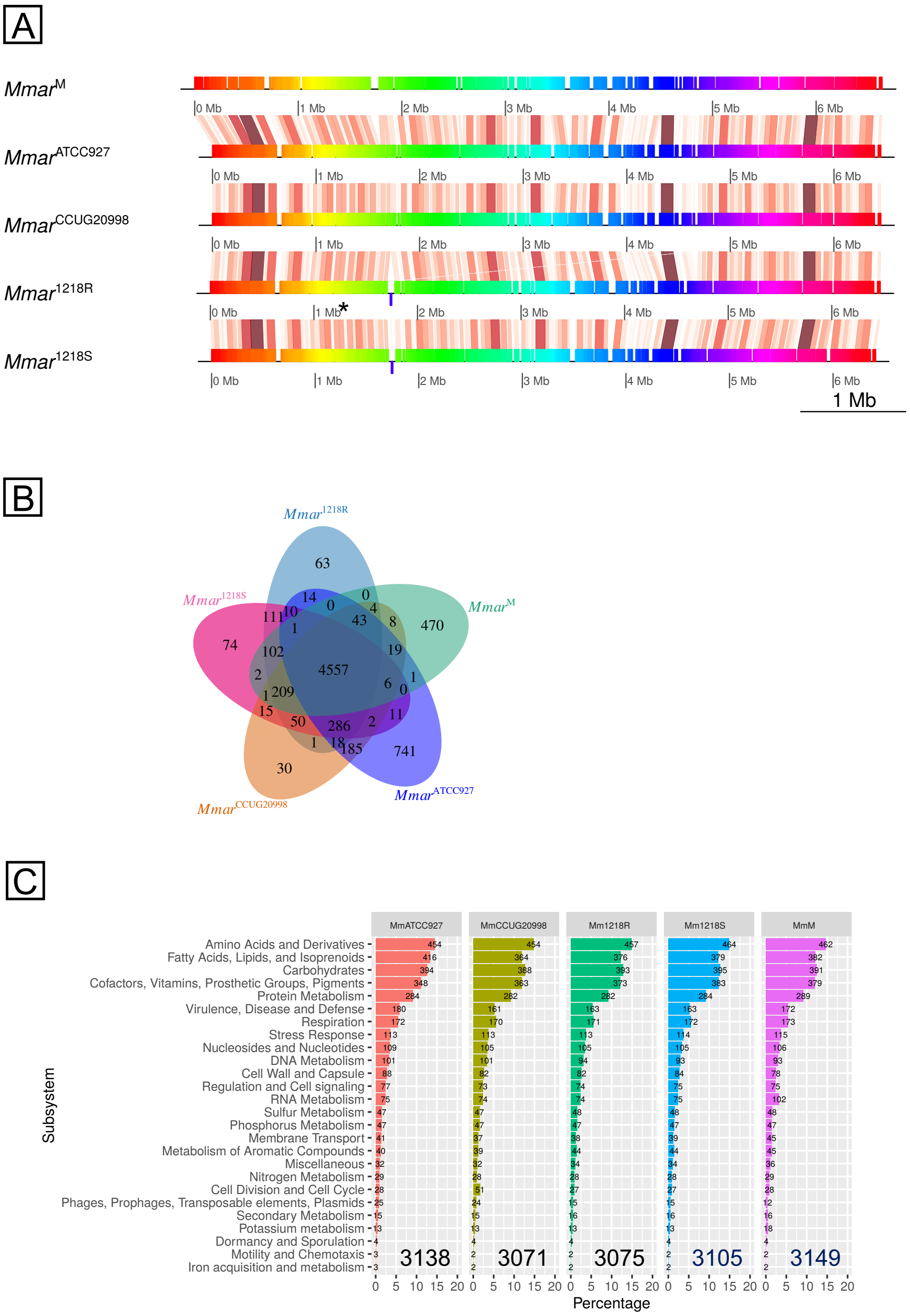

A
Mmar^M
Mmar^ATCC927
Mmar^CCUG20998
Mmar^1218R
Mmar^1218S
0 Mb
1 Mb
2 Mb
3 Mb
4 Mb
5 Mb
6 Mb
1 Mb
B
Mmar^1218R
Mmar^1218S
Mmar^M
Mmar^CCUG20998
Mmar^ATCC927
63
74
470
30
741
4557
C
MmATCC927
MmCCUG20998
Mm1218R
Mm1218S
MmM
Subsystem
Amino Acids and Derivatives
Fatty Acids, Lipids, and Isoprenoids
Carbohydrates
Cofactors, Vitamins, Prosthetic Groups, Pigments
Protein Metabolism
Virulence, Disease and Defense
Respiration
Stress Response
Nucleosides and Nucleotides
DNA Metabolism
Cell Wall and Capsule
Regulation and Cell signaling
RNA Metabolism
Sulfur Metabolism
Phosphorus Metabolism
Membrane Transport
Metabolism of Aromatic Compounds
Miscellaneous
Nitrogen Metabolism
Cell Division and Cell Cycle
Phages, Prophages, Transposable elements, Plasmids
Secondary Metabolism
Potassium metabolism
Dormancy and Sporulation
Motility and Chemotaxis
Iron acquisition and metabolism
3138
3071
3075
3105
3149
Percentage

**Figure S2** Analysis of the ESX genes.

(a) Sequence alignment of ESX-1 genes in 1218R and 1218S.

(b) Sequence alignment of ESX-6 genes in 1218R and 1218S.

(c) Gene synteny of the ESX-3, ESX-4 and ESX-5 in 1218R, 1218S, *Mmar*$^{ATCC927}$, *Mmar*$^{CCUG}$ and *Mmar*$^{M}$ as indicated. Arrows represent genes where blue colours mark genes with known function, grey hypothetical genes.

(d) Sequence alignment of ESX-3 genes in 1218R and 1218S.

(e) Sequence alignment of ESX-4 genes in 1218R and 1218S.

(f) Sequence alignment of ESX-5 genes in 1218R and 1218S.

Fig S2A

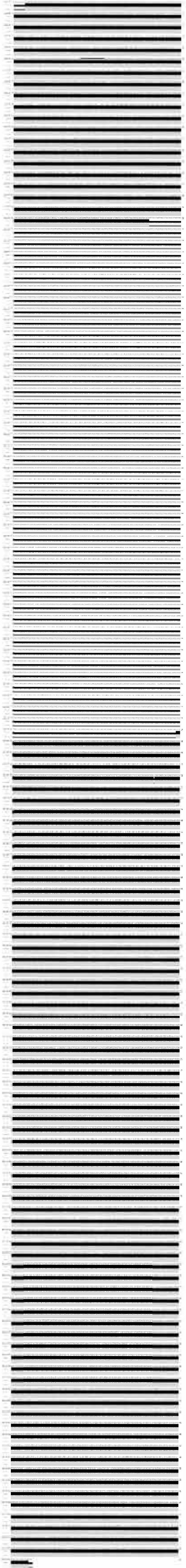

Fig S2B

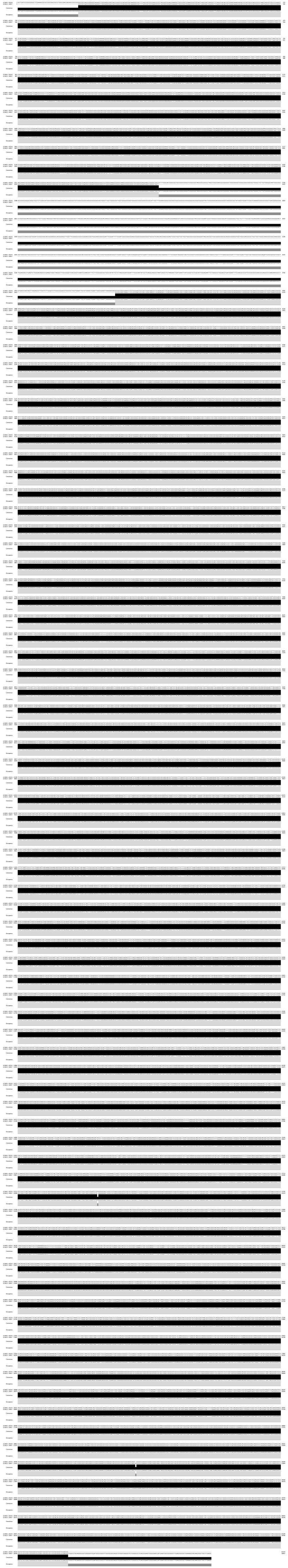

### Figure S2C

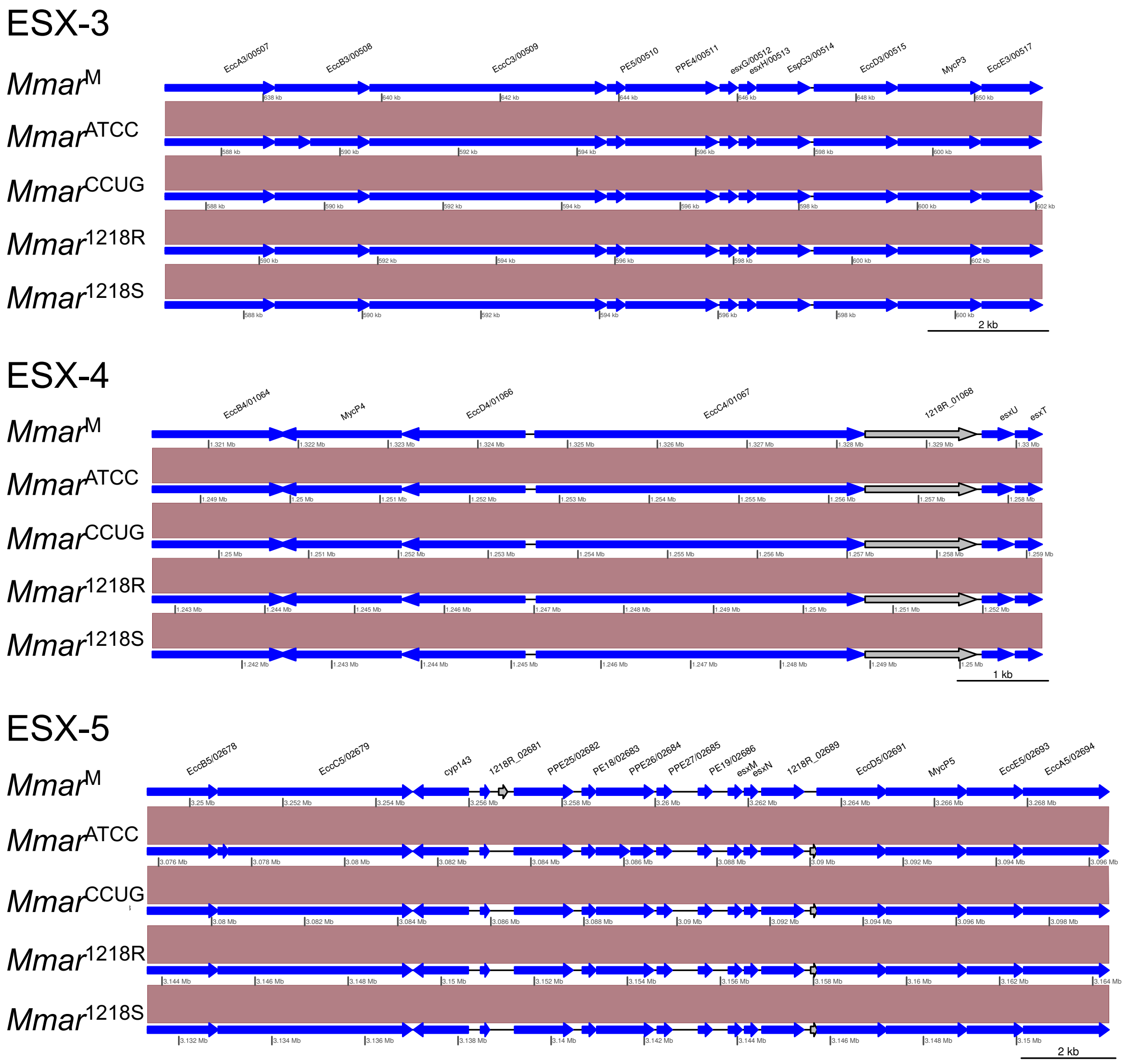
ESX-3
Mmar^M
Mmar^ATCC
Mmar^CCUG
Mmar^1218R
Mmar^1218S
EccA3/00507
EccB3/00508
EccC3/00509
PE5/00510
PPE4/00511
esxG/00512
esxH/00513
EspG3/00514
EccD3/00515
MycP3
EccE3/00517
2 kb
ESX-4
Mmar^M
Mmar^ATCC
Mmar^CCUG
Mmar^1218R
Mmar^1218S
EccB4/01064
MycP4
EccD4/01066
EccC4/01067
1218R_01068
esxU
esxT
1 kb
ESX-5
Mmar^M
Mmar^ATCC
Mmar^CCUG
Mmar^1218R
Mmar^1218S
EccB5/02678
EccC5/02679
cyp143
1218R_02681
PPE25/02682
PE18/02683
PPE26/02684
PPE27/02685
PE19/02686
esxM
esxN
1218R_02689
EccD5/02691
MycP5
EccE5/02693
EccA5/02694
2 kb

Fig S2D

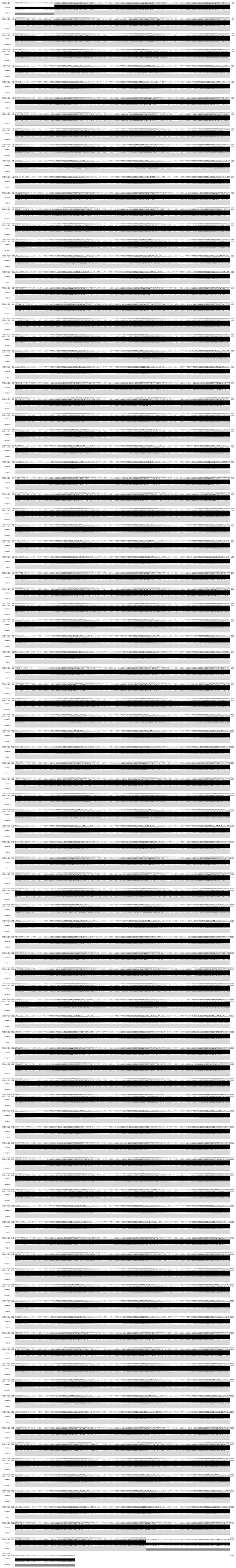

Fig S2E

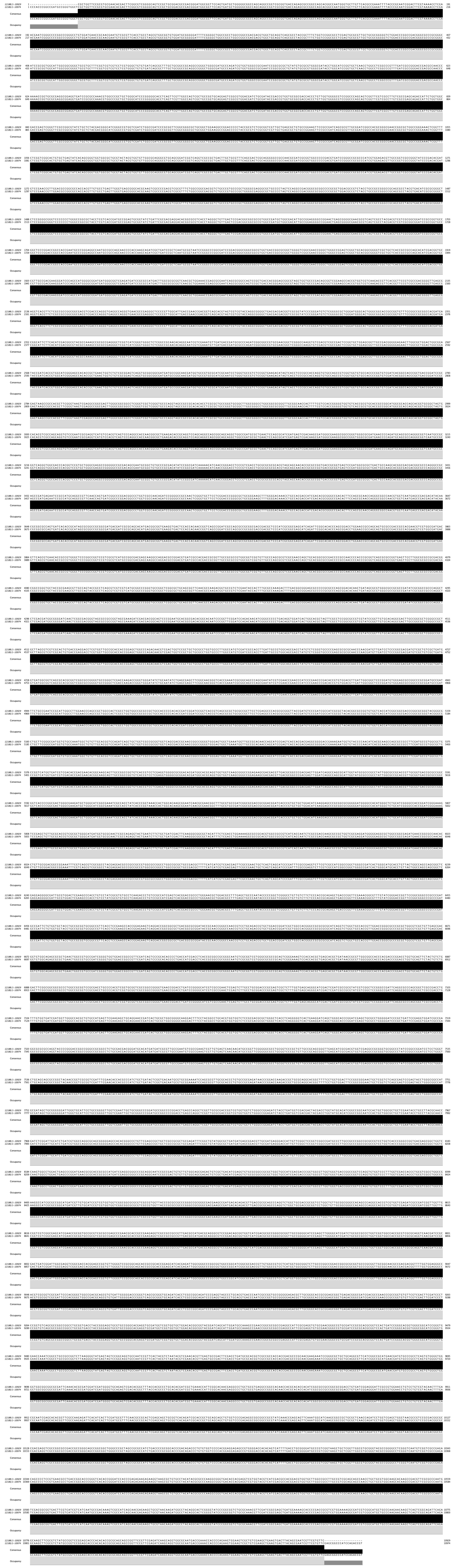

Fig S2F

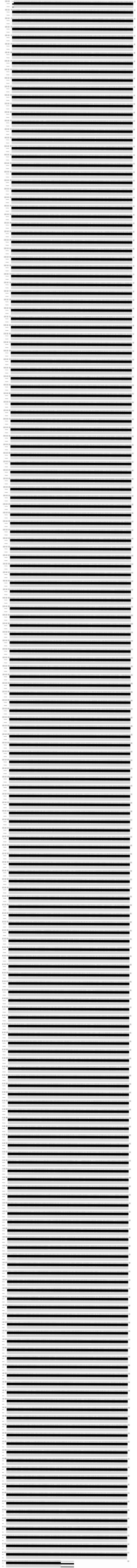

**Figure S3** Illustration of gene differences/annotation for selected genes in 1218R and 1218S.

(a) Gene synteny and annotation of the RNase J gene.

(b) Gene synteny and annotation of *ppsB*_3 (and *ppsB*_1).

(c) Gene synteny reveals deletion of genes in 1218S compared to 1218R.

For further details see Table S1a-d.

### Figure S3

#### A Alignment - RNase J gene

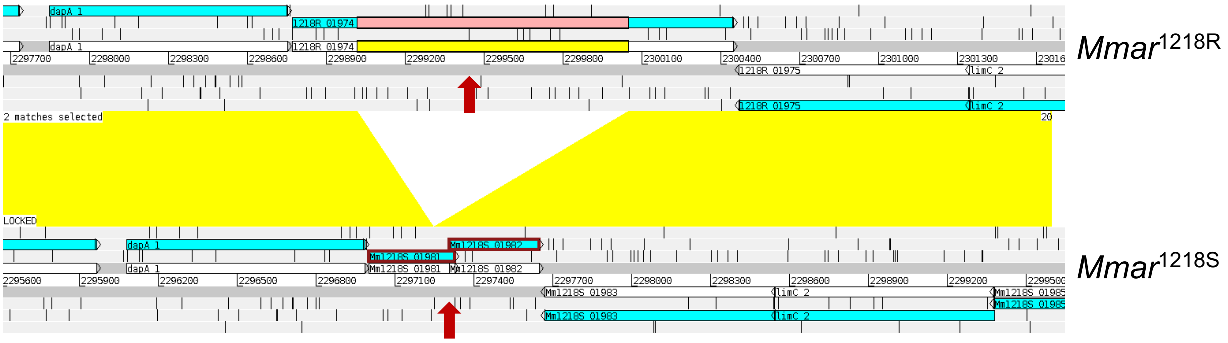


#### B Alignment - *rhlE*

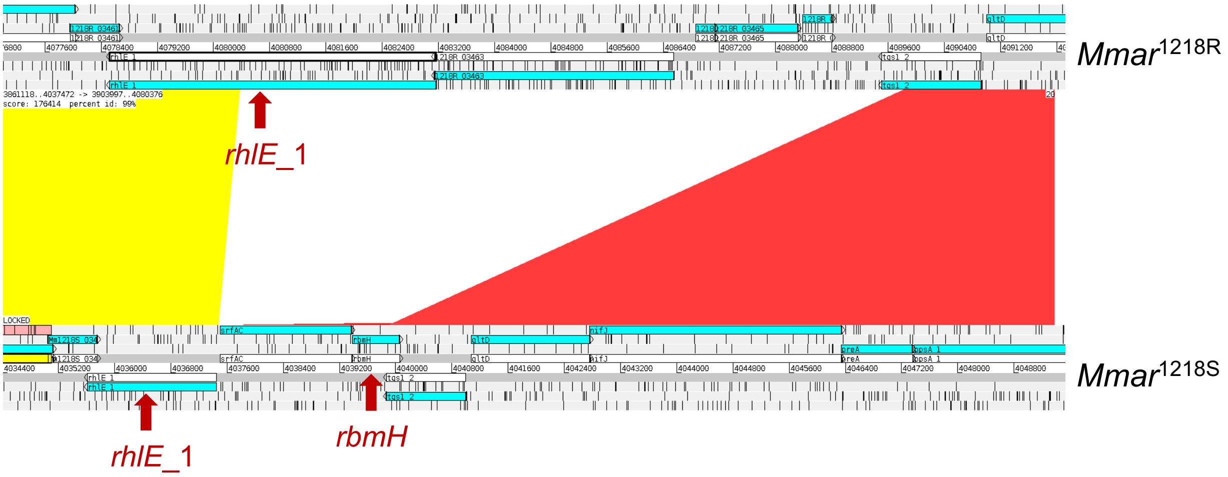


#### C Alignment - *ppsA*_1 and *ppsB*_3 in 1218R and *ppsB*_3 in 1218S

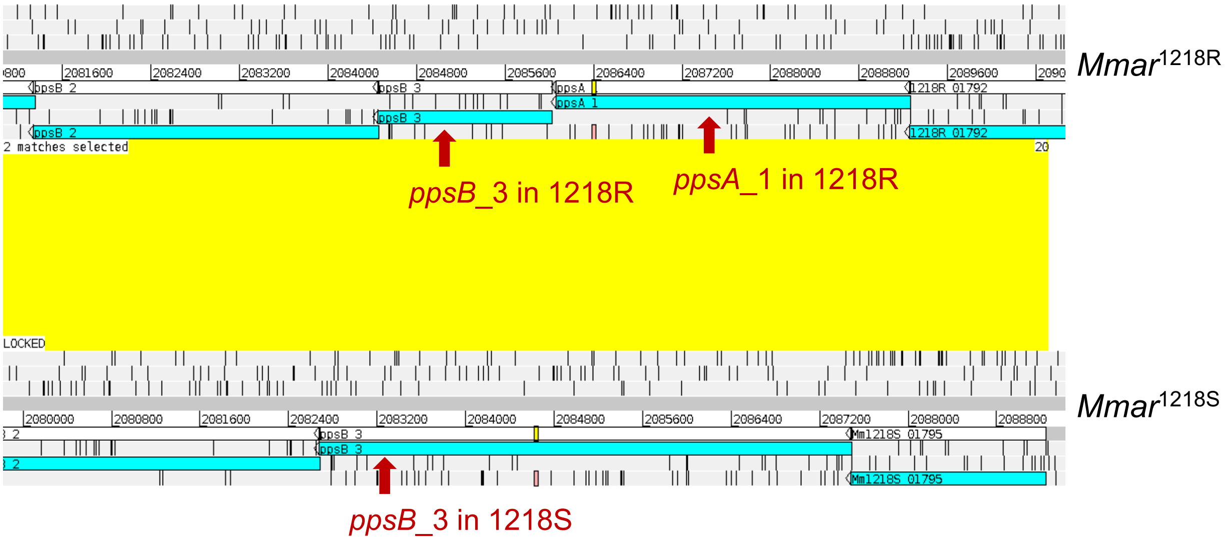

**Figure S4** Identification and classification of virulence-related genes in 1218R, 1218S, *Mmar*$^{ATCC927}$, *Mmar*$^{CCUG}$ and *Mmar*$^{M}$ using the VFanalyzer tool (the VF data base, VFDB; 333, 325, 250, 325 and 345 correspond to the total number of classified annotated virulence-related genes).

### Figure S4

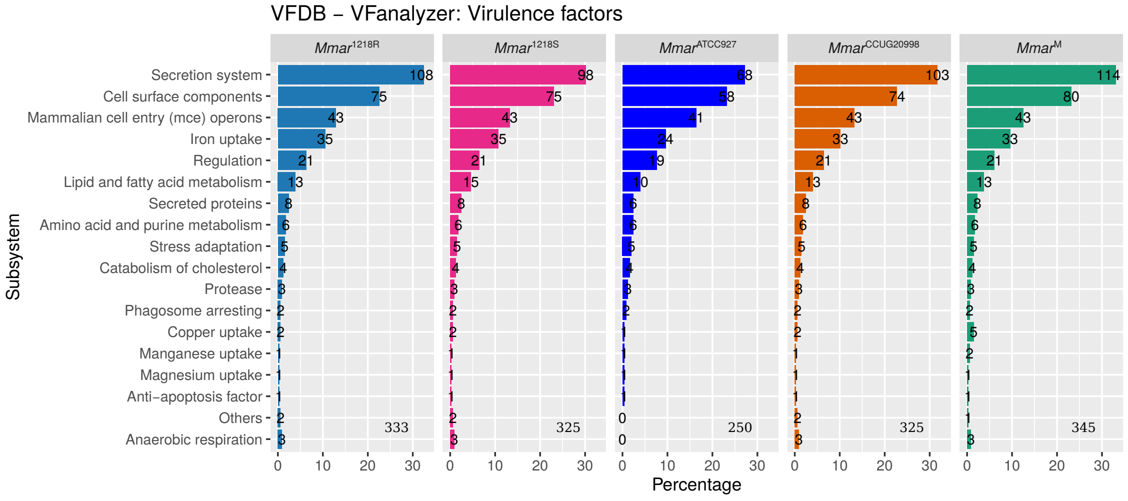

VFDB – VFanalyzer: Virulence factors
Mmar1218R
Mmar1218S
MmarATCC927
MmarCCUG20998
MmarM
Subsystem
Secretion system
Cell surface components
Mammalian cell entry (mce) operons
Iron uptake
Regulation
Lipid and fatty acid metabolism
Secreted proteins
Amino acid and purine metabolism
Stress adaptation
Catabolism of cholesterol
Protease
Phagosome arresting
Copper uptake
Manganese uptake
Magnesium uptake
Anti−apoptosis factor
Others
Anaerobic respiration
333
325
250
325
345
Percentage

**Figure S5** Analysis of LOS and related regulatory genes in 1218R and 1218S.

(a) Sequence alignment of LOS genes, *mutA* to *ileS*.

(b) Sequence alignment of *lsr2*.

(c) Sequence alignment of *whiB*4.

(d) Sequence alignment of *pknL*.

Fig S5A

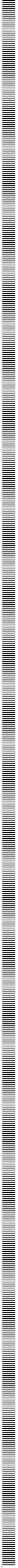

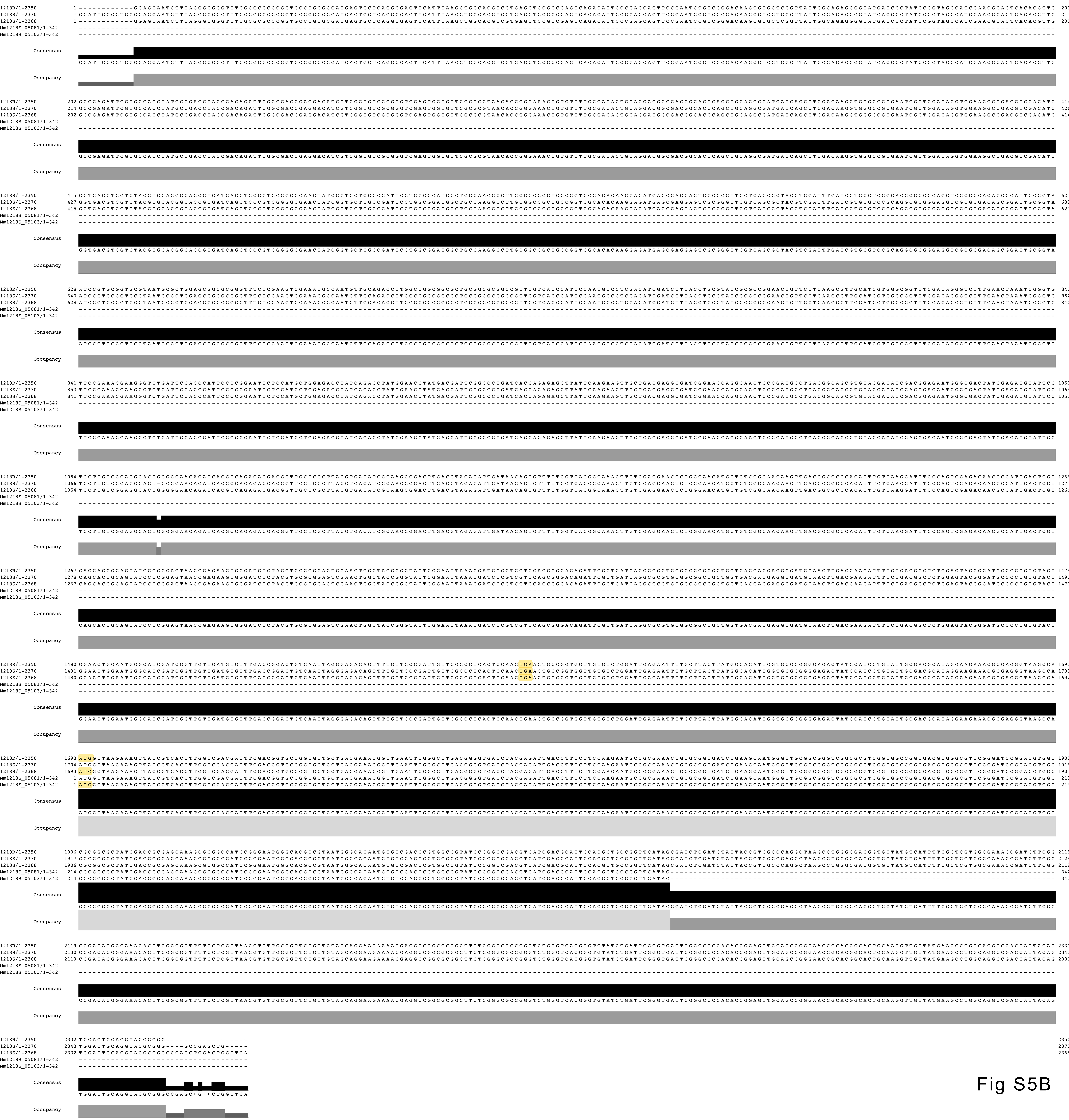
1218R/1-2350
1218S/1-2370
1218S/1-2368
Mm1218S_05081/1-342
Mm1218S_05103/1-342
Consensus
Occupancy

Fig S5B

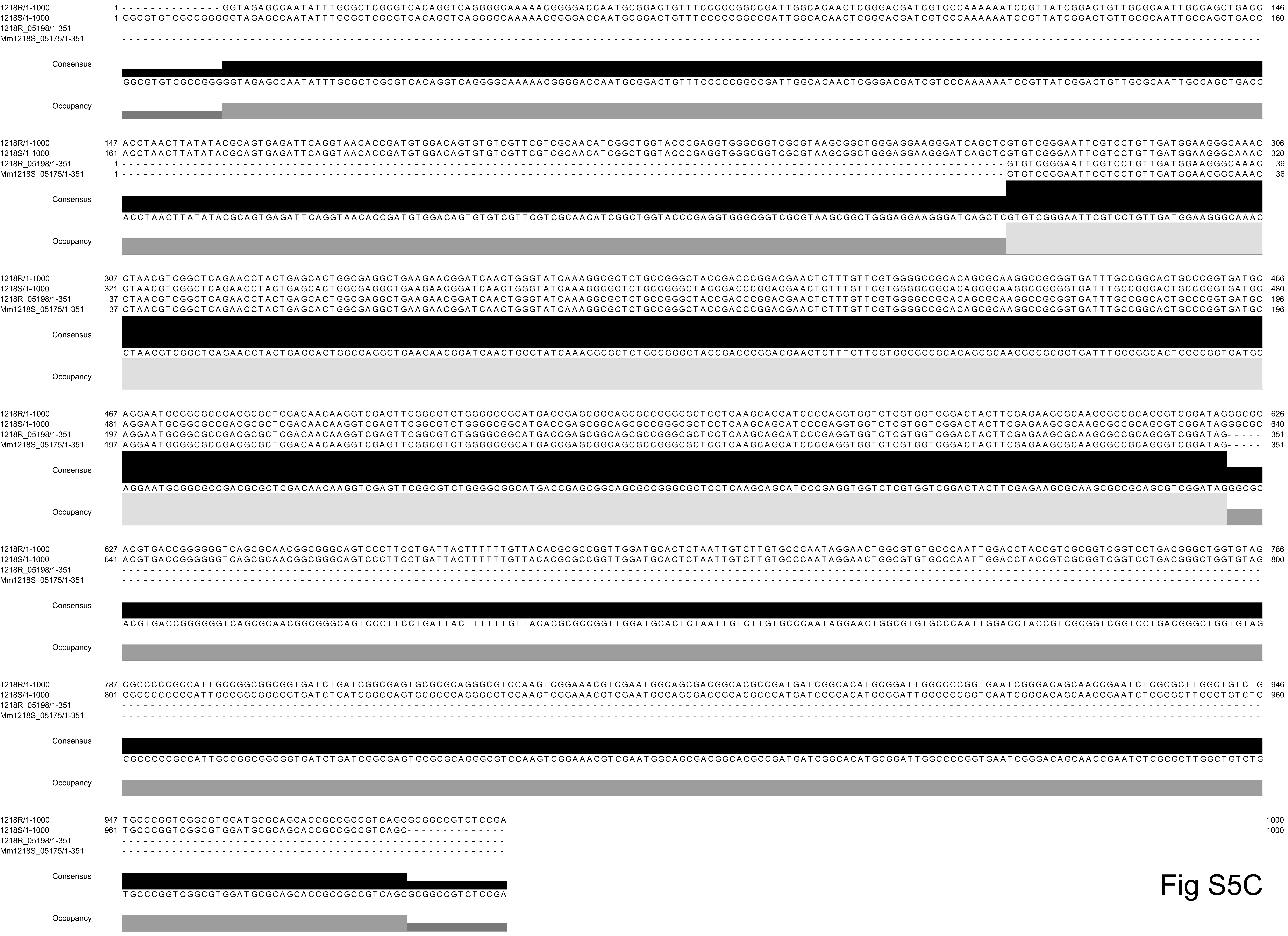
1218R/1-1000
1218S/1-1000
1218R_05198/1-351
Mm1218S_05175/1-351
Consensus
Occupancy

Fig S5C

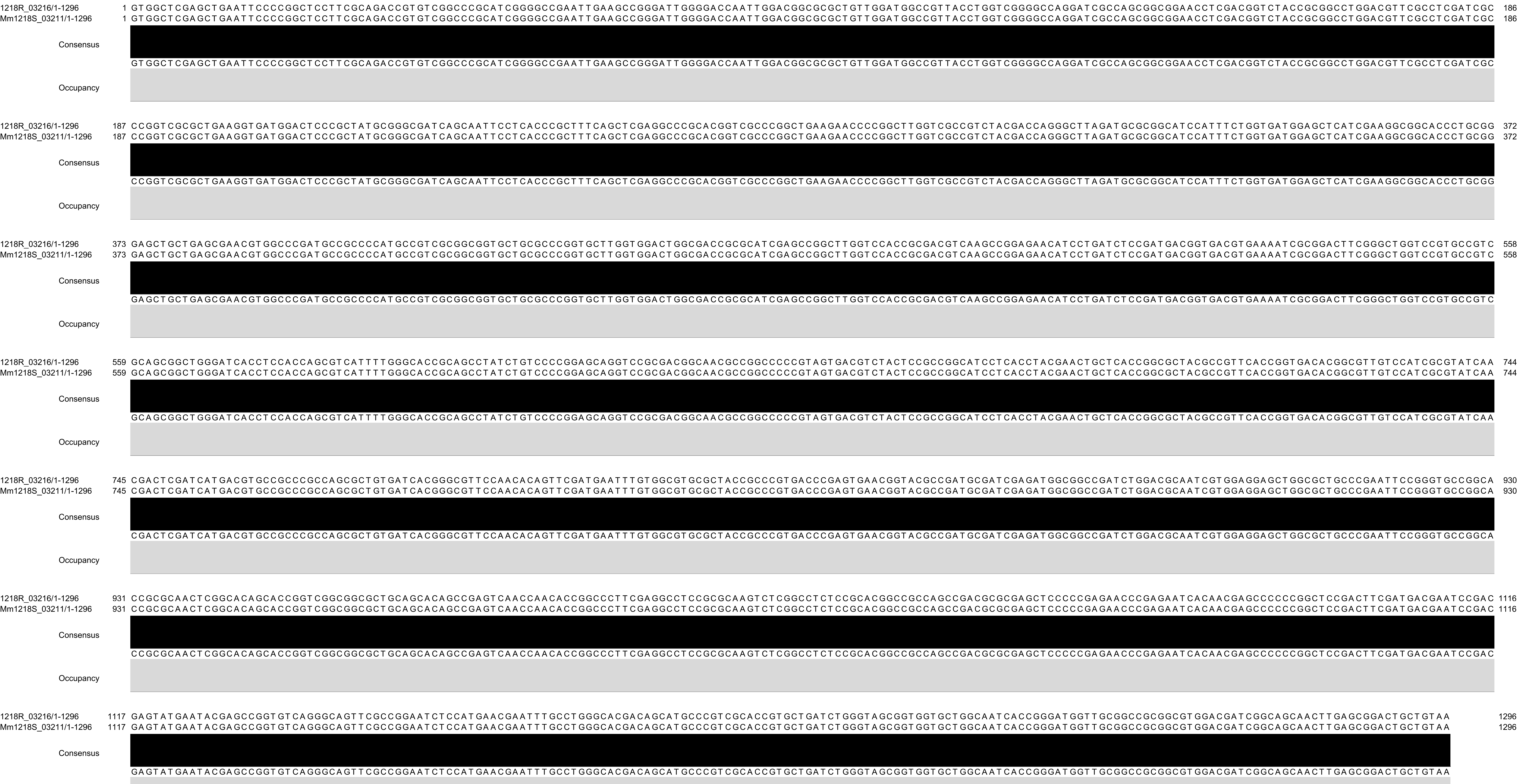
1218R_03216/1-1296 1 GTGGCTCGAGCTGAATTCCCCGGCTCCTTCGCAGACCGTGTCGGCCCGCATCGGGGCCGAATTGAAGCCGGGATTGGGGACCAATTGGACGGCGCGCTGTTGGATGGCCGTTACCTGGTCGGGGCCAGGATCGCCAGCGGCGGAACCTCGACGGTCTACCGCGGCCTGGACGTTCGCCTCGATCGC 186
Mm1218S_03211/1-1296 1 GTGGCTCGAGCTGAATTCCCCGGCTCCTTCGCAGACCGTGTCGGCCCGCATCGGGGCCGAATTGAAGCCGGGATTGGGGACCAATTGGACGGCGCGCTGTTGGATGGCCGTTACCTGGTCGGGGCCAGGATCGCCAGCGGCGGAACCTCGACGGTCTACCGCGGCCTGGACGTTCGCCTCGATCGC 186
Consensus
GTGGCTCGAGCTGAATTCCCCGGCTCCTTCGCAGACCGTGTCGGCCCGCATCGGGGCCGAATTGAAGCCGGGATTGGGGACCAATTGGACGGCGCGCTGTTGGATGGCCGTTACCTGGTCGGGGCCAGGATCGCCAGCGGCGGAACCTCGACGGTCTACCGCGGCCTGGACGTTCGCCTCGATCGC
Occupancy
1218R_03216/1-1296 187 CCGGTCGCGCTGAAGGTGATGGACTCCCGCTATGCGGGCGATCAGCAATTCCTCACCCGCTTTCAGCTCGAGGCCCGCACGGTCGCCCGGCTGAAGAACCCCGGCTTGGTCGCCGTCTACGACCAGGGCTTAGATGCGCGGCATCCATTTCTGGTGATGGAGCTCATCGAAGGCGGCACCCTGCGG 372
Mm1218S_03211/1-1296 187 CCGGTCGCGCTGAAGGTGATGGACTCCCGCTATGCGGGCGATCAGCAATTCCTCACCCGCTTTCAGCTCGAGGCCCGCACGGTCGCCCGGCTGAAGAACCCCGGCTTGGTCGCCGTCTACGACCAGGGCTTAGATGCGCGGCATCCATTTCTGGTGATGGAGCTCATCGAAGGCGGCACCCTGCGG 372
Consensus
CCGGTCGCGCTGAAGGTGATGGACTCCCGCTATGCGGGCGATCAGCAATTCCTCACCCGCTTTCAGCTCGAGGCCCGCACGGTCGCCCGGCTGAAGAACCCCGGCTTGGTCGCCGTCTACGACCAGGGCTTAGATGCGCGGCATCCATTTCTGGTGATGGAGCTCATCGAAGGCGGCACCCTGCGG
Occupancy
1218R_03216/1-1296 373 GAGCTGCTGAGCGAACGTGGCCCGATGCCGCCCCATGCCGTCGCGGCGGTGCTGCGCCCGGTGCTTGGTGGACTGGCGACCGCGCATCGAGCCGGCTTGGTCCACCGCGACGTCAAGCCGGAGAACATCCTGATCTCCGATGACGGTGACGTGAAAATCGCGGACTTCGGGCTGGTCCGTGCCGTC 558
Mm1218S_03211/1-1296 373 GAGCTGCTGAGCGAACGTGGCCCGATGCCGCCCCATGCCGTCGCGGCGGTGCTGCGCCCGGTGCTTGGTGGACTGGCGACCGCGCATCGAGCCGGCTTGGTCCACCGCGACGTCAAGCCGGAGAACATCCTGATCTCCGATGACGGTGACGTGAAAATCGCGGACTTCGGGCTGGTCCGTGCCGTC 558
Consensus
GAGCTGCTGAGCGAACGTGGCCCGATGCCGCCCCATGCCGTCGCGGCGGTGCTGCGCCCGGTGCTTGGTGGACTGGCGACCGCGCATCGAGCCGGCTTGGTCCACCGCGACGTCAAGCCGGAGAACATCCTGATCTCCGATGACGGTGACGTGAAAATCGCGGACTTCGGGCTGGTCCGTGCCGTC
Occupancy
1218R_03216/1-1296 559 GCAGCGGCTGGGATCACCTCCACCAGCGTCATTTTGGGCACCGCAGCCTATCTGTCCCCGGAGCAGGTCCGCGACGGCAACGCCGGCCCCCGTAGTGACGTCTACTCCGCCGGCATCCTCACCTACGAACTGCTCACCGGCGCTACGCCGTTCACCGGTGACACGGCGTTGTCCATCGCGTATCAA 744
Mm1218S_03211/1-1296 559 GCAGCGGCTGGGATCACCTCCACCAGCGTCATTTTGGGCACCGCAGCCTATCTGTCCCCGGAGCAGGTCCGCGACGGCAACGCCGGCCCCCGTAGTGACGTCTACTCCGCCGGCATCCTCACCTACGAACTGCTCACCGGCGCTACGCCGTTCACCGGTGACACGGCGTTGTCCATCGCGTATCAA 744
Consensus
GCAGCGGCTGGGATCACCTCCACCAGCGTCATTTTGGGCACCGCAGCCTATCTGTCCCCGGAGCAGGTCCGCGACGGCAACGCCGGCCCCCGTAGTGACGTCTACTCCGCCGGCATCCTCACCTACGAACTGCTCACCGGCGCTACGCCGTTCACCGGTGACACGGCGTTGTCCATCGCGTATCAA
Occupancy
1218R_03216/1-1296 745 CGACTCGATCATGACGTGCCGCCCGCCAGCGCTGTGATCACGGGCGTTCCAACACAGTTCGATGAATTTGTGGCGTGCGCTACCGCCCGTGACCCGAGTGAACGGTACGCCGATGCGATCGAGATGGCGGCCGATCTGGACGCAATCGTGGAGGAGCTGGCGCTGCCCGAATTCCGGGTGCCGGCA 930
Mm1218S_03211/1-1296 745 CGACTCGATCATGACGTGCCGCCCGCCAGCGCTGTGATCACGGGCGTTCCAACACAGTTCGATGAATTTGTGGCGTGCGCTACCGCCCGTGACCCGAGTGAACGGTACGCCGATGCGATCGAGATGGCGGCCGATCTGGACGCAATCGTGGAGGAGCTGGCGCTGCCCGAATTCCGGGTGCCGGCA 930
Consensus
CGACTCGATCATGACGTGCCGCCCGCCAGCGCTGTGATCACGGGCGTTCCAACACAGTTCGATGAATTTGTGGCGTGCGCTACCGCCCGTGACCCGAGTGAACGGTACGCCGATGCGATCGAGATGGCGGCCGATCTGGACGCAATCGTGGAGGAGCTGGCGCTGCCCGAATTCCGGGTGCCGGCA
Occupancy
1218R_03216/1-1296 931 CCGCGCAACTCGGCACAGCACCGGTCGGCGGCGCTGCAGCACAGCCGAGTCAACCAACACCGGCCCTTCGAGGCCTCCGCGCAAGTCTCGGCCTCTCCGCACGGCCGCCAGCCGACGCGCGAGCTCCCCCGAGAACCCGAGAATCACAACGAGCCCCCCGGCTCCGACTTCGATGACGAATCCGAC 1116
Mm1218S_03211/1-1296 931 CCGCGCAACTCGGCACAGCACCGGTCGGCGGCGCTGCAGCACAGCCGAGTCAACCAACACCGGCCCTTCGAGGCCTCCGCGCAAGTCTCGGCCTCTCCGCACGGCCGCCAGCCGACGCGCGAGCTCCCCCGAGAACCCGAGAATCACAACGAGCCCCCCGGCTCCGACTTCGATGACGAATCCGAC 1116
Consensus
CCGCGCAACTCGGCACAGCACCGGTCGGCGGCGCTGCAGCACAGCCGAGTCAACCAACACCGGCCCTTCGAGGCCTCCGCGCAAGTCTCGGCCTCTCCGCACGGCCGCCAGCCGACGCGCGAGCTCCCCCGAGAACCCGAGAATCACAACGAGCCCCCCGGCTCCGACTTCGATGACGAATCCGAC
Occupancy
1218R_03216/1-1296 1117 GAGTATGAATACGAGCCGGTGTCAGGGCAGTTCGCCGGAATCTCCATGAACGAATTTGCCTGGGCACGACAGCATGCCCGTCGCACCGTGCTGATCTGGGTAGCGGTGGTGCTGGCAATCACCGGGATGGTTGCGGCCGCGGCGTGGACGATCGGCAGCAACTTGAGCGGACTGCTGTAA 1296
Mm1218S_03211/1-1296 1117 GAGTATGAATACGAGCCGGTGTCAGGGCAGTTCGCCGGAATCTCCATGAACGAATTTGCCTGGGCACGACAGCATGCCCGTCGCACCGTGCTGATCTGGGTAGCGGTGGTGCTGGCAATCACCGGGATGGTTGCGGCCGCGGCGTGGACGATCGGCAGCAACTTGAGCGGACTGCTGTAA 1296
Consensus
GAGTATGAATACGAGCCGGTGTCAGGGCAGTTCGCCGGAATCTCCATGAACGAATTTGCCTGGGCACGACAGCATGCCCGTCGCACCGTGCTGATCTGGGTAGCGGTGGTGCTGGCAATCACCGGGATGGTTGCGGCCGCGGCGTGGACGATCGGCAGCAACTTGAGCGGACTGCTGTAA
Occupancy

Fig S5D

**Figure S6** Bar plots showing transcription of virulence genes in 1218R, 1218S, *Mmar*$^{CCUG}$ and *Mmar*$^{M}$ .

(a) Transcript levels (distribution, expressed in TPM values) in exponentially growing and stationary 1218R cells.

(b) Transcript levels (distribution, expressed in TPM values) in exponentially growing and stationary 1218S cells.

(c) Comparing transcript levels in exponentially growing (red colour) and stationary (turquoise) 1218R vs 1218S cells. Negative $\log_2$-values suggest that the corresponding mRNA is more abundant in 1218S cells while a positive value suggests higher levels in 1218R cells.

(d) Transcript levels (distribution, expressed in TPM values) in exponentially growing and stationary *Mmar*$^{CCUG}$ cells.

(e) Comparing transcript levels in exponentially growing (red colour) and stationary (turquoise) 1218R vs *Mmar*$^{CCUG}$ cells. Negative $\log_2$-values suggest that the corresponding mRNA is more abundant in *Mmar*$^{CCUG}$ cells while a positive value suggests higher levels in 1218R cells.

(f) Transcript levels (distribution, expressed in TPM values) in exponentially growing and stationary *Mmar*$^{M}$ cells.

(g) Comparing transcript levels in exponentially growing (red colour) and stationary (turquoise) 1218R vs *Mmar*$^{M}$ cells. Negative $\log_2$-values suggest that the corresponding mRNA is more abundant in *Mmar*$^{M}$ cells while a positive value suggests higher levels in 1218R cells.

Statistical significance, see Materials and Methods; *p-value < 0.05; **p-value < 0.01; ***p-value < 0.001.

Fig S6A

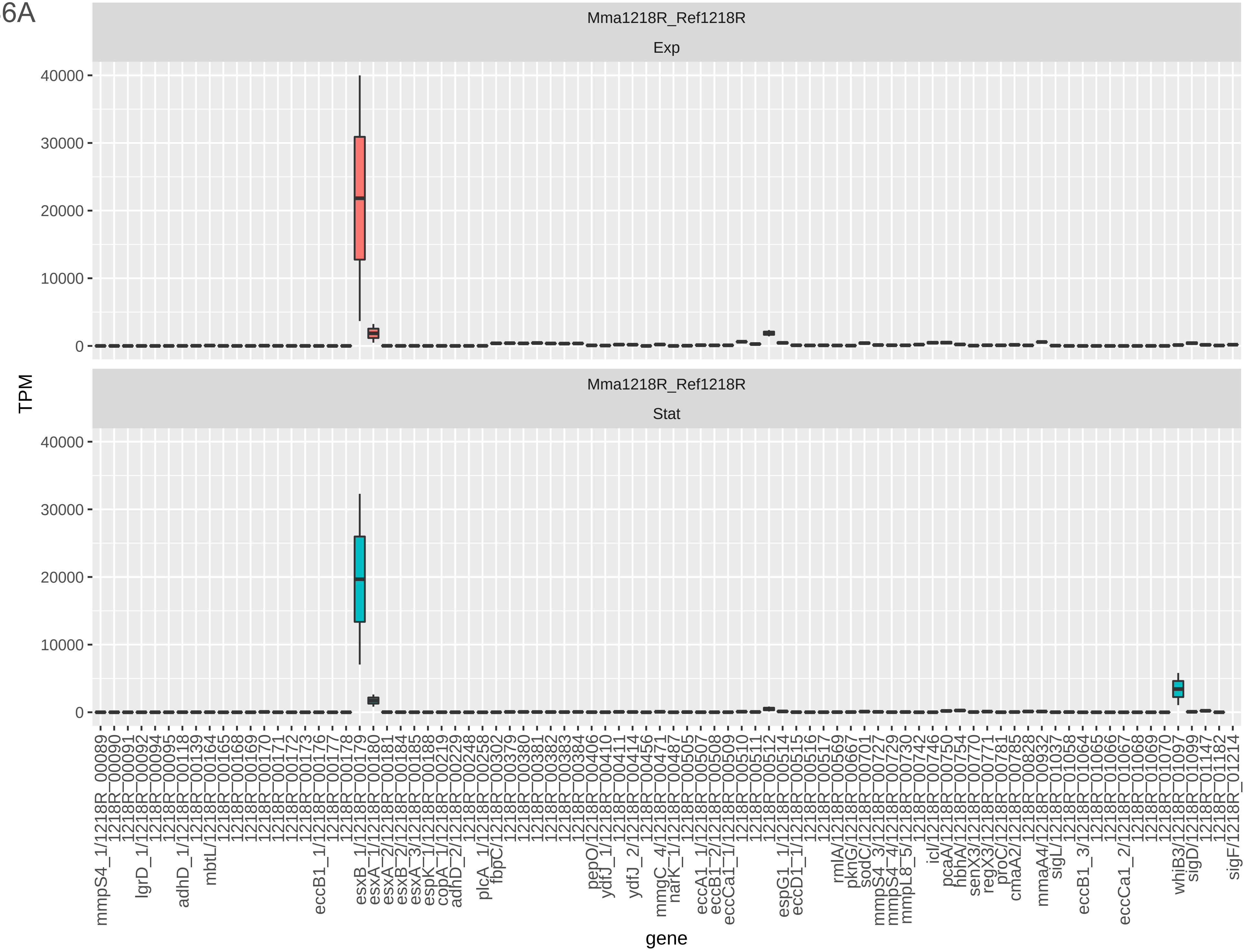

Mma1218R_Ref1218R
Exp
Mma1218R_Ref1218R
Stat
TPM
40000
30000
20000
10000
0
gene
mmpS4_1/1218R_00089
1218R_00090
1218R_00091
lgrD_1/1218R_00092
1218R_00094
1218R_00095
adhD_1/1218R_00118
1218R_00139
mbtL/1218R_00164
1218R_00165
1218R_00168
1218R_00169
1218R_00170
1218R_00171
1218R_00172
1218R_00173
1218R_00176
eccB1_1/1218R_00177
1218R_00178
esxB_1/1218R_00179
esxA_1/1218R_00180
esxA_2/1218R_00181
esxB_2/1218R_00184
esxA_3/1218R_00185
espK_1/1218R_00188
copA_1/1218R_00219
adhD_2/1218R_00229
1218R_00248
plcA_1/1218R_00258
fbpC/1218R_00302
1218R_00379
1218R_00380
1218R_00381
1218R_00382
1218R_00383
1218R_00384
pepO/1218R_00406
ydfJ_1/1218R_00410
1218R_00411
ydfJ_2/1218R_00414
1218R_00456
mmgC_4/1218R_00471
narK_1/1218R_00487
1218R_00505
eccA1_1/1218R_00507
eccB1_2/1218R_00508
eccCa1_1/1218R_00509
1218R_00510
1218R_00511
1218R_00512
espG1_1/1218R_00514
eccD1_1/1218R_00515
1218R_00516
1218R_00517
rmlA/1218R_00569
pknG/1218R_00667
sodC/1218R_00701
mmpS4_3/1218R_00727
mmpS4_4/1218R_00729
mmpL8_5/1218R_00730
1218R_00742
icl/1218R_00746
pcaA/1218R_00750
hbhA/1218R_00754
senX3/1218R_00770
regX3/1218R_00771
proC/1218R_00781
cmaA2/1218R_00785
1218R_00828
mmaA4/1218R_00932
sigL/1218R_01037
1218R_01058
eccB1_3/1218R_01064
1218R_01065
1218R_01066
eccCa1_2/1218R_01067
1218R_01068
1218R_01069
1218R_01070
whiB3/1218R_01097
sigD/1218R_01099
1218R_01147
1218R_01182
sigF/1218R_01214

Fig S6A

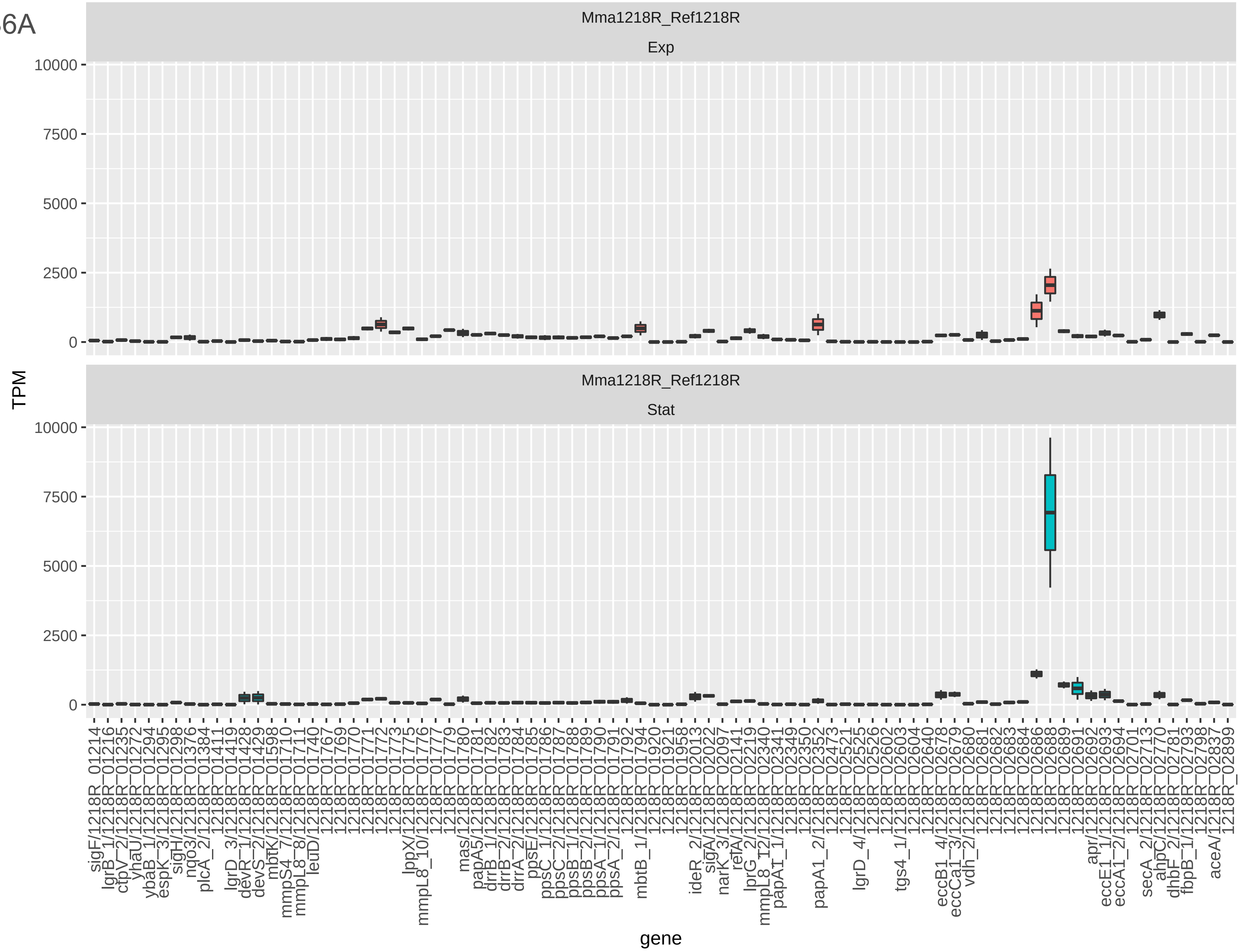
Mma1218R_Ref1218R
Exp
Mma1218R_Ref1218R
Stat
TPM
10000
7500
5000
2500
0
gene

Fig S6A

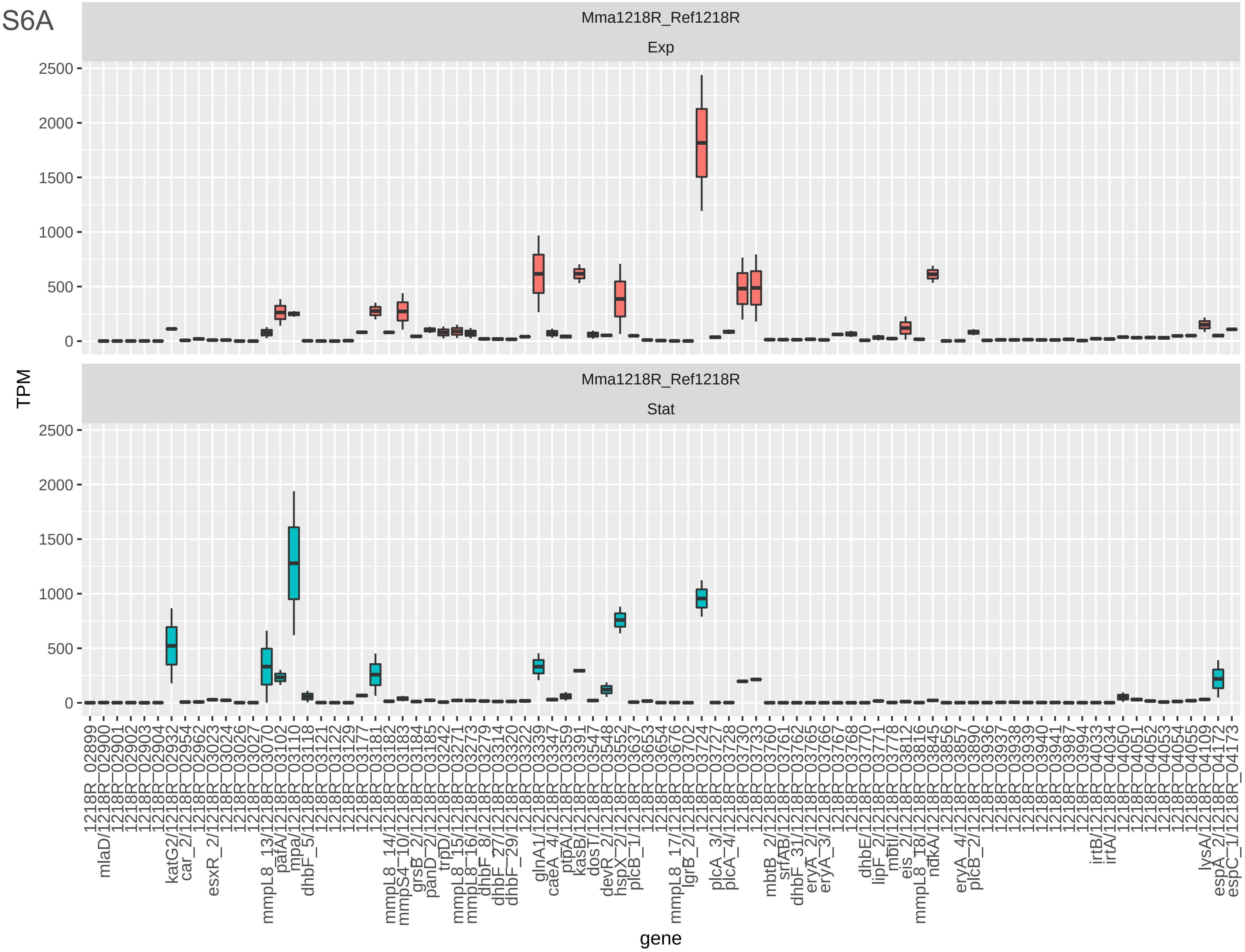

Mma1218R_Ref1218R
Exp
Mma1218R_Ref1218R
Stat
TPM
0
500
1000
1500
2000
2500
gene
1218R_02899
mlaD/1218R_02900
1218R_02901
1218R_02902
1218R_02903
1218R_02904
katG2/1218R_02932
car_2/1218R_02954
1218R_02962
esxR_2/1218R_03023
1218R_03024
1218R_03026
1218R_03027
mmpL8_13/1218R_03070
pafA/1218R_03102
mpa/1218R_03110
dhbF_5/1218R_03118
1218R_03121
1218R_03122
1218R_03129
1218R_03177
mmpL8_14/1218R_03181
mmpS4_10/1218R_03182
grsB_2/1218R_03183
panD_2/1218R_03184
trpD/1218R_03185
mmpL8_15/1218R_03242
mmpL8_16/1218R_03271
dhbF_8/1218R_03273
dhbF_27/1218R_03279
dhbF_29/1218R_03314
1218R_03320
1218R_03322
glnA1/1218R_03339
caeA_4/1218R_03347
ptpA/1218R_03359
kasB/1218R_03391
dosT/1218R_03547
devR_2/1218R_03548
hspX_2/1218R_03552
plcB_1/1218R_03637
1218R_03653
1218R_03654
mmpL8_17/1218R_03676
lgrB_2/1218R_03702
1218R_03724
plcA_3/1218R_03727
plcA_4/1218R_03728
1218R_03730
1218R_03733
mbtB_2/1218R_03760
srfAB/1218R_03761
dhbF_31/1218R_03762
eryA_2/1218R_03765
eryA_3/1218R_03766
1218R_03767
1218R_03768
dhbE/1218R_03770
lipF_2/1218R_03771
mbtI/1218R_03778
eis_2/1218R_03812
mmpL8_18/1218R_03816
ndkA/1218R_03845
1218R_03856
eryA_4/1218R_03857
plcB_2/1218R_03890
1218R_03936
1218R_03937
1218R_03938
1218R_03939
1218R_03940
1218R_03941
1218R_03987
1218R_03994
irtB/1218R_04033
irtA/1218R_04034
1218R_04050
1218R_04051
1218R_04052
1218R_04053
1218R_04054
1218R_04055
lysA/1218R_04109
espA_2/1218R_04172
espC_1/1218R_04173

Fig S6A

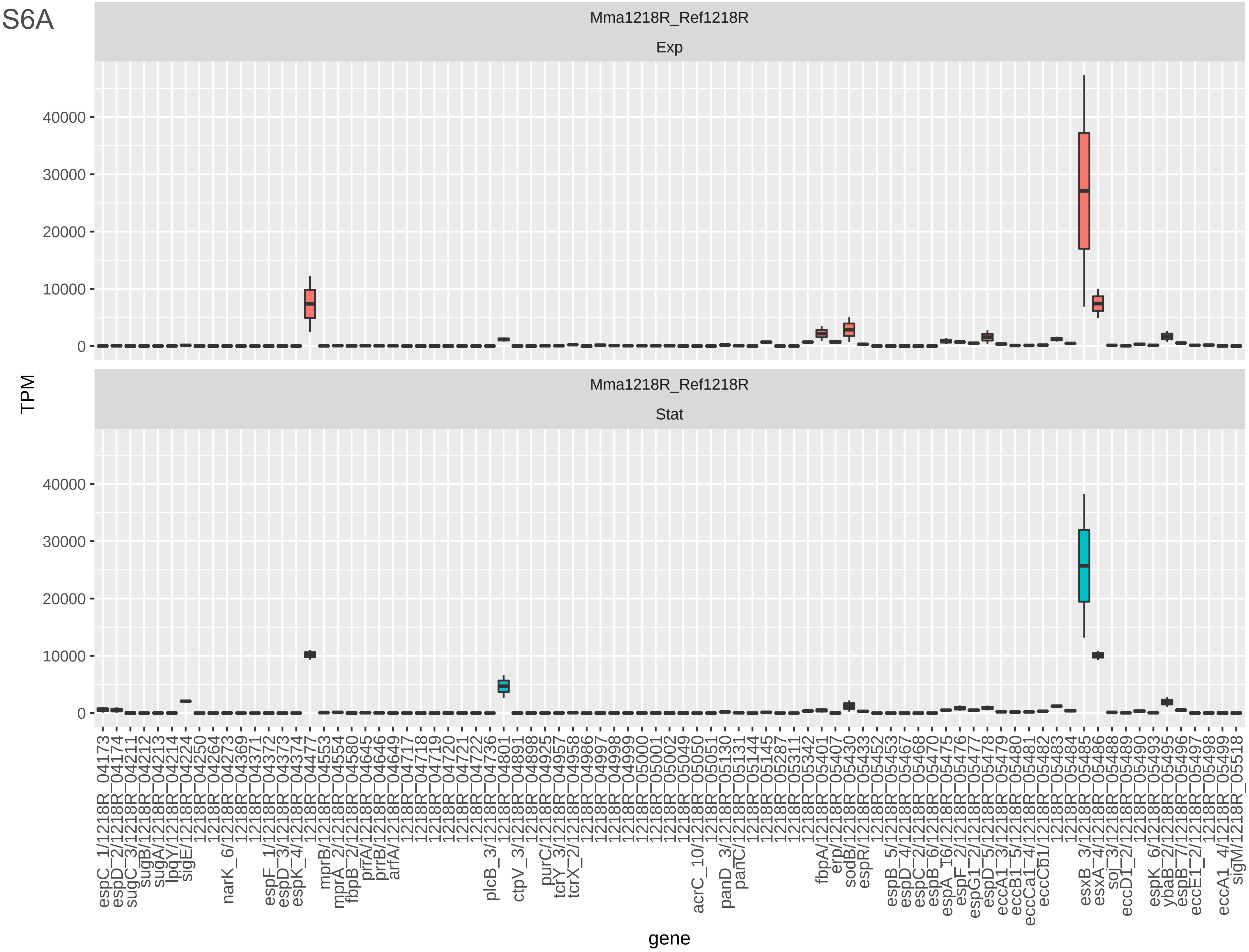

Mma1218R_Ref1218R
Exp
Mma1218R_Ref1218R
Stat
TPM
40000
30000
20000
10000
0
gene
espC_1/1218R_04173
espD_2/1218R_04174
sugC_3/1218R_04211
sugB/1218R_04212
sugA/1218R_04213
lpqY/1218R_04214
sigE/1218R_04224
1218R_04250
1218R_04264
narK_6/1218R_04273
1218R_04369
1218R_04371
espF_1/1218R_04372
espD_3/1218R_04373
espK_4/1218R_04374
1218R_04477
mprB/1218R_04553
mprA_2/1218R_04554
fbpB_2/1218R_04580
prrA/1218R_04645
prrB/1218R_04646
arfA/1218R_04649
1218R_04717
1218R_04718
1218R_04719
1218R_04720
1218R_04721
1218R_04722
plcB_3/1218R_04736
1218R_04801
ctpV_3/1218R_04891
1218R_04898
purC/1218R_04925
tcrY_3/1218R_04957
tcrX_2/1218R_04958
1218R_04986
1218R_04997
1218R_04998
1218R_04999
1218R_05000
1218R_05001
1218R_05002
1218R_05049
acrC_10/1218R_05050
1218R_05051
panD_3/1218R_05130
panC/1218R_05131
1218R_05144
1218R_05145
1218R_05287
1218R_05311
1218R_05342
fbpA/1218R_05401
erp/1218R_05407
sodB/1218R_05430
espR/1218R_05433
1218R_05452
espB_5/1218R_05453
espD_4/1218R_05467
espC_2/1218R_05468
espB_6/1218R_05470
espA_I6/1218R_05475
espF_2/1218R_05476
espG1_2/1218R_05477
espD_5/1218R_05478
eccA1_3/1218R_05479
eccB1_5/1218R_05480
eccCa1_4/1218R_05481
eccCb1/1218R_05482
1218R_05483
1218R_05484
esxB_3/1218R_05485
esxA_4/1218R_05486
soj_3/1218R_05488
eccD1_2/1218R_05489
1218R_05490
espK_6/1218R_05493
ybaB_2/1218R_05495
espB_7/1218R_05496
eccE1_2/1218R_05497
1218R_05498
eccA1_4/1218R_05499
sigM/1218R_05518

Fig S6B

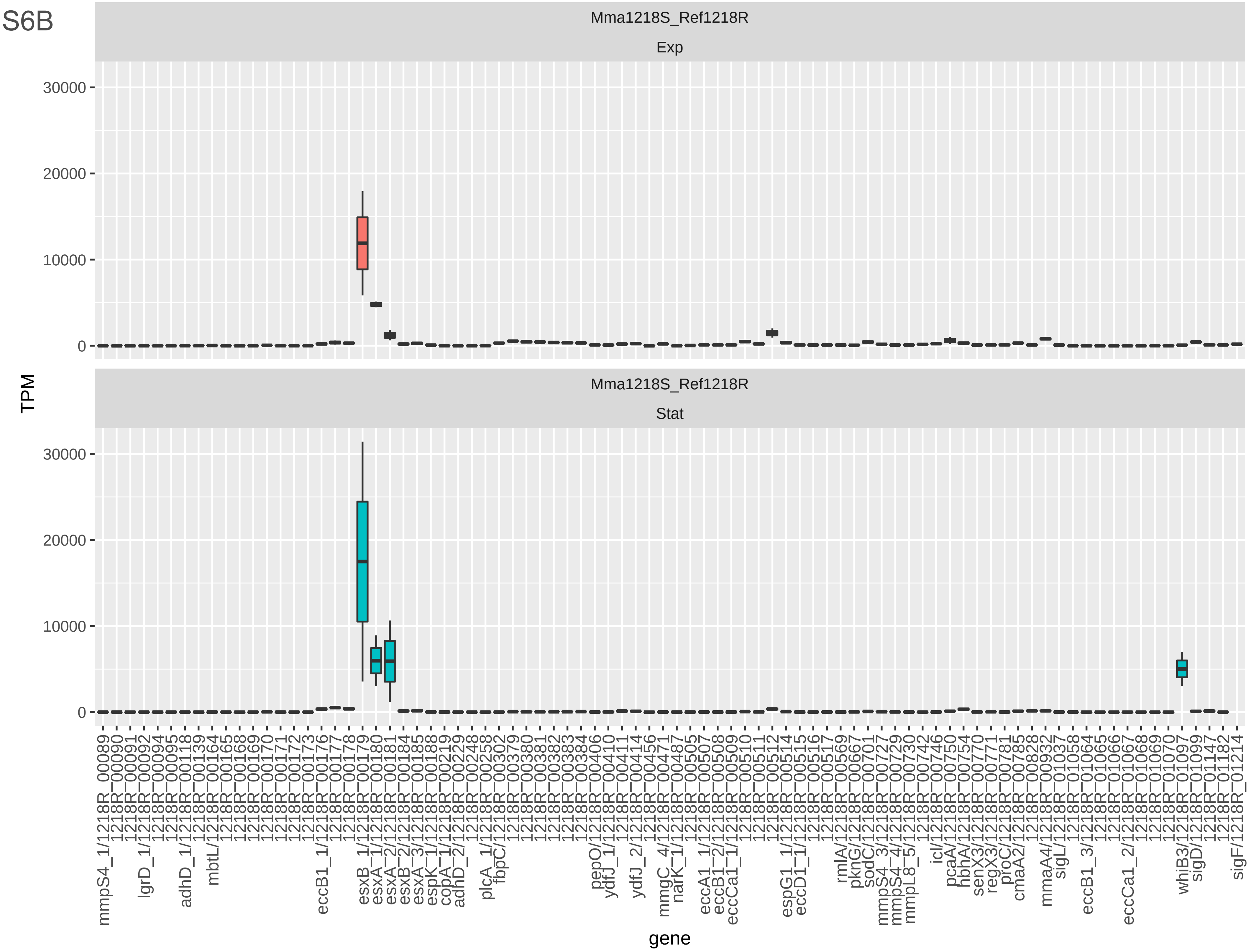
Mma1218S_Ref1218R
Exp
Mma1218S_Ref1218R
Stat
TPM
30000
20000
10000
0
gene
mmpS4_1/1218R_00089
1218R_00090
1218R_00091
lgrD_1/1218R_00092
1218R_00094
1218R_00095
adhD_1/1218R_00118
1218R_00139
mbtL/1218R_00164
1218R_00165
1218R_00168
1218R_00169
1218R_00170
1218R_00171
1218R_00172
1218R_00173
eccB1_1/1218R_00176
1218R_00177
1218R_00178
esxB_1/1218R_00179
esxA_1/1218R_00180
esxA_2/1218R_00181
esxB_2/1218R_00184
esxA_3/1218R_00185
espK_1/1218R_00188
copA_1/1218R_00219
adhD_2/1218R_00229
1218R_00248
plcA_1/1218R_00258
fbpC/1218R_00302
1218R_00379
1218R_00380
1218R_00381
1218R_00382
1218R_00383
1218R_00384
pepO/1218R_00406
ydfJ_1/1218R_00410
1218R_00411
ydfJ_2/1218R_00414
1218R_00456
mmgC_4/1218R_00471
narK_1/1218R_00487
1218R_00505
eccA1_1/1218R_00507
eccB1_2/1218R_00508
eccCa1_1/1218R_00509
1218R_00510
1218R_00511
1218R_00512
espG1_1/1218R_00514
eccD1_1/1218R_00515
1218R_00516
1218R_00517
rmlA/1218R_00569
pknG/1218R_00667
sodC/1218R_00701
mmpS4_3/1218R_00727
mmpS4_4/1218R_00729
mmpL8_5/1218R_00730
1218R_00742
icl/1218R_00746
pcaA/1218R_00750
hbhA/1218R_00754
senX3/1218R_00770
regX3/1218R_00771
proC/1218R_00781
cmaA2/1218R_00785
1218R_00828
mmaA4/1218R_00932
sigL/1218R_01037
1218R_01058
eccB1_3/1218R_01064
1218R_01065
1218R_01066
eccCa1_2/1218R_01067
1218R_01068
1218R_01069
1218R_01070
whiB3/1218R_01097
sigD/1218R_01099
1218R_01147
1218R_01182
sigF/1218R_01214

Fig S6B

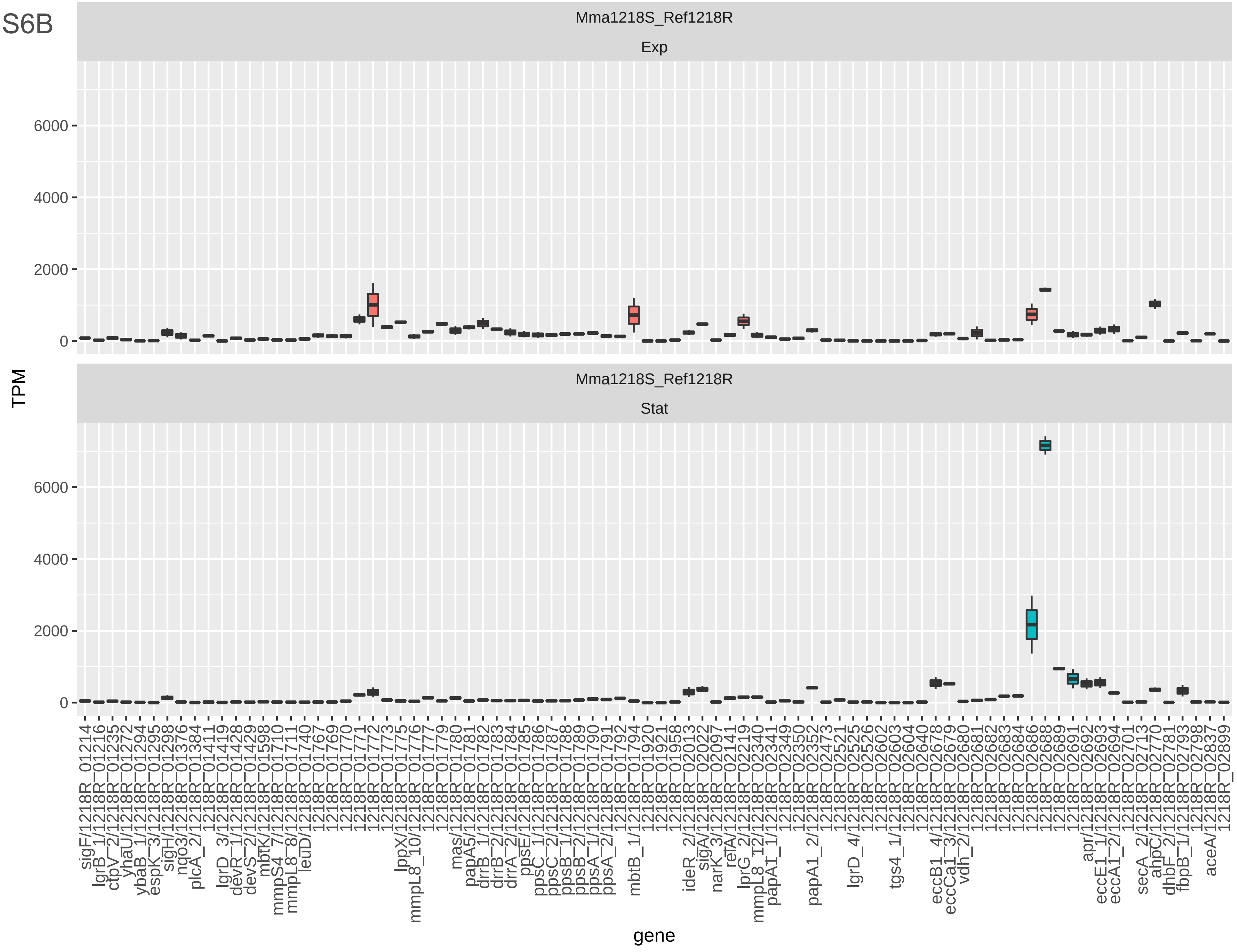
Mma1218S_Ref1218R
Exp
Mma1218S_Ref1218R
Stat
TPM
0
2000
4000
6000
gene
sigF/1218R_01214
lgrB_1/1218R_01216
ctpV_2/1218R_01235
yhaU/1218R_01272
ybaB_1/1218R_01294
espK_3/1218R_01295
sigH/1218R_01298
ngo3/1218R_01376
plcA_2/1218R_01384
1218R_01411
lgrD_3/1218R_01419
devR_1/1218R_01428
devS_2/1218R_01429
mbtK/1218R_01598
mmpS4_7/1218R_01710
mmpL8_8/1218R_01711
leuD/1218R_01740
1218R_01767
1218R_01769
1218R_01770
1218R_01771
1218R_01772
1218R_01773
lppX/1218R_01775
mmpL8_10/1218R_01776
1218R_01777
1218R_01779
mas/1218R_01780
papA5/1218R_01781
drrB_1/1218R_01782
drrB_2/1218R_01783
drrA_2/1218R_01784
ppsE/1218R_01785
ppsC_1/1218R_01786
ppsC_2/1218R_01787
ppsB_1/1218R_01788
ppsB_2/1218R_01789
ppsA_1/1218R_01790
ppsA_2/1218R_01791
1218R_01792
mbtB_1/1218R_01794
1218R_01920
1218R_01921
1218R_01958
ideR_2/1218R_02013
sigA/1218R_02022
narK_3/1218R_02097
relA/1218R_02141
lprG_2/1218R_02219
mmpL8_12/1218R_02340
papA1_1/1218R_02341
1218R_02349
1218R_02350
papA1_2/1218R_02352
1218R_02473
1218R_02521
lgrD_4/1218R_02525
1218R_02526
1218R_02602
tgs4_1/1218R_02603
1218R_02604
1218R_02640
eccB1_4/1218R_02678
eccCa1_3/1218R_02679
vdh_2/1218R_02680
1218R_02681
1218R_02682
1218R_02683
1218R_02684
1218R_02686
1218R_02688
1218R_02689
apr/1218R_02691
eccE1_1/1218R_02692
eccA1_2/1218R_02693
1218R_02694
1218R_02701
secA_2/1218R_02713
ahpC/1218R_02770
dhbF_2/1218R_02781
fbpB_1/1218R_02793
1218R_02798
aceA/1218R_02837
1218R_02899

Fig S6B

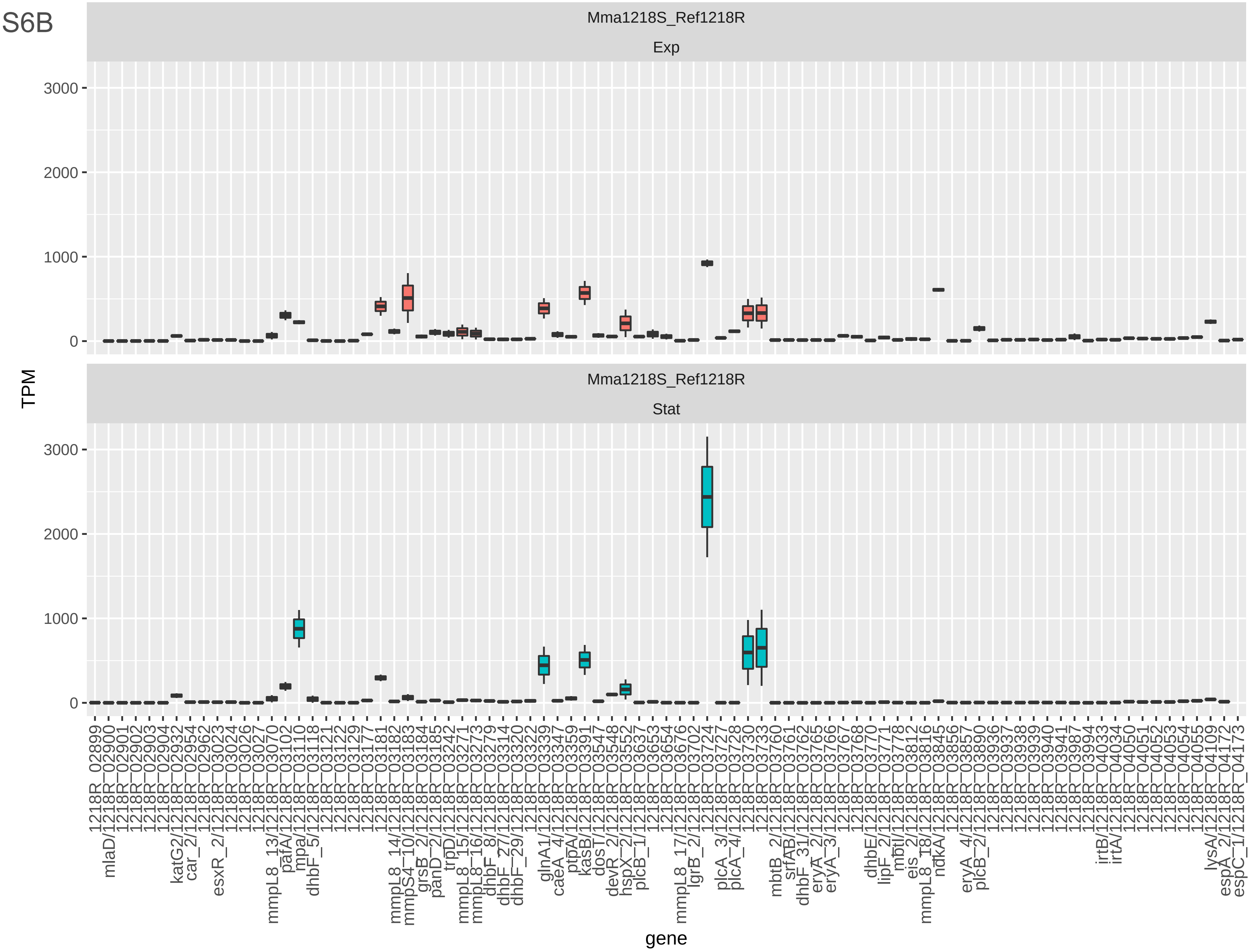

Mma1218S_Ref1218R
Exp
Mma1218S_Ref1218R
Stat
3000
2000
1000
0
TPM
gene

Fig S6B

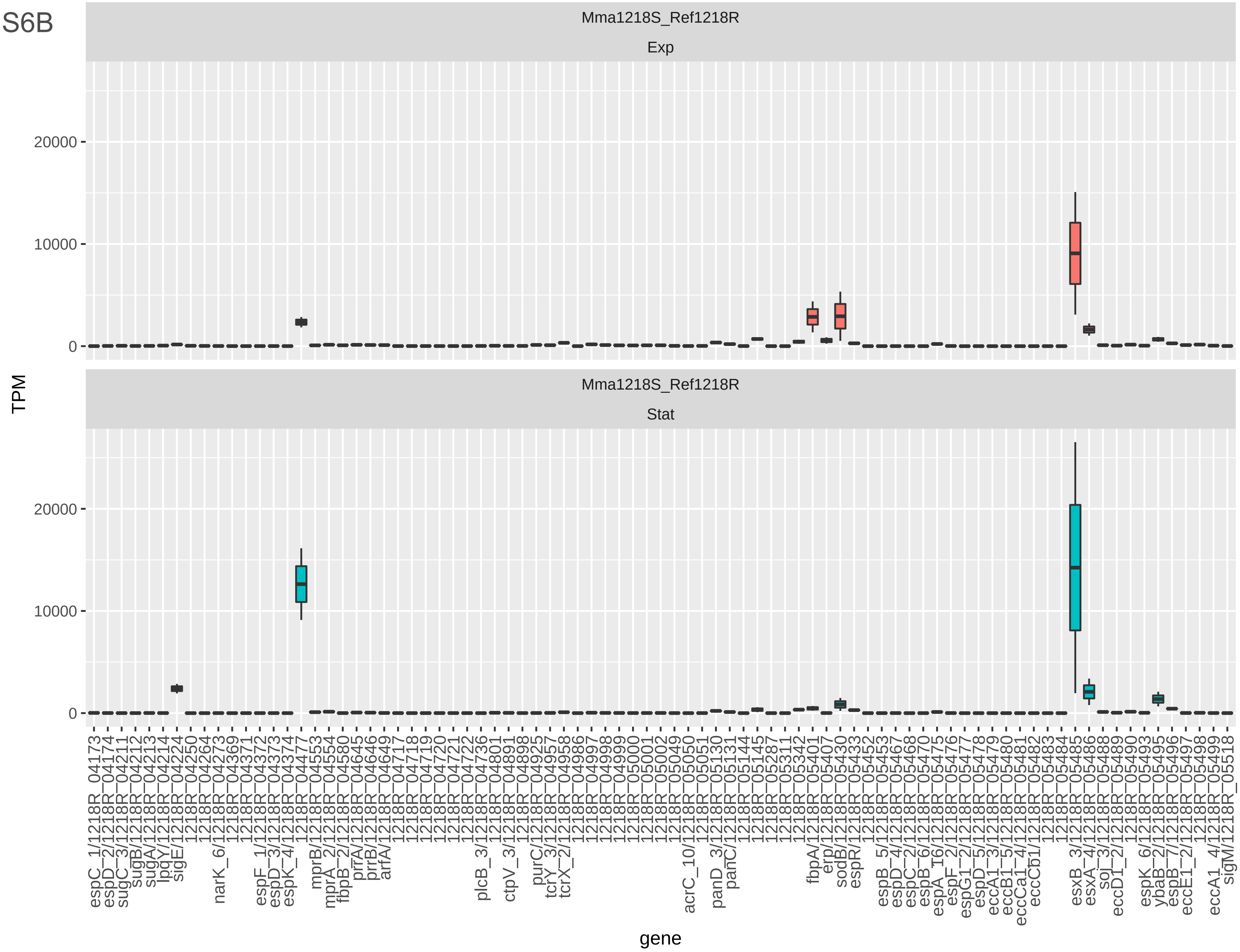
Mma1218S_Ref1218R
Exp
Mma1218S_Ref1218R
Stat
TPM
0
10000
20000
gene
espC_1/1218R_04173
espD_2/1218R_04174
sugC_3/1218R_04211
sugB/1218R_04212
sugA/1218R_04213
lpqY/1218R_04214
sigE/1218R_04224
1218R_04250
1218R_04264
narK_6/1218R_04273
1218R_04369
1218R_04371
espF_1/1218R_04372
espD_3/1218R_04373
espK_4/1218R_04374
1218R_04477
mprB/1218R_04553
mprA_2/1218R_04554
fbpB_2/1218R_04580
prrA/1218R_04645
prrB/1218R_04646
arfA/1218R_04649
1218R_04717
1218R_04718
1218R_04719
1218R_04720
1218R_04721
1218R_04722
plcB_3/1218R_04736
1218R_04801
ctpV_3/1218R_04891
1218R_04898
purC/1218R_04925
tcrY_3/1218R_04957
tcrX_2/1218R_04958
1218R_04986
1218R_04997
1218R_04998
1218R_04999
1218R_05000
1218R_05001
1218R_05002
1218R_05049
acrC_10/1218R_05050
1218R_05051
panD_3/1218R_05130
panC/1218R_05131
1218R_05144
1218R_05145
1218R_05287
1218R_05311
1218R_05342
fbpA/1218R_05401
erp/1218R_05407
sodB/1218R_05430
espR/1218R_05433
1218R_05452
espB_5/1218R_05453
espD_4/1218R_05467
espC_2/1218R_05468
espB_6/1218R_05470
espA_16/1218R_05475
espF_2/1218R_05476
espG1_2/1218R_05477
espD_5/1218R_05478
eccA1_3/1218R_05479
eccB1_5/1218R_05480
eccCa1_4/1218R_05481
eccCb1/1218R_05482
1218R_05483
1218R_05484
esxB_3/1218R_05485
esxA_4/1218R_05486
soj_3/1218R_05488
eccD1_2/1218R_05489
1218R_05490
espK_6/1218R_05493
ybaB_2/1218R_05495
espB_7/1218R_05496
eccE1_2/1218R_05497
1218R_05498
eccA1_4/1218R_05499
sigM/1218R_05518

Fig S6C

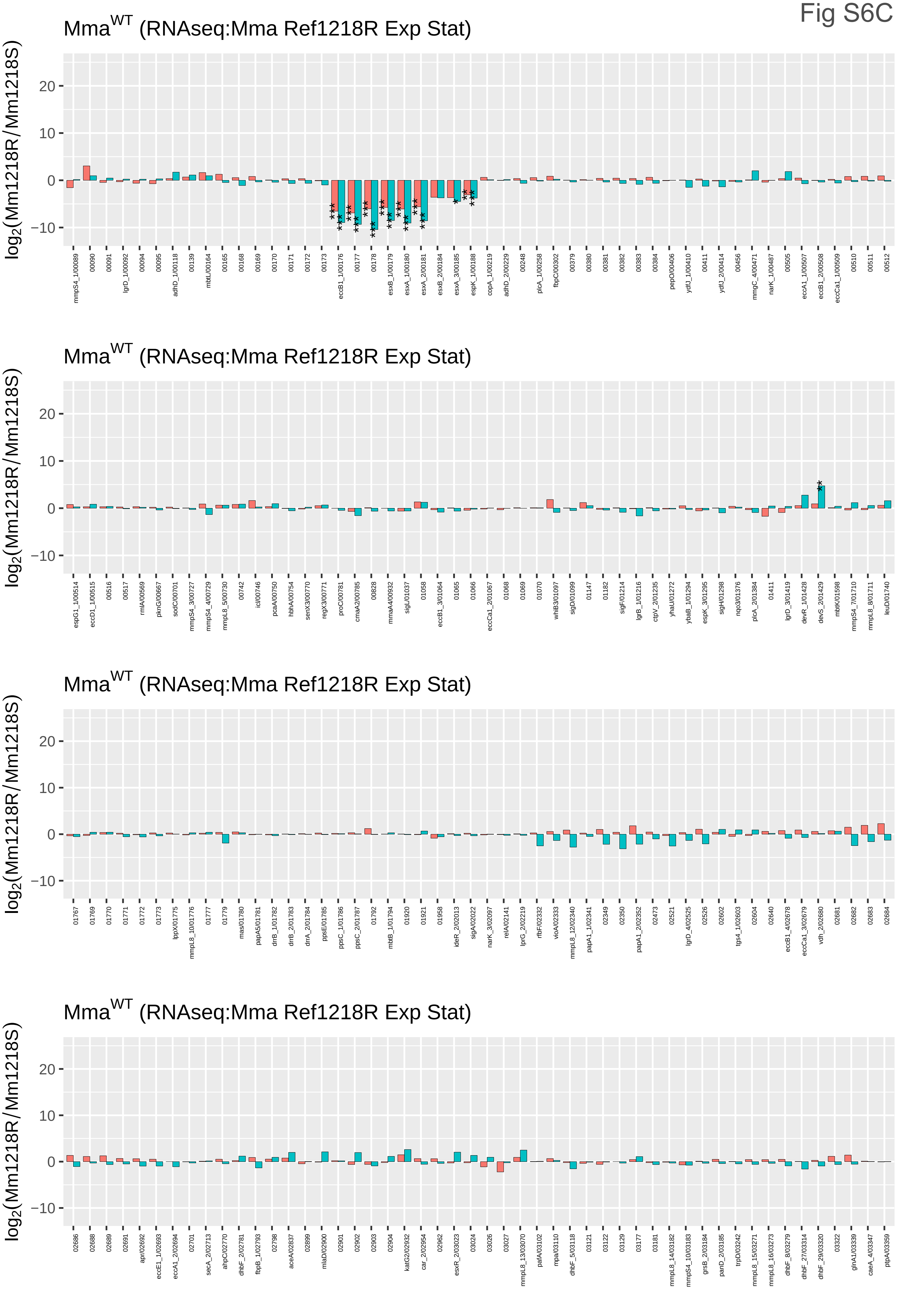

MmaWT (RNAseq:Mma Ref1218R Exp Stat)
log2(Mm1218R/Mm1218S)
20
10
0
−10
MmaWT (RNAseq:Mma Ref1218R Exp Stat)
log2(Mm1218R/Mm1218S)
MmaWT (RNAseq:Mma Ref1218R Exp Stat)
log2(Mm1218R/Mm1218S)
MmaWT (RNAseq:Mma Ref1218R Exp Stat)
log2(Mm1218R/Mm1218S)

Fig 6C

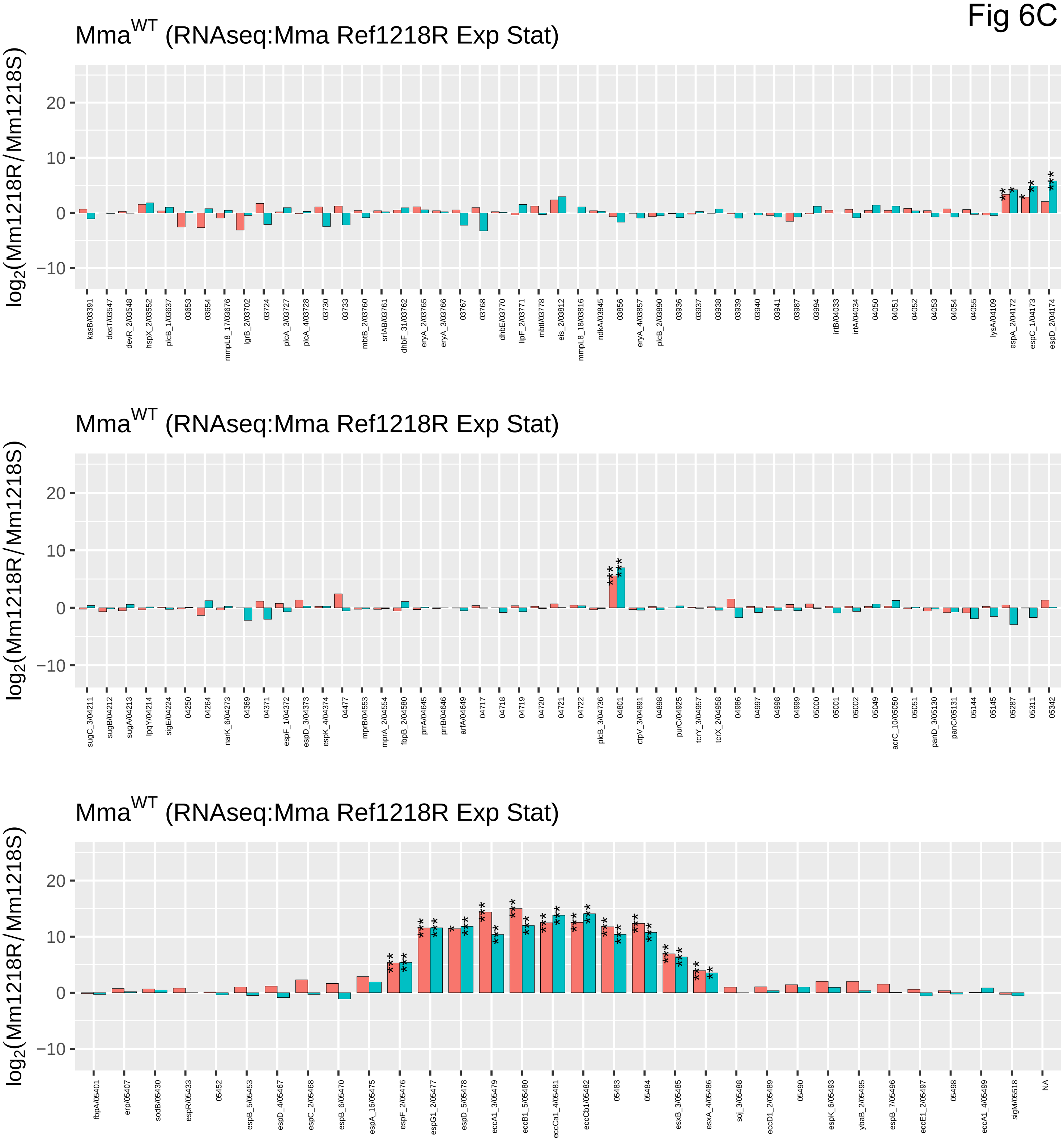

Mma^WT (RNAseq:Mma Ref1218R Exp Stat)
log2(Mm1218R/Mm1218S)
20
10
0
−10
kasB/03391
dosT/03547
devR_2/03548
hspX_2/03552
plcB_1/03637
03653
03654
mmpL8_17/03676
lgrB_2/03702
03724
plcA_3/03727
plcA_4/03728
03730
03733
mbtB_2/03760
srfAB/03761
dhbF_31/03762
eryA_2/03765
eryA_3/03766
03767
03768
dhbE/03770
lipF_2/03771
mbtI/03778
eis_2/03812
mmpL8_18/03816
ndkA/03845
03856
eryA_4/03857
plcB_2/03890
03936
03937
03938
03939
03940
03941
03987
03994
irtB/04033
irtA/04034
04050
04051
04052
04053
04054
04055
lysA/04109
espA_2/04172
espC_1/04173
espD_2/04174
Mma^WT (RNAseq:Mma Ref1218R Exp Stat)
log2(Mm1218R/Mm1218S)
20
10
0
−10
sugC_3/04211
sugB/04212
sugA/04213
lpqY/04214
sigE/04224
04250
04264
narK_6/04273
04369
04371
espF_1/04372
espD_3/04373
espK_4/04374
04477
mprB/04553
mprA_2/04554
fbpB_2/04580
prrA/04645
prrB/04646
arfA/04649
04717
04718
04719
04720
04721
04722
plcB_3/04736
04801
ctpV_3/04891
04898
purC/04925
tcrY_3/04957
tcrX_2/04958
04986
04997
04998
04999
05000
05001
05002
05049
acrC_10/05050
05051
panD_3/05130
panC/05131
05144
05145
05287
05311
05342
Mma^WT (RNAseq:Mma Ref1218R Exp Stat)
log2(Mm1218R/Mm1218S)
20
10
0
−10
fbpA/05401
erp/05407
sodB/05430
espR/05433
05452
espB_5/05453
espD_4/05467
espC_2/05468
espB_6/05470
espA_16/05475
espF_2/05476
espG1_2/05477
espD_5/05478
eccA1_3/05479
eccB1_5/05480
eccCa1_4/05481
eccCb1/05482
05483
05484
esxB_3/05485
esxA_4/05486
soj_3/05488
eccD1_2/05489
05490
espK_6/05493
ybaB_2/05495
espB_7/05496
eccE1_2/05497
05498
eccA1_4/05499
sigM/05518
NA

Fig S6D

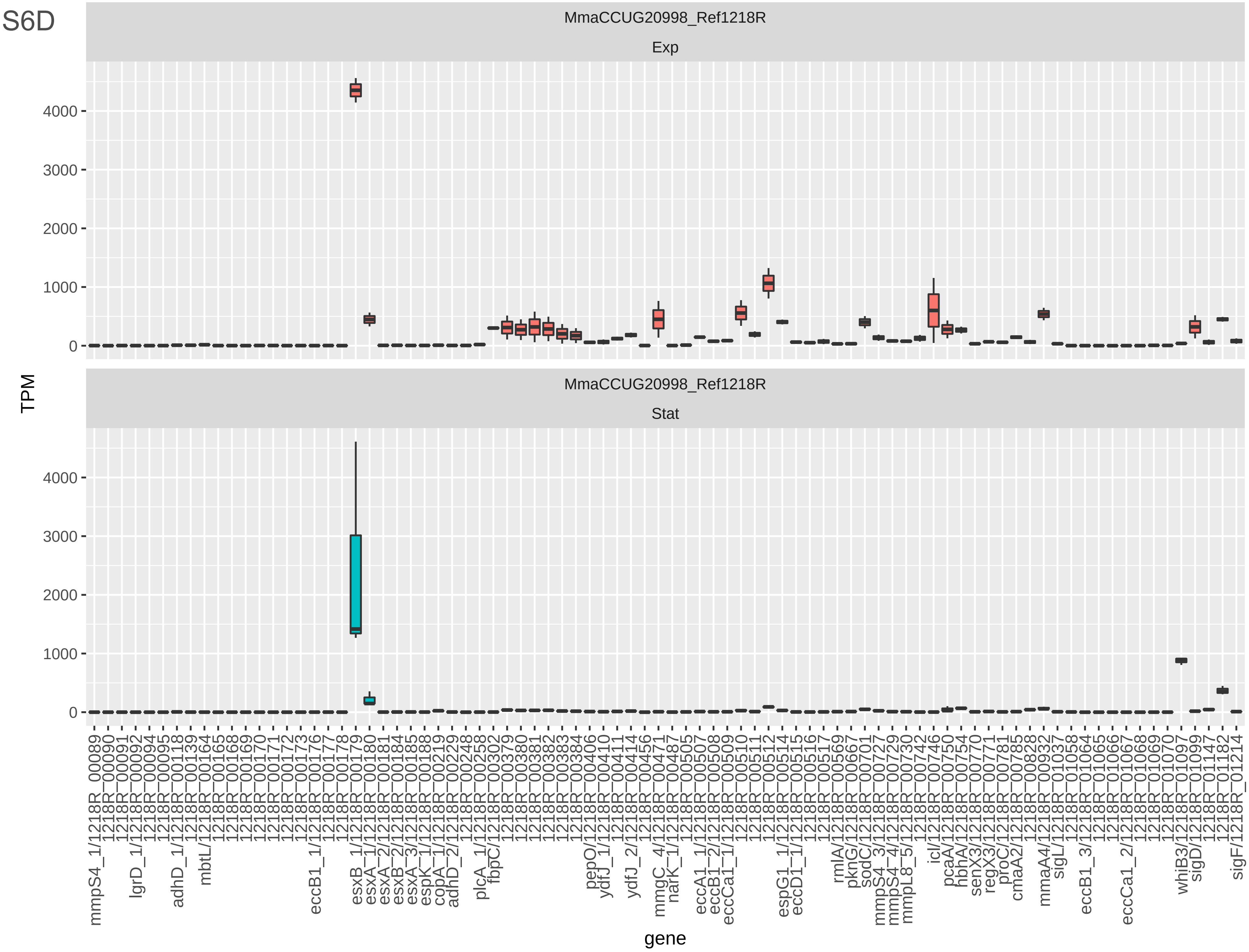
MmaCCUG20998_Ref1218R
Exp
MmaCCUG20998_Ref1218R
Stat
TPM
0
1000
2000
3000
4000
gene
mmpS4_1/1218R_00089
1218R_00090
1218R_00091
lgrD_1/1218R_00092
1218R_00094
1218R_00095
adhD_1/1218R_00118
1218R_00139
mbtL/1218R_00164
1218R_00165
1218R_00168
1218R_00169
1218R_00170
1218R_00171
1218R_00172
1218R_00173
eccB1_1/1218R_00176
1218R_00177
1218R_00178
esxB_1/1218R_00179
esxA_1/1218R_00180
esxA_2/1218R_00181
esxB_2/1218R_00184
esxA_3/1218R_00185
espK_1/1218R_00188
copA_1/1218R_00219
adhD_2/1218R_00229
1218R_00248
plcA_1/1218R_00258
fbpC/1218R_00302
1218R_00379
1218R_00380
1218R_00381
1218R_00382
1218R_00383
1218R_00384
pepO/1218R_00406
ydfJ_1/1218R_00410
1218R_00411
ydfJ_2/1218R_00414
1218R_00456
mmgC_4/1218R_00471
narK_1/1218R_00487
1218R_00505
eccA1_1/1218R_00507
eccB1_2/1218R_00508
eccCa1_1/1218R_00509
1218R_00510
1218R_00511
1218R_00512
espG1_1/1218R_00514
eccD1_1/1218R_00515
1218R_00516
1218R_00517
rmlA/1218R_00569
pknG/1218R_00667
sodC/1218R_00701
mmpS4_3/1218R_00727
mmpS4_4/1218R_00729
mmpL8_5/1218R_00730
1218R_00742
icl/1218R_00746
pcaA/1218R_00750
hbhA/1218R_00754
senX3/1218R_00770
regX3/1218R_00771
proC/1218R_00781
cmaA2/1218R_00785
1218R_00828
mmaA4/1218R_00932
sigL/1218R_01037
1218R_01058
eccB1_3/1218R_01064
1218R_01065
1218R_01066
eccCa1_2/1218R_01067
1218R_01068
1218R_01069
1218R_01070
whiB3/1218R_01097
sigD/1218R_01099
1218R_01147
1218R_01182
sigF/1218R_01214

Fig S6D

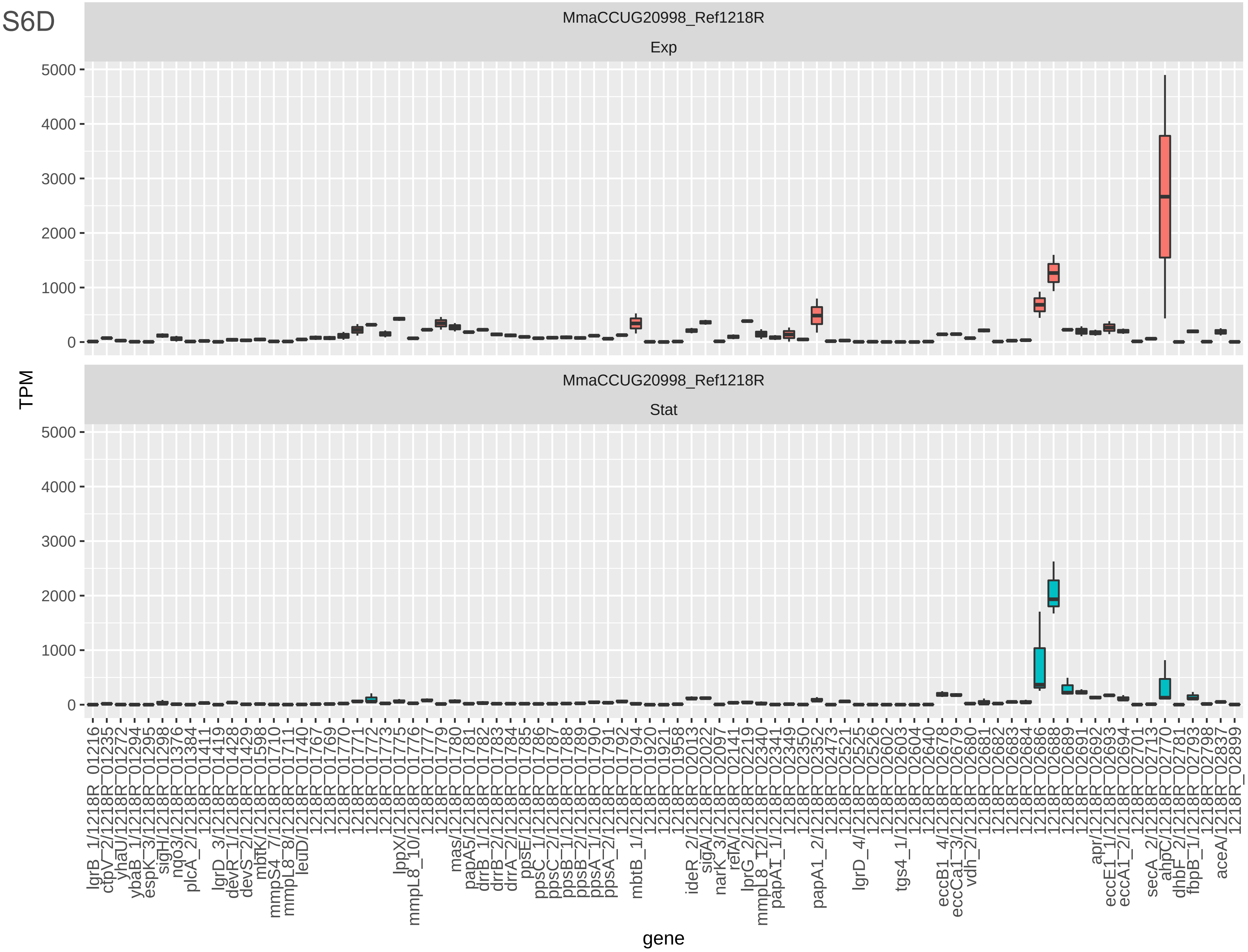

MmaCCUG20998_Ref1218R
Exp
MmaCCUG20998_Ref1218R
Stat
5000
4000
3000
2000
1000
0
TPM
gene

Fig S6D

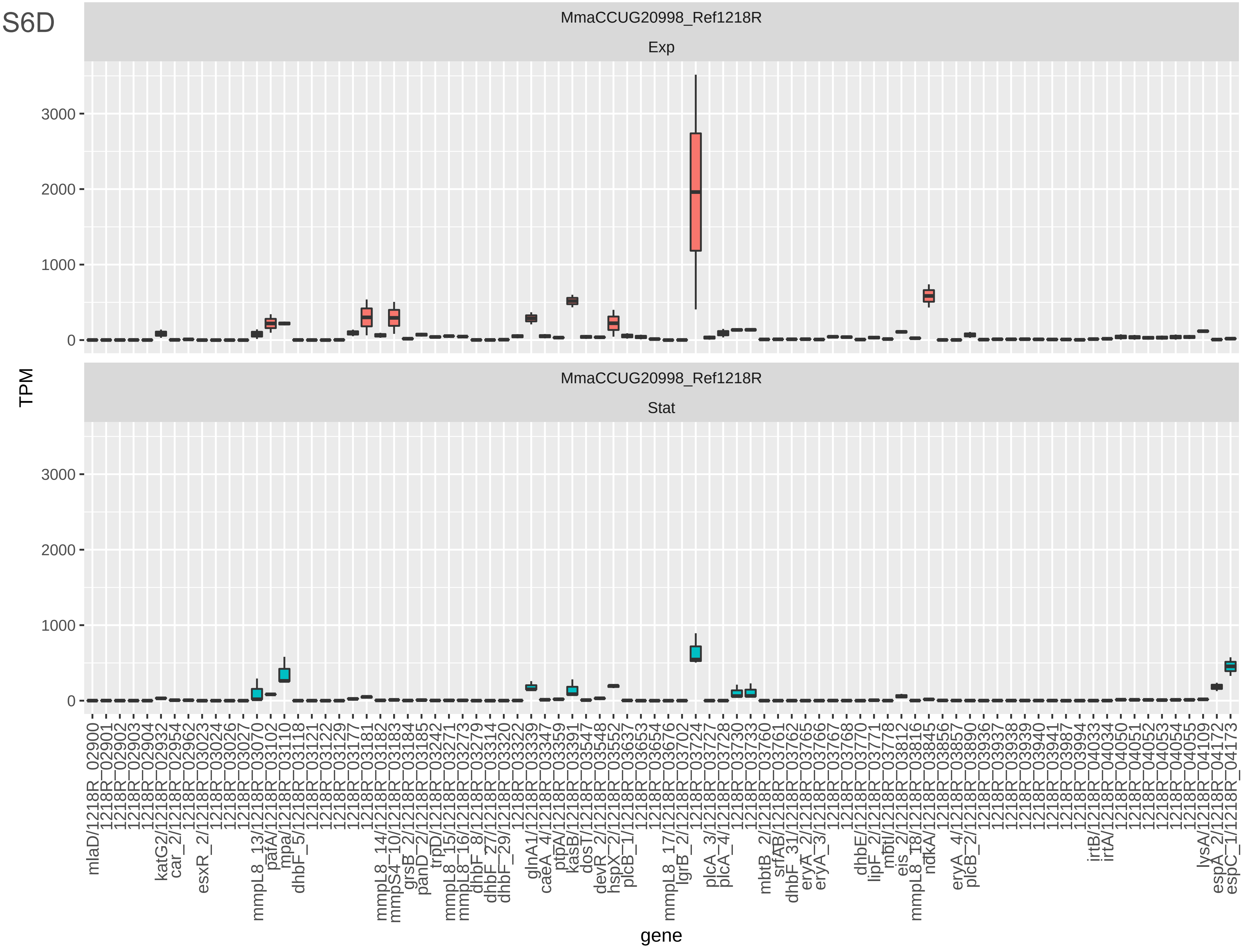

MmaCCUG20998_Ref1218R
Exp
MmaCCUG20998_Ref1218R
Stat
TPM
0
1000
2000
3000
gene

Fig S6D

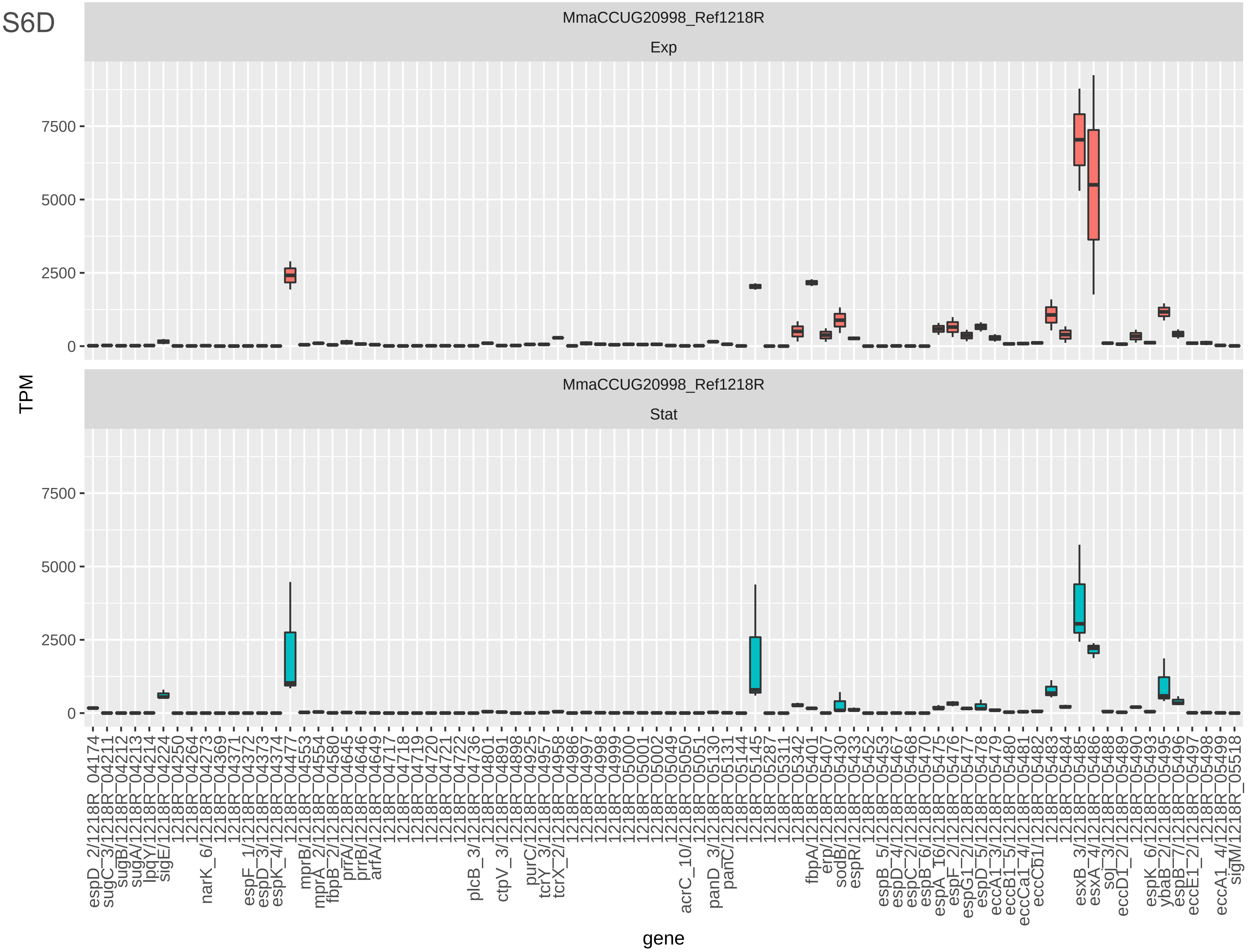
MmaCCUG20998_Ref1218R
Exp
MmaCCUG20998_Ref1218R
Stat
TPM
7500
5000
2500
0
7500
5000
2500
0
gene
espD_2/1218R_04174
sugC_3/1218R_04211
sugB/1218R_04212
sugA/1218R_04213
lpqY/1218R_04214
sigE/1218R_04224
1218R_04250
1218R_04264
narK_6/1218R_04273
1218R_04369
1218R_04371
espF_1/1218R_04372
espD_3/1218R_04373
espK_4/1218R_04374
1218R_04477
mprB/1218R_04553
mprA_2/1218R_04554
fbpB_2/1218R_04580
prrA/1218R_04645
prrB/1218R_04646
arfA/1218R_04649
1218R_04717
1218R_04718
1218R_04719
1218R_04720
1218R_04721
1218R_04722
plcB_3/1218R_04736
1218R_04801
ctpV_3/1218R_04891
1218R_04898
purC/1218R_04925
tcrY_3/1218R_04957
tcrX_2/1218R_04958
1218R_04986
1218R_04997
1218R_04998
1218R_04999
1218R_05000
1218R_05001
1218R_05002
1218R_05049
acrC_10/1218R_05050
1218R_05051
panD_3/1218R_05130
panC/1218R_05131
1218R_05144
1218R_05145
1218R_05287
1218R_05311
1218R_05342
fbpA/1218R_05401
erp/1218R_05407
sodB/1218R_05430
espR/1218R_05433
1218R_05452
espB_5/1218R_05453
espD_4/1218R_05467
espC_2/1218R_05468
espB_6/1218R_05470
espA_T6/1218R_05475
espF_2/1218R_05476
espG1_2/1218R_05477
espD_5/1218R_05478
eccA1_3/1218R_05479
eccB1_5/1218R_05480
eccCa1_4/1218R_05481
eccCb1/1218R_05482
1218R_05483
1218R_05484
esxB_3/1218R_05485
esxA_4/1218R_05486
soj_3/1218R_05488
eccD1_2/1218R_05489
1218R_05490
espK_6/1218R_05493
ybaB_2/1218R_05495
espB_7/1218R_05496
eccE1_2/1218R_05497
1218R_05498
eccA1_4/1218R_05499
sigM/1218R_05518

Fig S6E

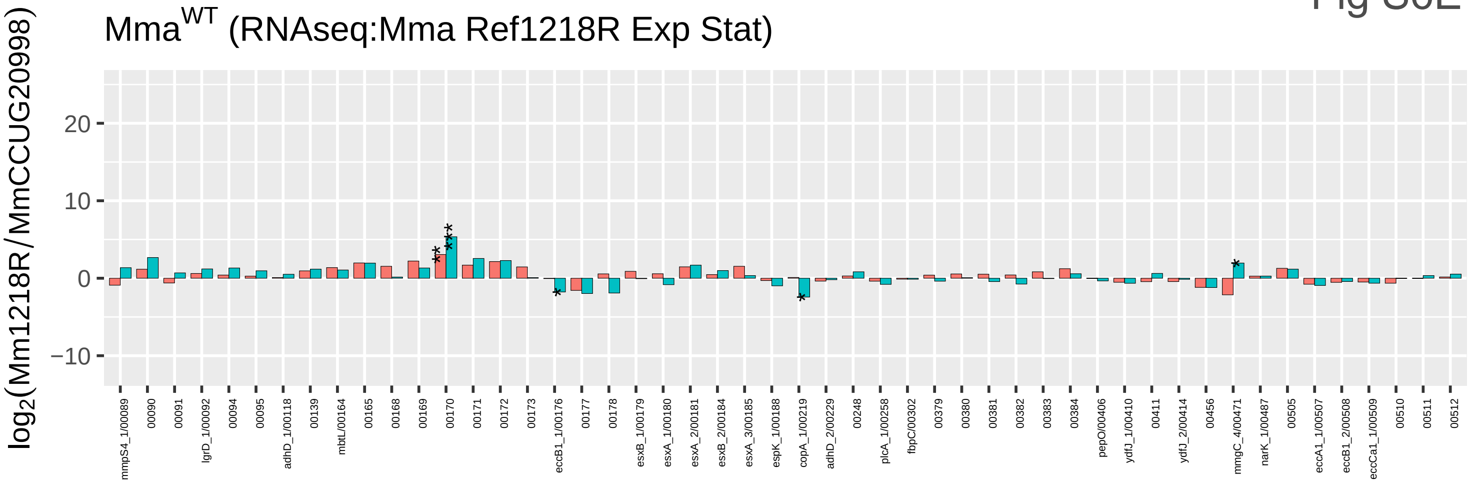
Mma^WT (RNAseq:Mma Ref1218R Exp Stat)
log2(Mm1218R/MmCCUG20998)
20
10
0
−10

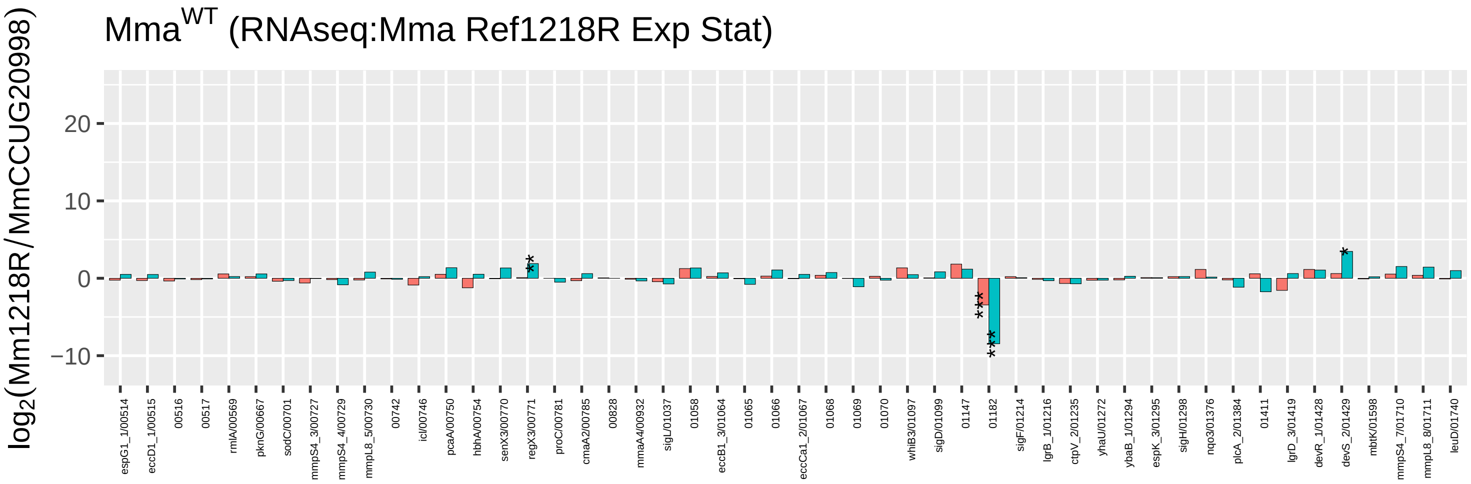
Mma^WT (RNAseq:Mma Ref1218R Exp Stat)
log2(Mm1218R/MmCCUG20998)
20
10
0
−10

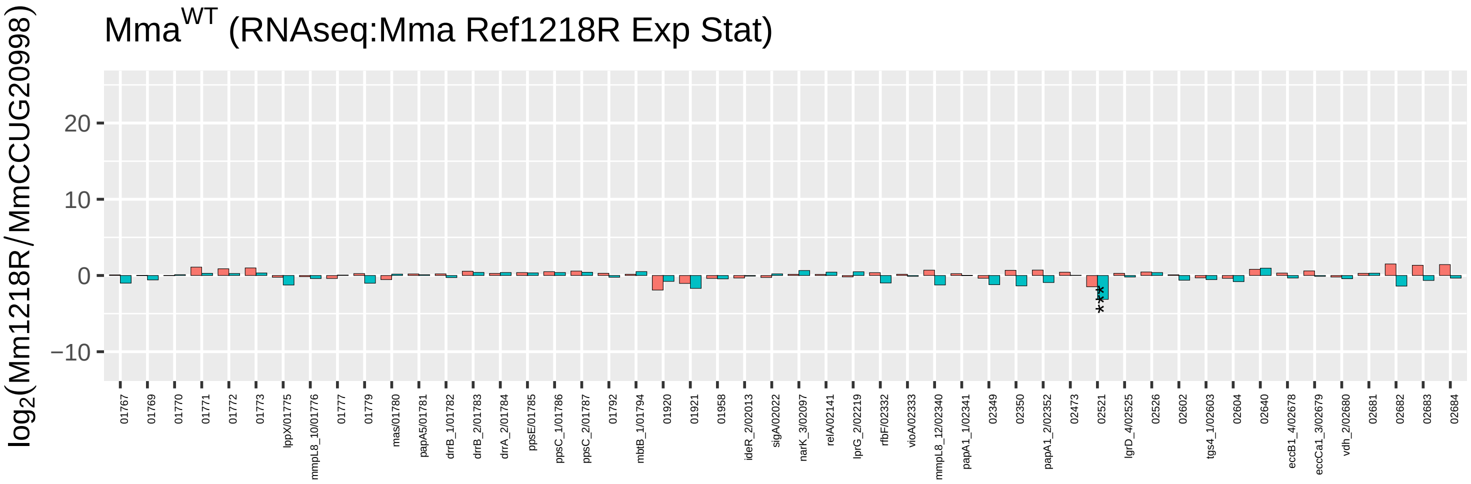
Mma^WT (RNAseq:Mma Ref1218R Exp Stat)
log2(Mm1218R/MmCCUG20998)
20
10
0
−10

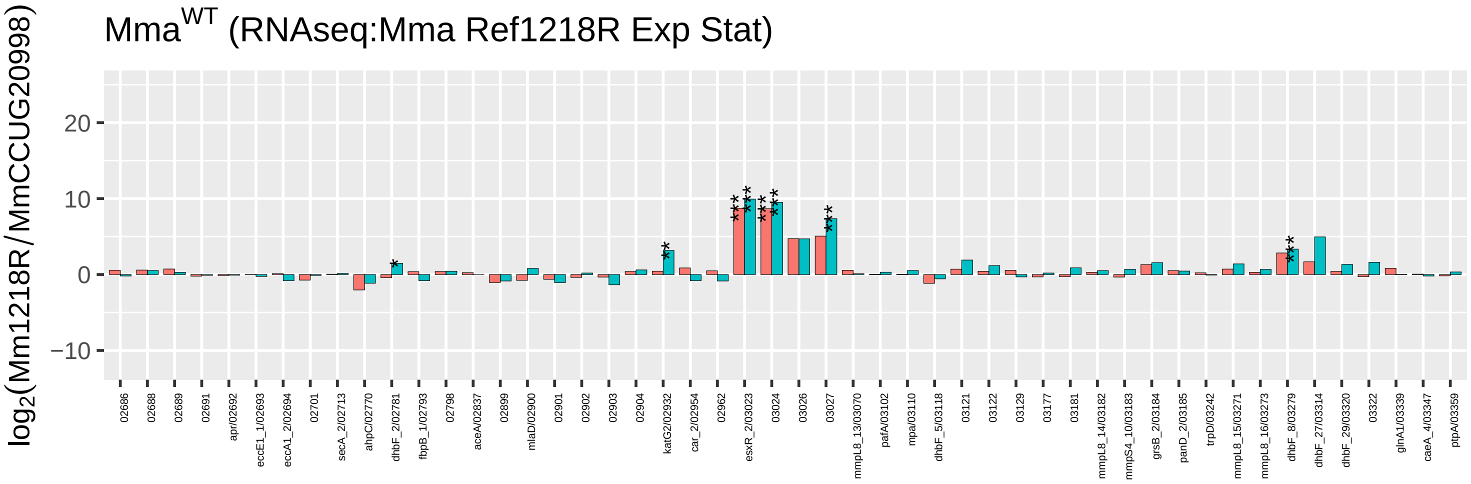
Mma^WT (RNAseq:Mma Ref1218R Exp Stat)
log2(Mm1218R/MmCCUG20998)
20
10
0
−10

Fig S6E

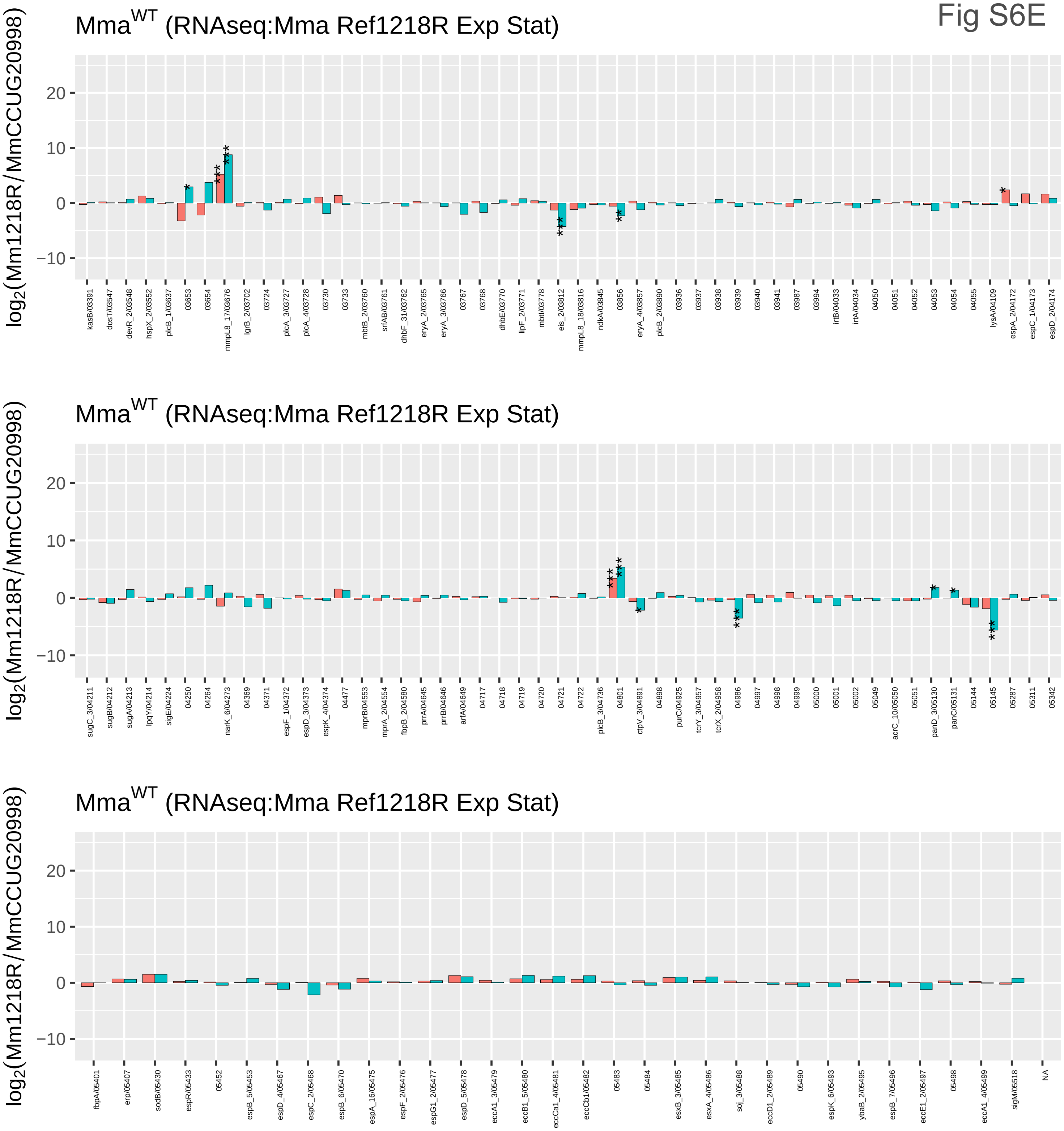

Mma^WT (RNAseq:Mma Ref1218R Exp Stat)
log2(Mm1218R/MmCCUG20998)
20
10
0
−10
kasB/03391
dosT/03547
devR_2/03548
hspX_2/03552
plcB_1/03637
03653
03654
mmpL8_17/03676
lgrB_2/03702
03724
plcA_3/03727
plcA_4/03728
03730
03733
mbtB_2/03760
srfAB/03761
dhbF_31/03762
eryA_2/03765
eryA_3/03766
03767
03768
dhbE/03770
lipF_2/03771
mbtI/03778
eis_2/03812
mmpL8_18/03816
ndkA/03845
03856
eryA_4/03857
plcB_2/03890
03936
03937
03938
03939
03940
03941
03987
03994
irtB/04033
irtA/04034
04050
04051
04052
04053
04054
04055
lysA/04109
espA_2/04172
espC_1/04173
espD_2/04174
Mma^WT (RNAseq:Mma Ref1218R Exp Stat)
log2(Mm1218R/MmCCUG20998)
20
10
0
−10
sugC_3/04211
sugB/04212
sugA/04213
lpqY/04214
sigE/04224
04250
04264
narK_6/04273
04369
04371
espF_1/04372
espD_3/04373
espK_4/04374
04477
mprB/04553
mprA_2/04554
fbpB_2/04580
prrA/04645
prrB/04646
arfA/04649
04717
04718
04719
04720
04721
04722
plcB_3/04736
04801
ctpV_3/04891
04898
purC/04925
tcrY_3/04957
tcrX_2/04958
04986
04997
04998
04999
05000
05001
05002
05049
acrC_10/05050
05051
panD_3/05130
panC/05131
05144
05145
05287
05311
05342
Mma^WT (RNAseq:Mma Ref1218R Exp Stat)
log2(Mm1218R/MmCCUG20998)
20
10
0
−10
fbpA/05401
erp/05407
sodB/05430
espR/05433
05452
espB_5/05453
espD_4/05467
espC_2/05468
espB_6/05470
espA_16/05475
espF_2/05476
espG1_2/05477
espD_5/05478
eccA1_3/05479
eccB1_5/05480
eccCa1_4/05481
eccCb1/05482
05483
05484
esxB_3/05485
esxA_4/05486
soj_3/05488
eccD1_2/05489
05490
espK_6/05493
ybaB_2/05495
espB_7/05496
eccE1_2/05497
05498
eccA1_4/05499
sigM/05518
NA

Fig S6F

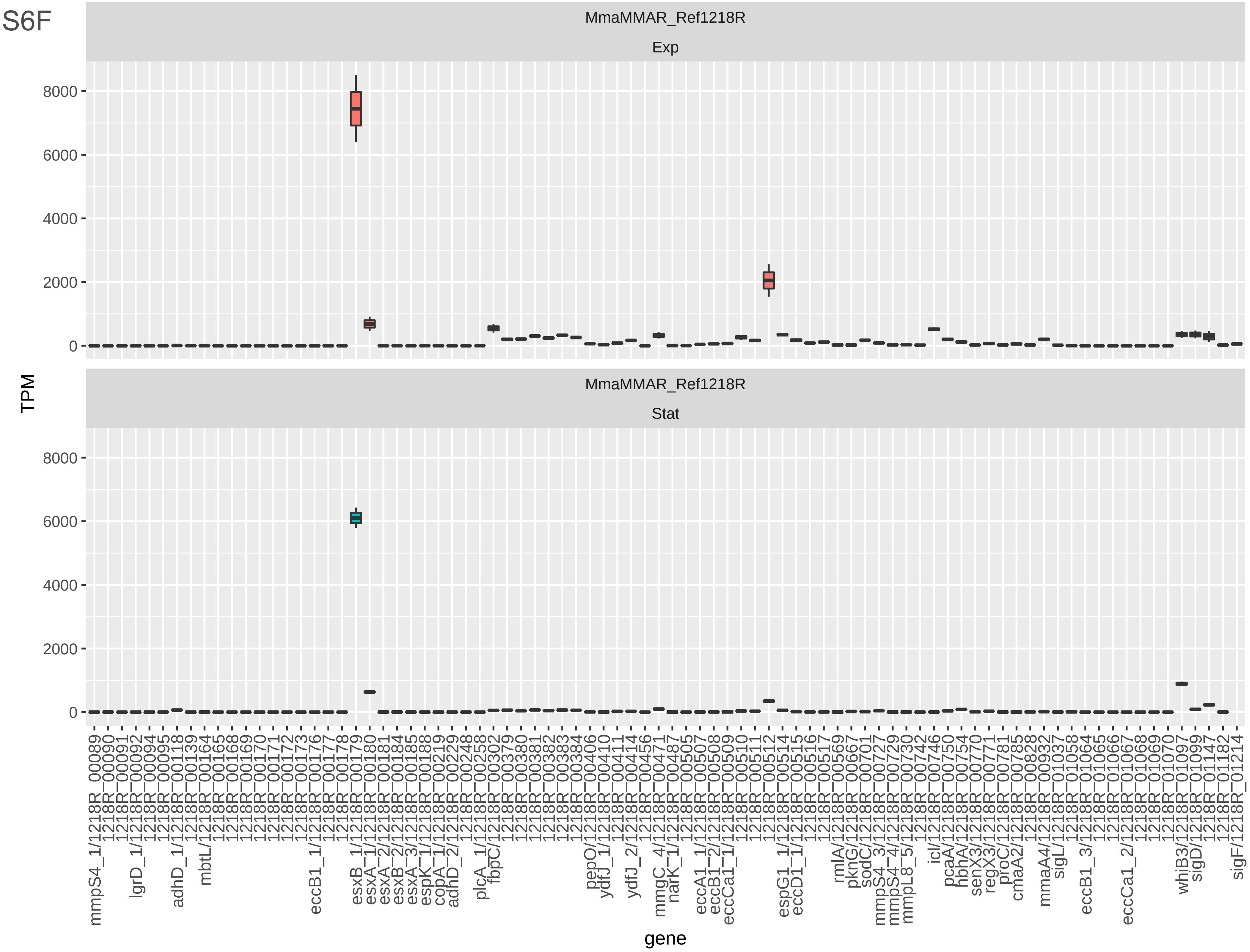

MmaMMAR_Ref1218R
Exp
MmaMMAR_Ref1218R
Stat
TPM
gene
0
2000
4000
6000
8000

Fig S6F

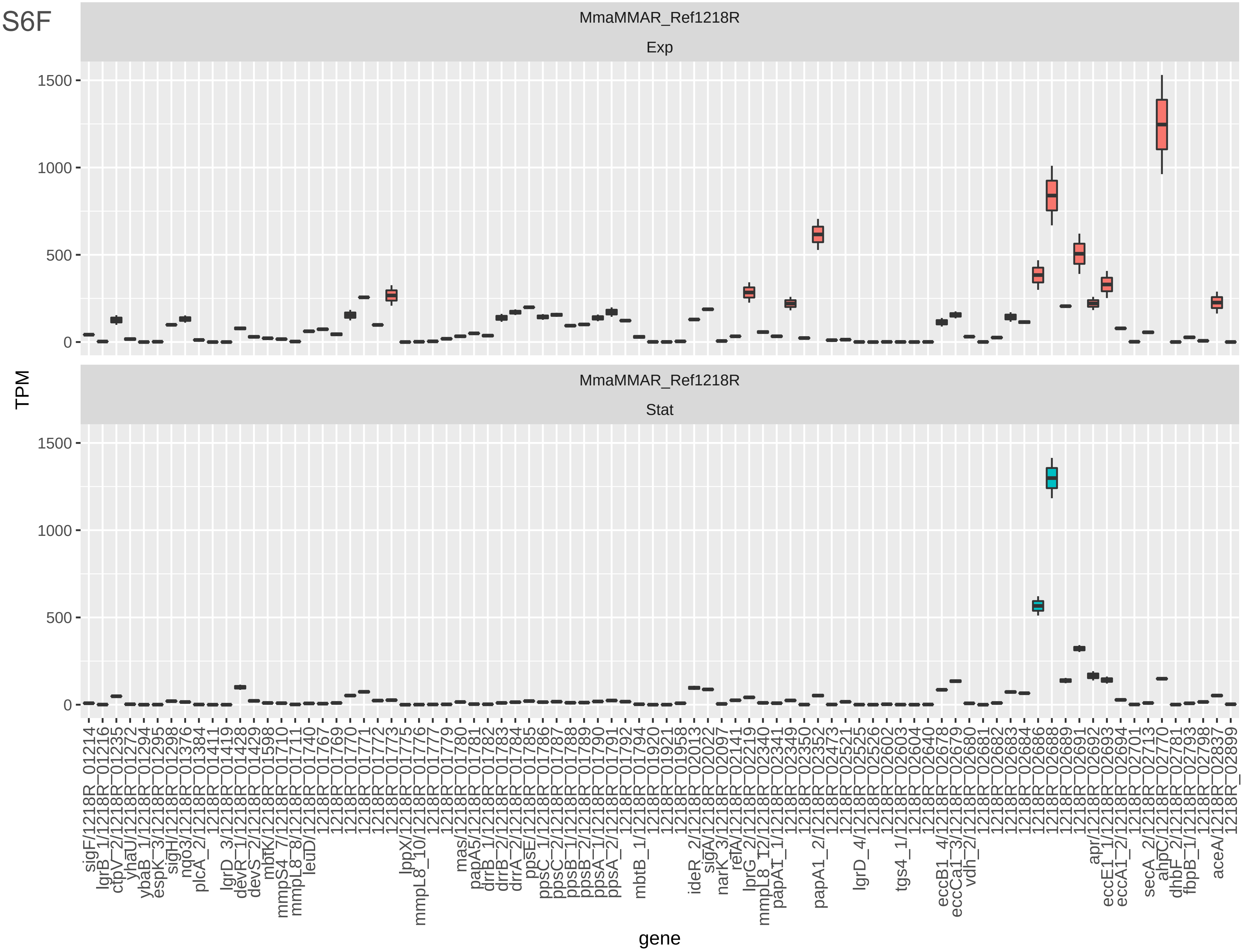

Fig S6F

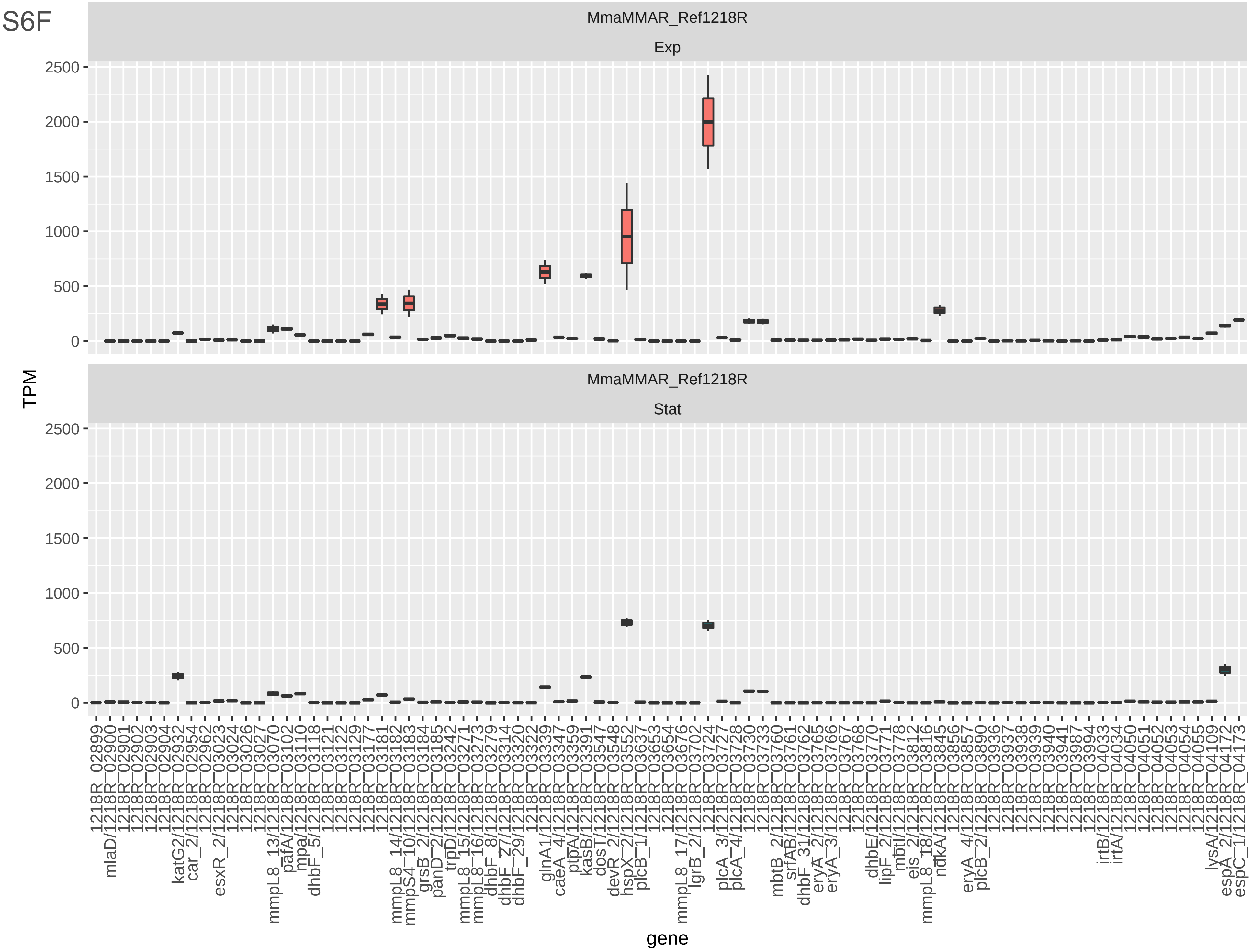
MmaMMAR_Ref1218R
Exp
MmaMMAR_Ref1218R
Stat
TPM
2500
2000
1500
1000
500
0
gene
1218R_02899
mlaD/1218R_02900
1218R_02901
1218R_02902
1218R_02903
1218R_02904
katG2/1218R_02932
car_2/1218R_02954
1218R_02962
esxR_2/1218R_03023
1218R_03024
1218R_03026
1218R_03027
mmpL8_13/1218R_03070
pafA/1218R_03102
mpa/1218R_03110
dhbF_5/1218R_03118
1218R_03121
1218R_03122
1218R_03129
1218R_03177
1218R_03181
mmpL8_14/1218R_03182
mmpS4_10/1218R_03183
grsB_2/1218R_03184
panD_2/1218R_03185
trpD/1218R_03242
mmpL8_15/1218R_03271
mmpL8_16/1218R_03273
dhbF_8/1218R_03279
dhbF_27/1218R_03314
dhbF_29/1218R_03320
1218R_03322
glnA1/1218R_03339
caeA_4/1218R_03347
ptpA/1218R_03359
kasB/1218R_03391
dosT/1218R_03547
devR_2/1218R_03548
hspX_2/1218R_03552
plcB_1/1218R_03637
1218R_03653
1218R_03654
mmpL8_17/1218R_03676
lgrB_2/1218R_03702
1218R_03724
plcA_3/1218R_03727
plcA_4/1218R_03728
1218R_03730
1218R_03733
mbtB_2/1218R_03760
srfAB/1218R_03761
dhbF_31/1218R_03762
eryA_2/1218R_03765
eryA_3/1218R_03766
1218R_03767
1218R_03768
dhbE/1218R_03770
lipF_2/1218R_03771
mbtI/1218R_03778
eis_2/1218R_03812
mmpL8_18/1218R_03816
ndkA/1218R_03845
1218R_03856
eryA_4/1218R_03857
plcB_2/1218R_03890
1218R_03936
1218R_03937
1218R_03938
1218R_03939
1218R_03940
1218R_03941
1218R_03987
1218R_03994
irtB/1218R_04033
irtA/1218R_04034
1218R_04050
1218R_04051
1218R_04052
1218R_04053
1218R_04054
1218R_04055
lysA/1218R_04109
espA_2/1218R_04172
espC_1/1218R_04173

Fig S6F

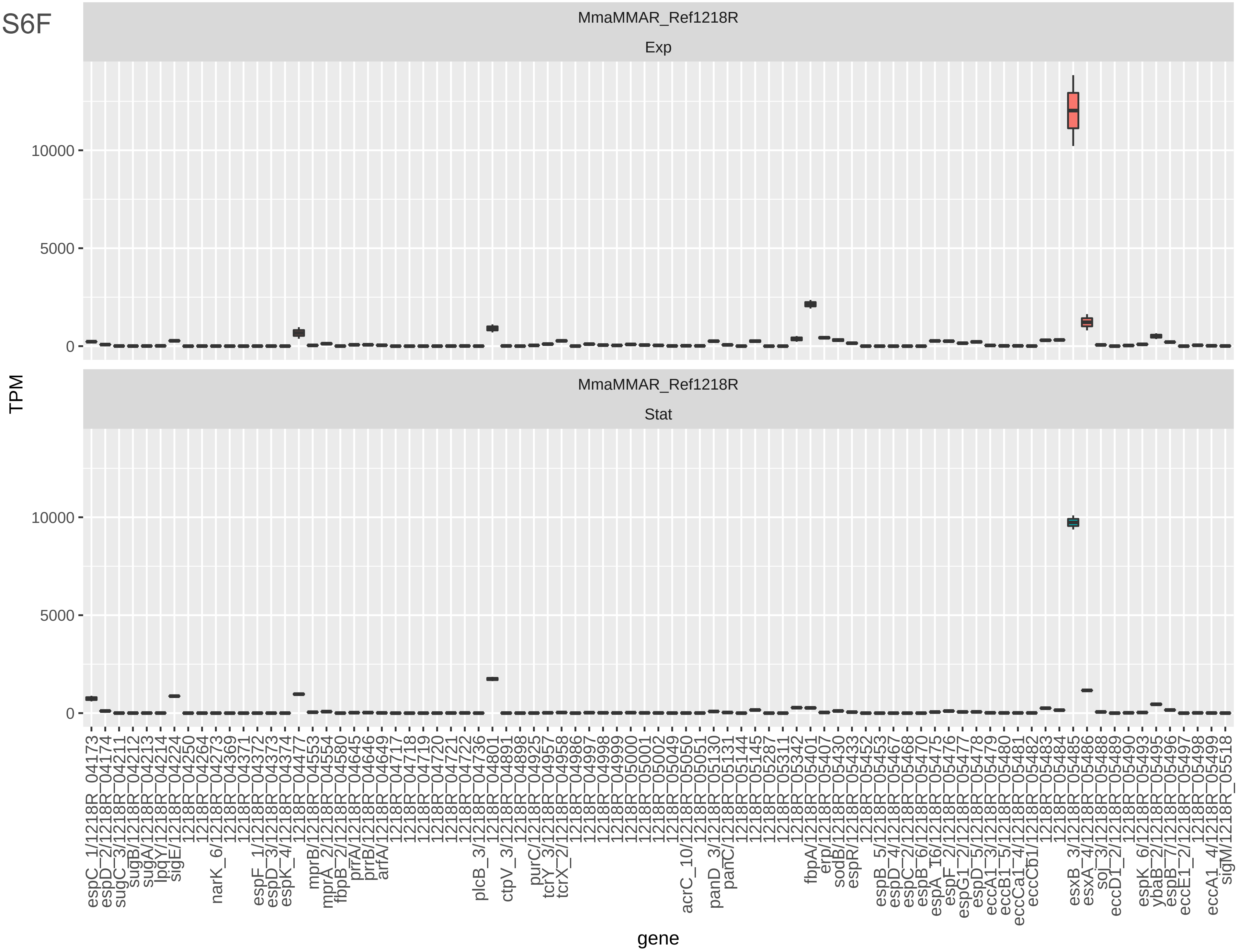
MmaMMAR_Ref1218R
Exp
MmaMMAR_Ref1218R
Stat
TPM
gene
10000
5000
0

Fig S6G

Mma$^{WT}$ (RNAseq:Mma Ref1218R Exp Stat)

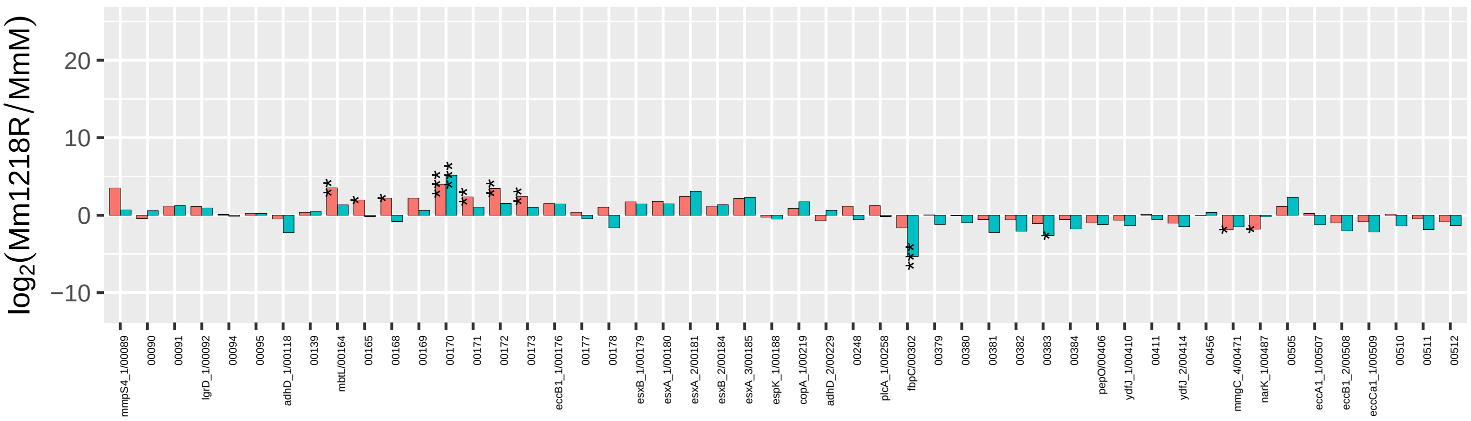


Mma$^{WT}$ (RNAseq:Mma Ref1218R Exp Stat)

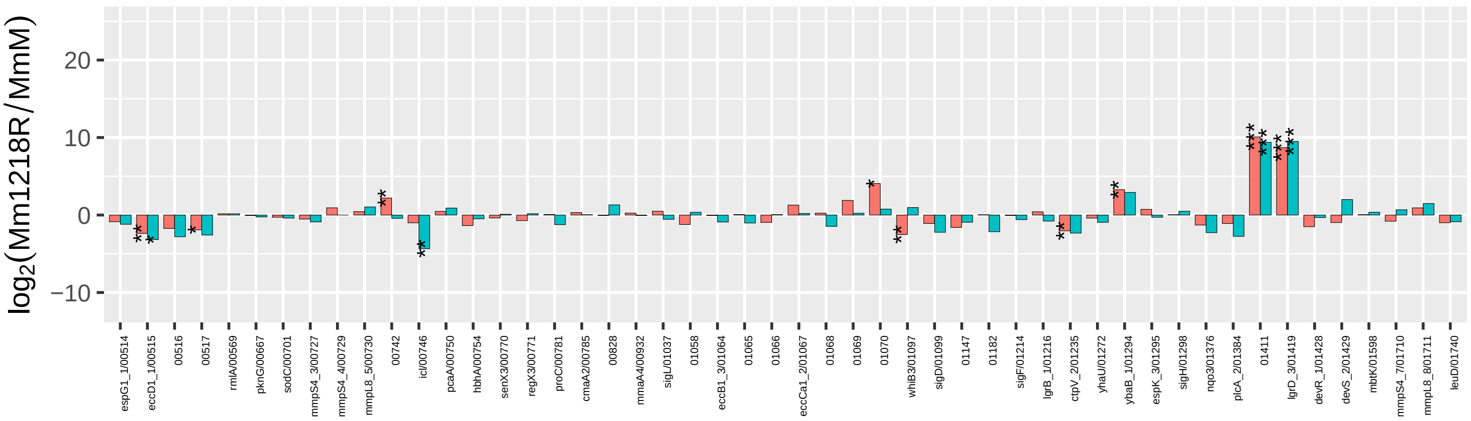


Mma$^{WT}$ (RNAseq:Mma Ref1218R Exp Stat)

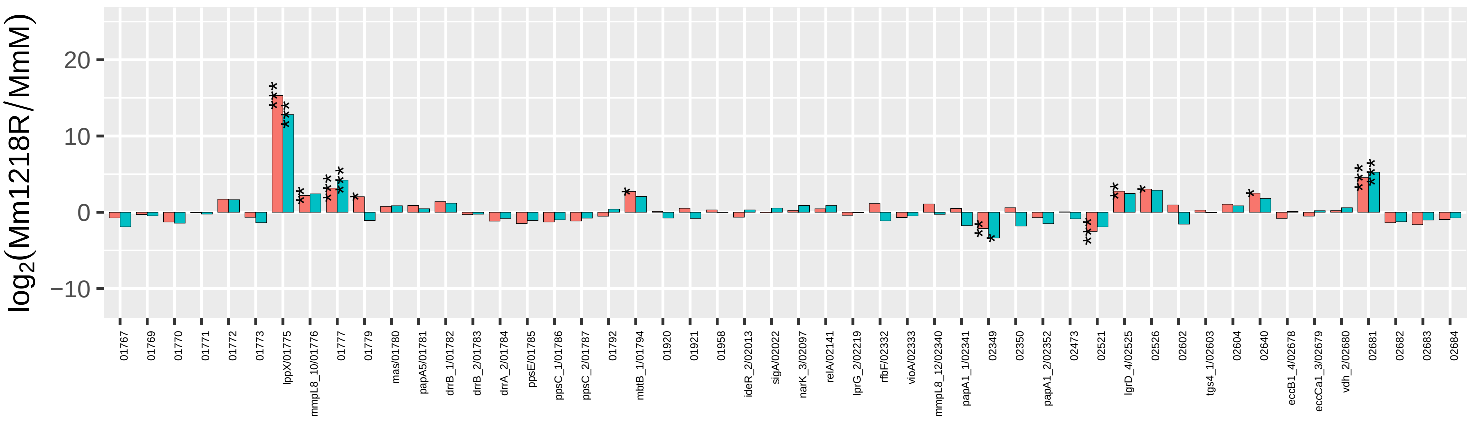


Mma$^{WT}$ (RNAseq:Mma Ref1218R Exp Stat)

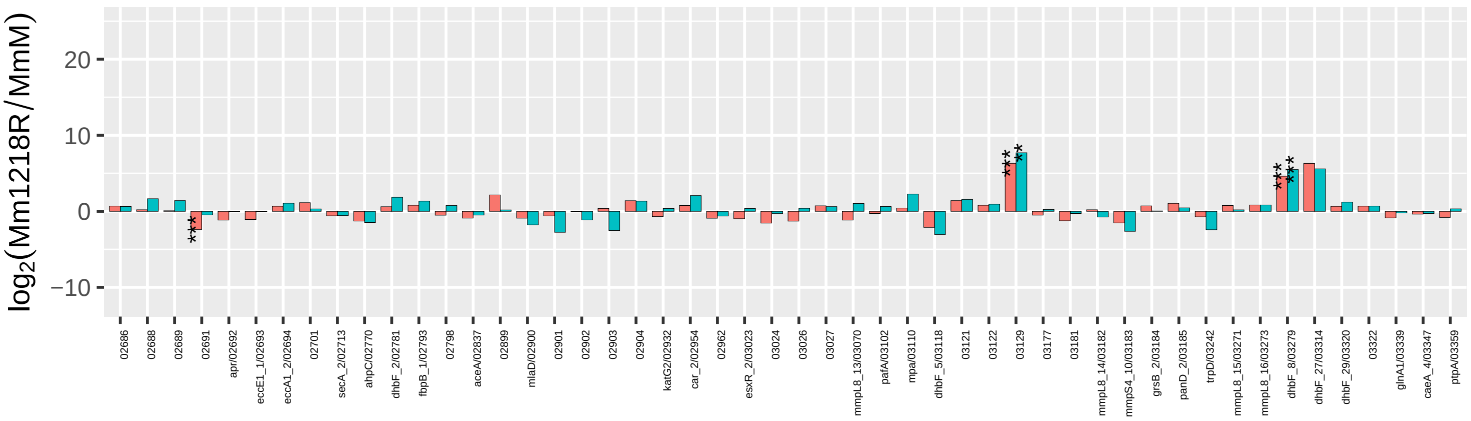

Fig S6G

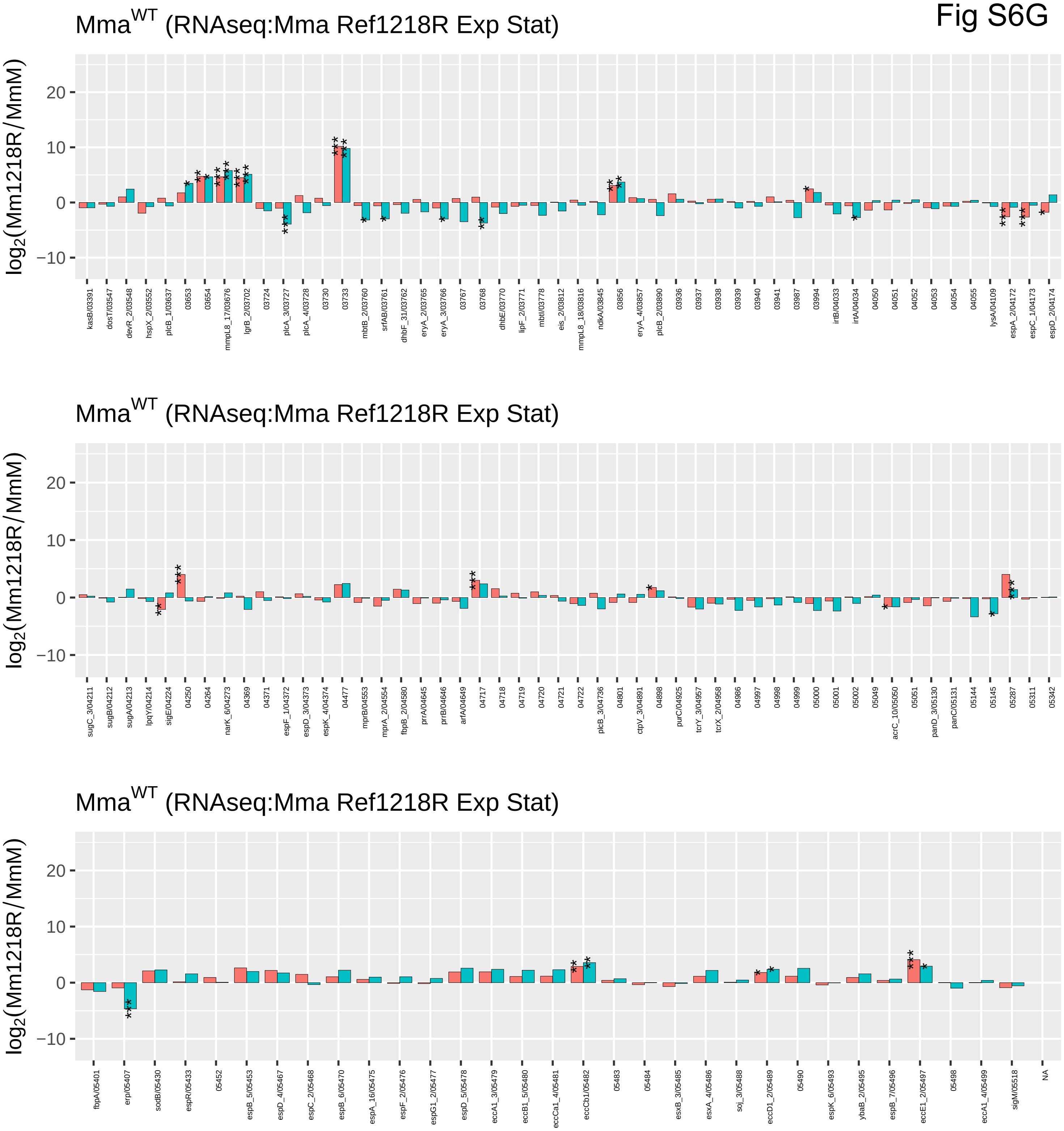

Mma^WT (RNAseq:Mma Ref1218R Exp Stat)
log2(Mm1218R/MmM)
Mma^WT (RNAseq:Mma Ref1218R Exp Stat)
log2(Mm1218R/MmM)
Mma^WT (RNAseq:Mma Ref1218R Exp Stat)
log2(Mm1218R/MmM)

**Figure S7** Bar plots showing transcription of ESX-1, ESX-3, ESX-4, ESX-5 and ESX-6 genes in 1218R, 1218S, *Mmar*$^{\mathrm{CCUG}}$ and *Mmar*$^{\mathrm{M}}$.

(a) Transcript levels (distribution) in exponentially growing 1218R cells.

(b) Transcript levels (distribution) in stationary 1218R cells.

(c) Change in transcript levels, expressed as $\log_2$-fold change, comparing levels in exponentially growing and stationary 1218R cells. A negative $\log_2$-value suggests that the corresponding mRNA is more abundant in exponentially growing cells while a positive value suggests higher levels in stationary cells. Statistical significance, see Materials and Methods; *p-value $< 0.05$; **p-value $< 0.01$; ***p-value $< 0.001$.

(d) Transcript levels (distribution) in exponentially growing 1218S cells.

(e) Transcript levels (distribution) in stationary 1218S cells.

(f) Change in transcript levels, expressed as $\log_2$-fold change, comparing levels in exponentially growing and stationary 1218S cells. A negative $\log_2$-value suggests that the corresponding mRNA is more abundant in exponentially growing cells while a positive value suggests higher levels in stationary cells. Statistical significance, see Materials and Methods; *p-value $< 0.05$; **p-value $< 0.01$; ***p-value $< 0.001$.

(g) Transcript levels (distribution) in exponentially growing *Mmar*$^{\mathrm{CCUG}}$ cells.

(h) Transcript levels (distribution) in stationary *Mmar*$^{\mathrm{CCUG}}$ cells.

(i) Change in transcript levels, expressed as $\log_2$-fold change, comparing levels in exponentially growing and stationary *Mmar*$^{\mathrm{CCUG}}$ cells. A negative $\log_2$-value suggests that the corresponding mRNA is more abundant in exponentially growing cells while a positive value suggests higher levels in stationary cells. Statistical significance, see Materials and Methods; *p-value $< 0.05$; **p-value $< 0.01$; ***p-value $< 0.001$.

(j) Transcript levels (distribution) in exponentially growing *Mmar*$^{\mathrm{M}}$ cells.

(k) Transcript levels (distribution) in exponentially growing *Mmar*$^{\mathrm{M}}$ cells.

(l) Change in transcript levels, expressed as $log_2$-fold change, comparing levels in exponentially growing and stationary *Mmar*$^{M}$ cells. A negative $log_2$-value suggests that the corresponding mRNA is more abundant in exponentially growing cells while a positive value suggests higher levels in stationary cells.

Statistical significance, see Materials and Methods; *p-value < 0.05; **p-value < 0.01; ***p-value < 0.001.

### Figure S7A-C

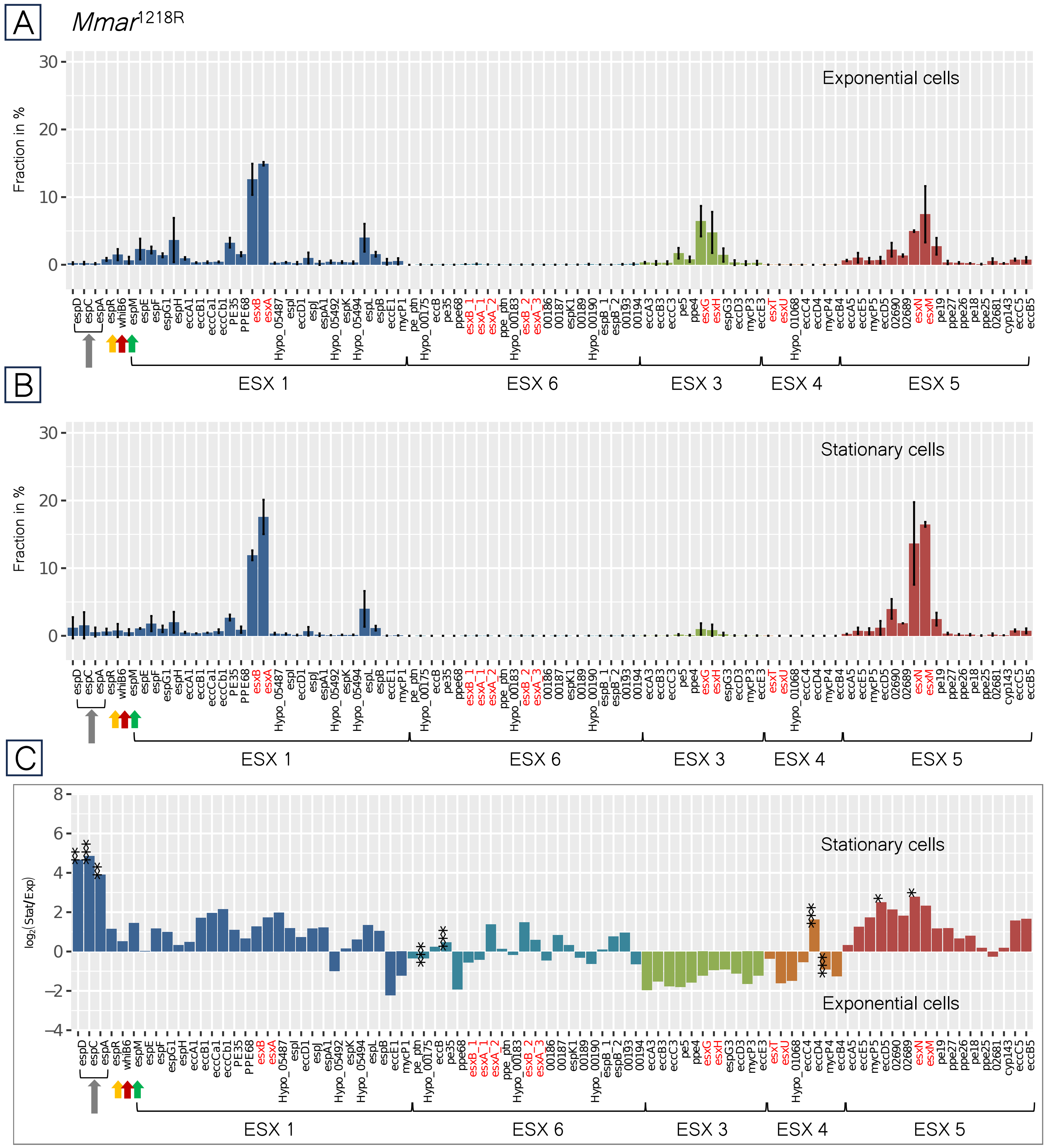

A
Mmar1218R
Exponential cells
Fraction in %
B
Stationary cells
Fraction in %
C
Stationary cells
Exponential cells
log2(Stat/Exp)
ESX 1
ESX 6
ESX 3
ESX 4
ESX 5
espD espC espA espR whiB6 espM espE espF espG1 espH eccA1 eccB1 eccCa1 eccCb1 PE35 PPE68 esxB esxA Hypo_05487 espI eccD1 espJ espA1 Hypo_05492 espK Hypo_05494 espL espB eccE1 mycP1 pe_ptn Hypo_00175 eccB pe35 ppe68 esxB_1 esxA_1 esxA_2 ppe_ptn Hypo_00183 esxB_2 esxA_3 00186 00187 espK1 00189 Hypo_00190 espB_1 espB_2 00193 00194 eccA3 eccB3 eccC3 pe5 ppe4 esxG esxH espG3 eccD3 mycP3 eccE3 esxT esxU Hypo_01068 eccC4 eccD4 mycP4 eccB4 eccA5 eccE5 mycP5 eccD5 02690 02689 esxN esxM pe19 ppe27 ppe26 pe18 ppe25 02681 cyp143 eccC5 eccB5

### Figure S7D-F

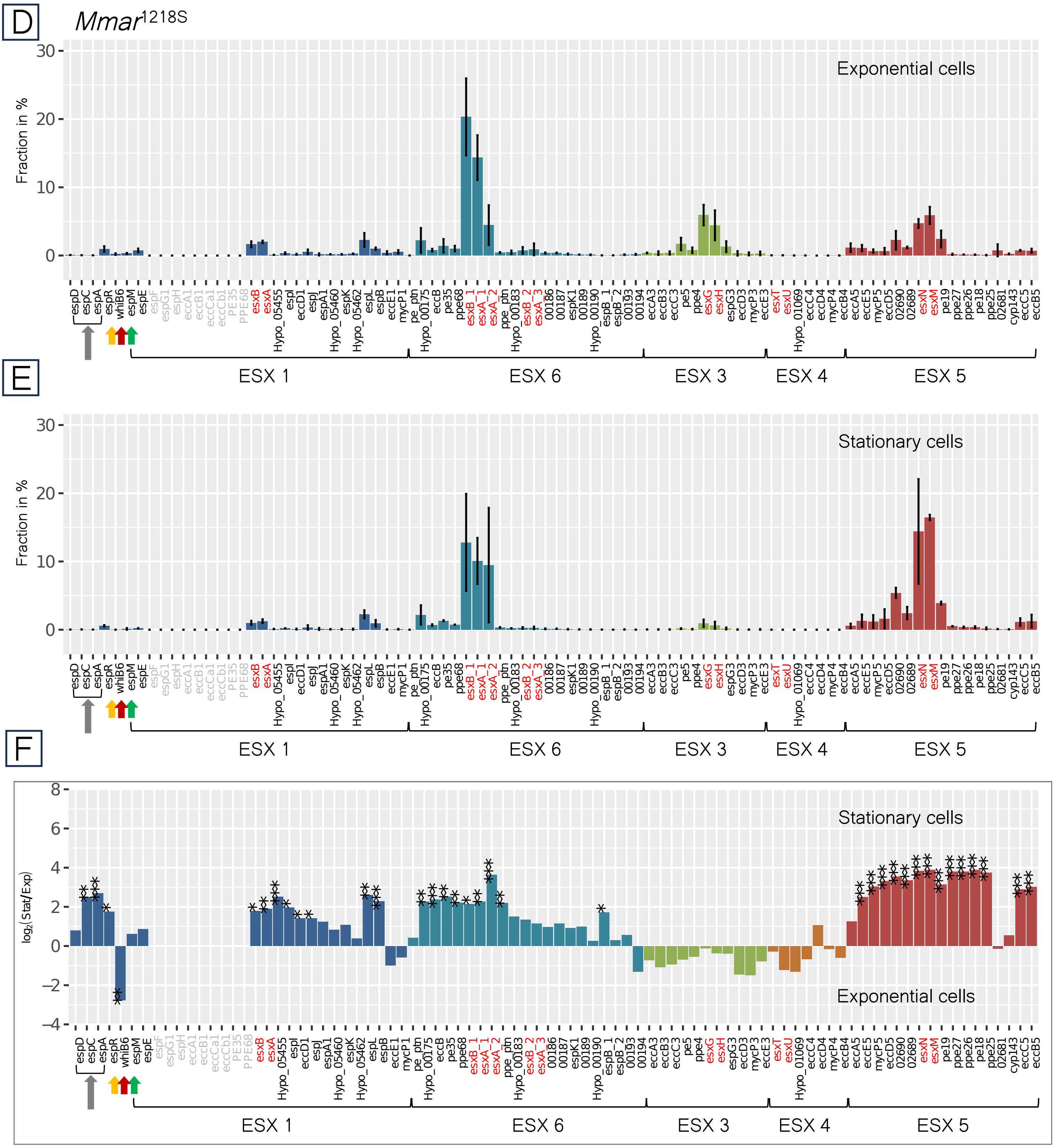

D
E
F
Mmar1218S
Exponential cells
Stationary cells
Fraction in %
log2(Stat/Exp)
espD espC espA espR whiB6 espM espE espF espG1 espH eccA1 eccB1 eccCa1 eccCb1 PE35 PPE68 esxB esxA Hypo_05455 espI eccD1 espJ espA1 Hypo_05460 espK Hypo_05462 espL espB eccE1 mycP1 pe_ptn Hypo_00175 eccB pe35 ppe68 esxB_1 esxA_1 esxA_2 ppe_ptn Hypo_00183 esxB_2 esxA_3 00186 00187 espK1 00189 Hypo_00190 espB_1 espB_2 00193 00194 eccA3 eccB3 eccC3 pe5 ppe4 esxG esxH espG3 eccD3 mycP3 eccE3 esxT esxU Hypo_01069 eccC4 eccD4 mycP4 eccB4 eccA5 eccE5 mycP5 eccD5 02690 02689 esxN esxM pe19 ppe27 ppe26 pe18 ppe25 02681 cyp143 eccC5 eccB5
ESX 1
ESX 6
ESX 3
ESX 4
ESX 5

### Figure S7G -I

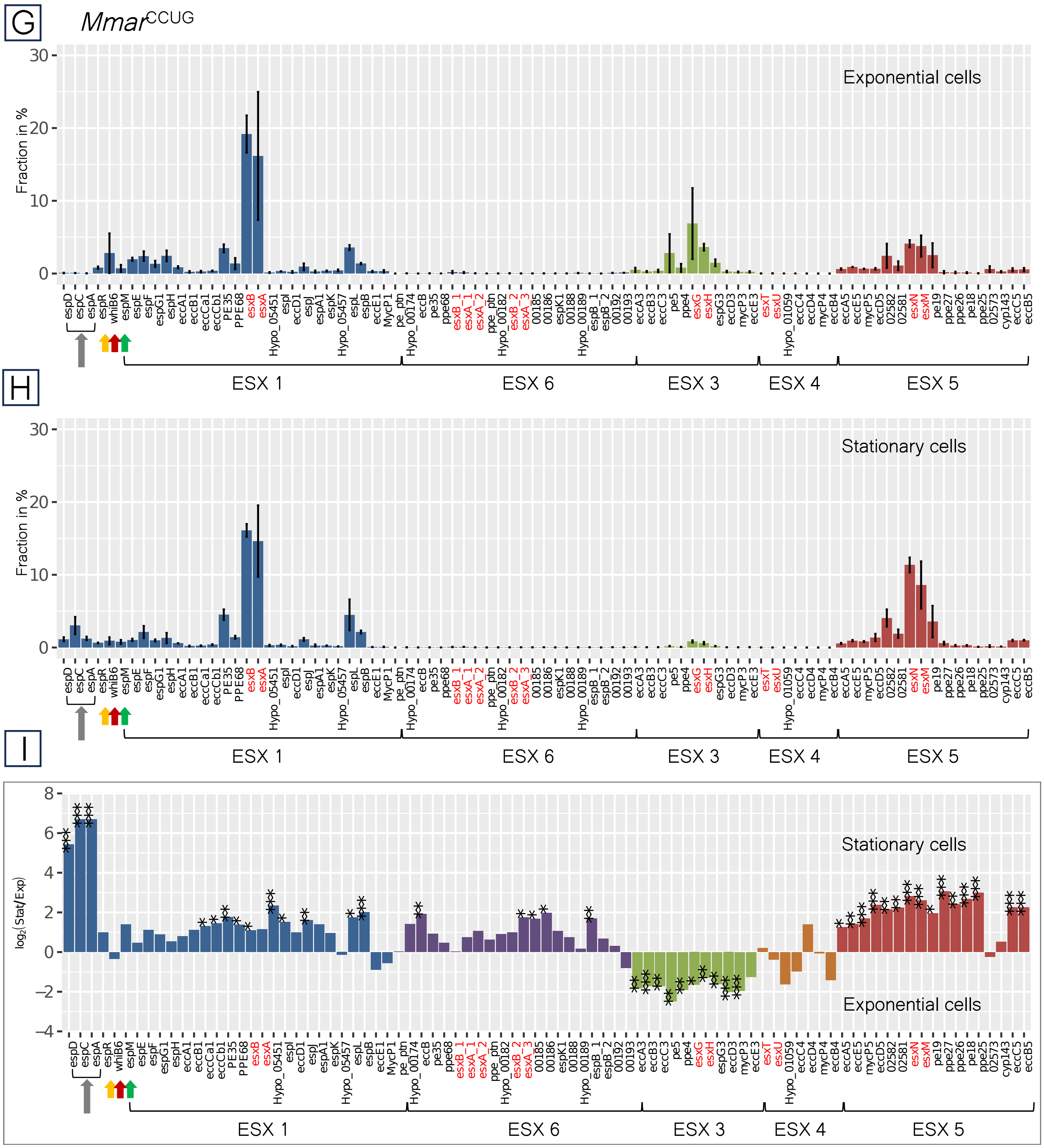
G
Mmar CCUG
Exponential cells
Fraction in %
espD espC espA espR whiB6 espM espE espF espG1 espH eccA1 eccB1 eccCa1 eccCb1 PE35 PPE68 esxB esxA Hypo_05451 espI eccD1 espJ espA1 espK Hypo_05457 espL espB eccE1 MycP1 pe_ptn Hypo_00174 eccB pe35 ppe68 esxB_1 esxA_1 esxA_2 ppe_ptn Hypo_00182 esxB_2 esxA_3 00185 00186 espK1 00188 Hypo_00189 espB_1 espB_2 00192 00193 eccA3 eccB3 eccC3 pe5 ppe4 esxG esxH espG3 eccD3 mycP3 eccE3 esxT esxU Hypo_01059 eccC4 mycP4 eccB4 eccA5 eccE5 mycP5 eccD5 02582 02581 esxN esxM pe19 ppe27 ppe26 pe18 ppe25 02573 cyp143 eccC5 eccB5
ESX 1
ESX 6
ESX 3
ESX 4
ESX 5
H
Stationary cells
Fraction in %
espD espC espA espR whiB6 espM espE espF espG1 espH eccA1 eccB1 eccCa1 eccCb1 PE35 PPE68 esxB esxA Hypo_05451 espI eccD1 espJ espA1 espK Hypo_05457 espL espB eccE1 MycP1 pe_ptn Hypo_00174 eccB pe35 ppe68 esxB_1 esxA_1 esxA_2 ppe_ptn Hypo_00182 esxB_2 esxA_3 00185 00186 espK1 00188 Hypo_00189 espB_1 espB_2 00192 00193 eccA3 eccB3 eccC3 pe5 ppe4 esxG esxH espG3 eccD3 mycP3 eccE3 esxT esxU Hypo_01059 eccC4 mycP4 eccB4 eccA5 eccE5 mycP5 eccD5 02582 02581 esxN esxM pe19 ppe27 ppe26 pe18 ppe25 02573 cyp143 eccC5 eccB5
ESX 1
ESX 6
ESX 3
ESX 4
ESX 5
I
Stationary cells
Exponential cells
log2(Stat/Exp)
espD espC espA espR whiB6 espM espE espF espG1 espH eccA1 eccB1 eccCa1 eccCb1 PE35 PPE68 esxB esxA Hypo_05451 espI eccD1 espJ espA1 espK Hypo_05457 espL espB eccE1 MycP1 pe_ptn Hypo_00174 eccB pe35 ppe68 esxB_1 esxA_1 esxA_2 ppe_ptn Hypo_00182 esxB_2 esxA_3 00185 00186 espK1 00188 Hypo_00189 espB_1 espB_2 00192 00193 eccA3 eccB3 eccC3 pe5 ppe4 esxG esxH espG3 eccD3 mycP3 eccE3 esxT esxU Hypo_01059 eccC4 mycP4 eccB4 eccA5 eccE5 mycP5 eccD5 02582 02581 esxN esxM pe19 ppe27 ppe26 pe18 ppe25 02573 cyp143 eccC5 eccB5
ESX 1
ESX 6
ESX 3
ESX 4
ESX 5

### Figure S7J -L

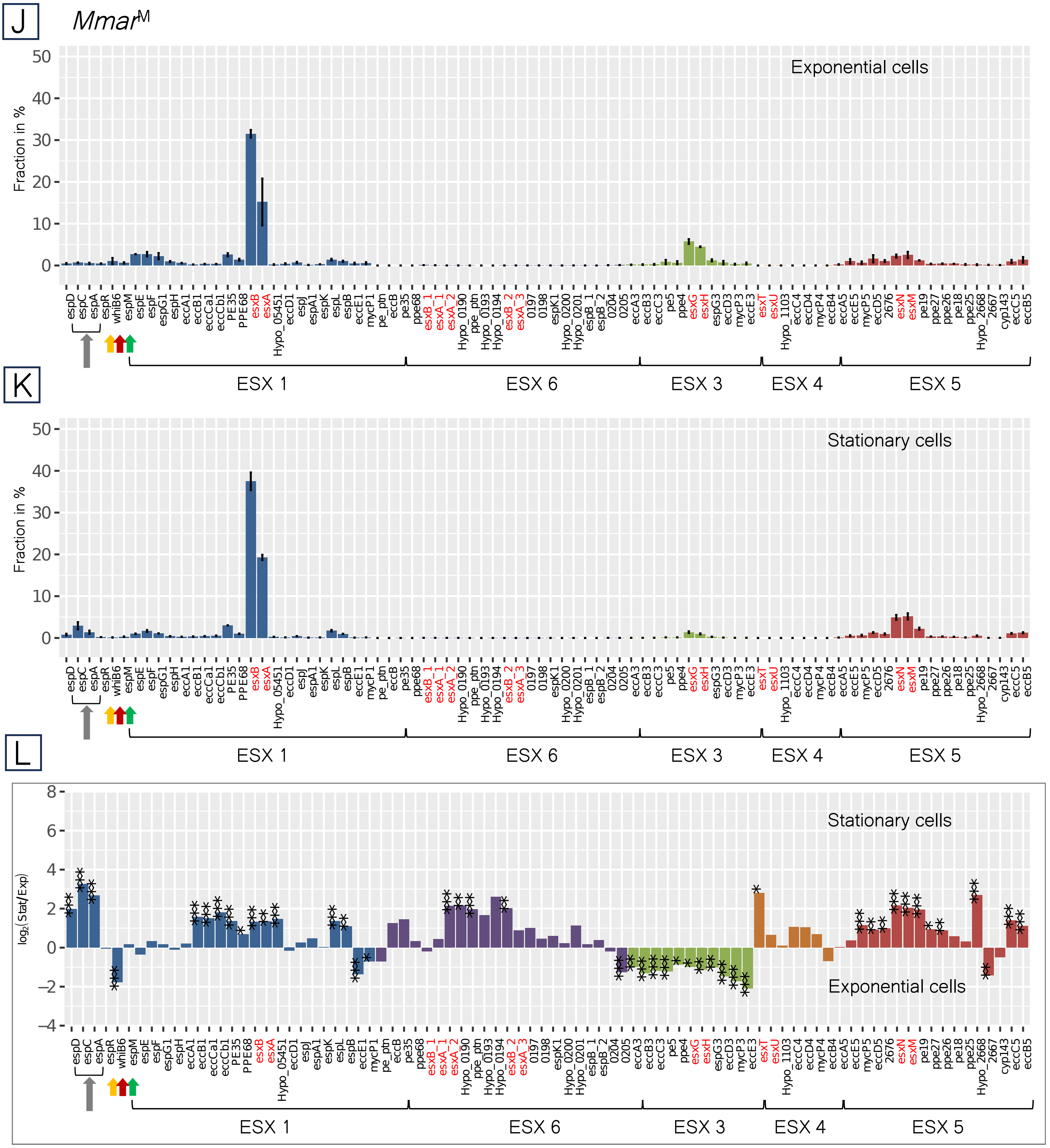
J
Mmar^M
Exponential cells
Fraction in %
K
Stationary cells
Fraction in %
L
Stationary cells
Exponential cells
log2(Stat/Exp)
ESX 1
ESX 6
ESX 3
ESX 4
ESX 5

**Figure S8** RNA-Seq reads showing transcription of *espF-esxB* in 1218S. The 1218R gene annotations/identity used as reference.

### Fig S8 RNA-Seq – 1218S *espF-esxB*

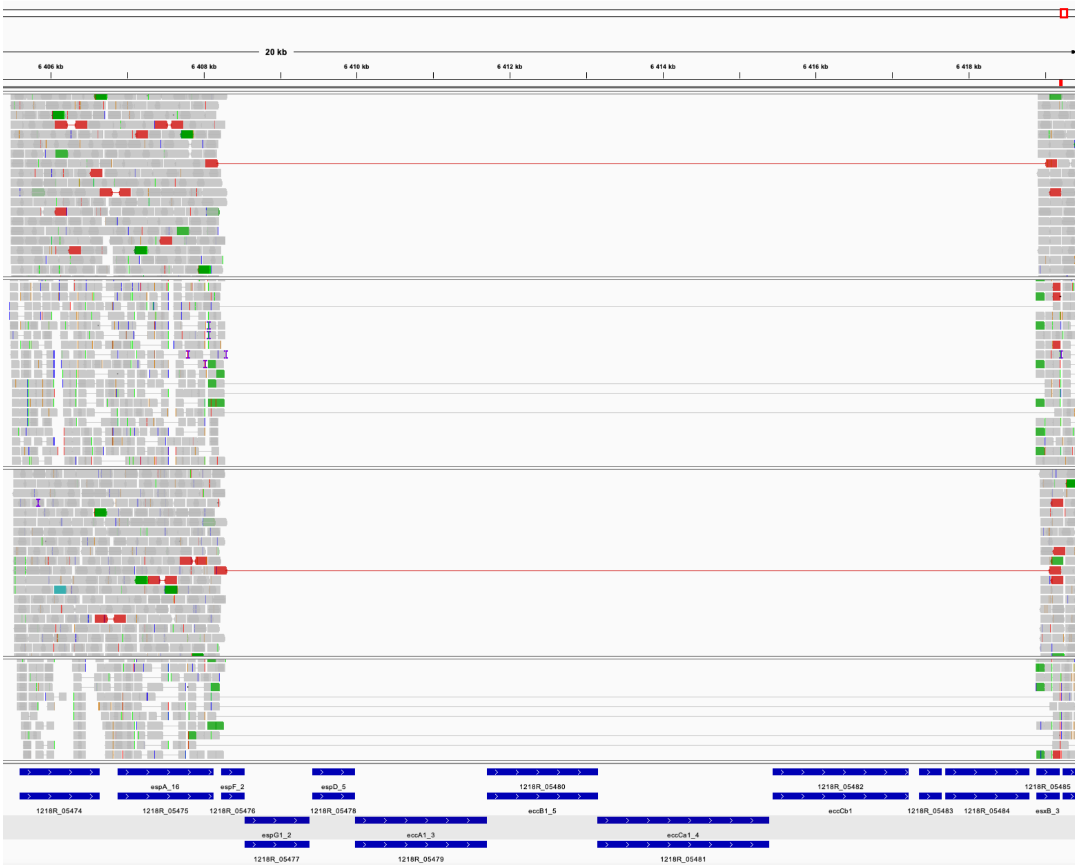


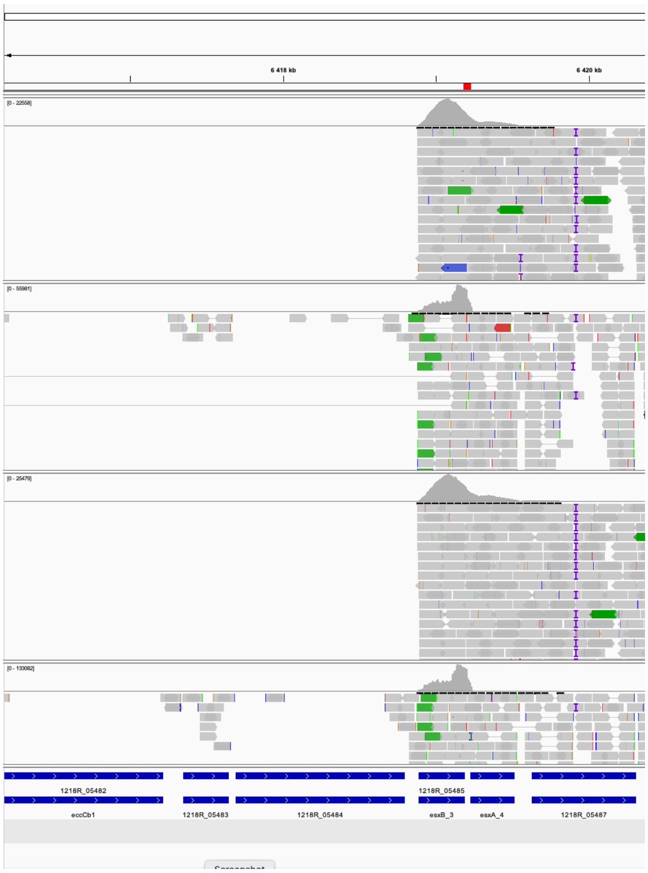

**Figure S9** Transcript levels of LOS genes in 1218R and 1218S.

(a) Transcript levels (distribution, expressed in TPM values) in exponentially growing and stationary 1218R cells.

(b) Comparing transcript levels in exponentially growing and stationary 1218R cells. Negative $\log_2$-values suggest that the corresponding mRNA is more abundant in exponential cells while a positive value suggests higher levels in stationary 1218R cells.

(c) Transcript levels (distribution, expressed in TPM values) in exponentially growing and stationary 1218S cells.

(d) Comparing transcript levels in exponentially growing and stationary 1218S cells. Negative $\log_2$-values suggest that the corresponding mRNA is more abundant in exponential cells while a positive value suggests higher levels in stationary 1218S cells. The 1218R gene annotations/identity used as reference.

Statistical significance, see Materials and Methods; *p-value < 0.05; **p-value < 0.01; ***p-value < 0.001.

### Fig S9A-B

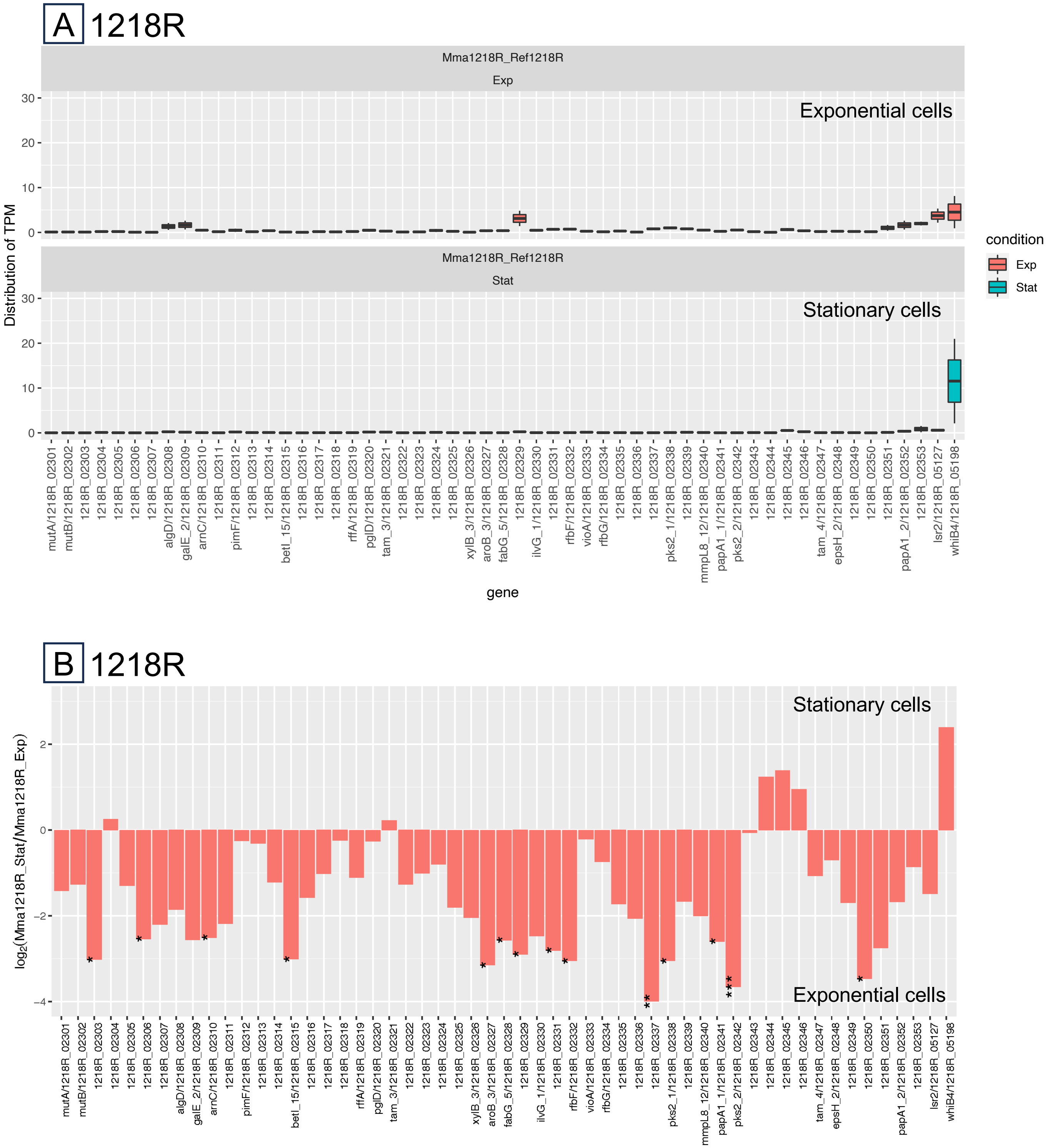
A
1218R
Mma1218R_Ref1218R
Exp
Exponential cells
Mma1218R_Ref1218R
Stat
Stationary cells
Distribution of TPM
condition
Exp
Stat
gene
mutA/1218R_02301
mutB/1218R_02302
1218R_02303
1218R_02304
1218R_02305
1218R_02306
1218R_02307
algD/1218R_02308
galE_2/1218R_02309
arnC/1218R_02310
1218R_02311
pimF/1218R_02312
1218R_02313
1218R_02314
betI_15/1218R_02315
1218R_02316
1218R_02317
1218R_02318
rffA/1218R_02319
pglD/1218R_02320
tam_3/1218R_02321
1218R_02322
1218R_02323
1218R_02324
1218R_02325
xylB_3/1218R_02326
aroB_3/1218R_02327
fabG_5/1218R_02328
1218R_02329
ilvG_1/1218R_02330
1218R_02331
rfbF/1218R_02332
vioA/1218R_02333
rfbG/1218R_02334
1218R_02335
1218R_02336
1218R_02337
pks2_1/1218R_02338
1218R_02339
mmpL8_12/1218R_02340
papA1_1/1218R_02341
pks2_2/1218R_02342
1218R_02343
1218R_02344
1218R_02345
1218R_02346
tam_4/1218R_02347
epsH_2/1218R_02348
1218R_02349
1218R_02350
1218R_02351
papA1_2/1218R_02352
1218R_02353
lsr2/1218R_05127
whiB4/1218R_05198
B
1218R
Stationary cells
Exponential cells
log2(Mma1218R_Stat/Mma1218R_Exp)

### Fig S9C-D

C 1218S

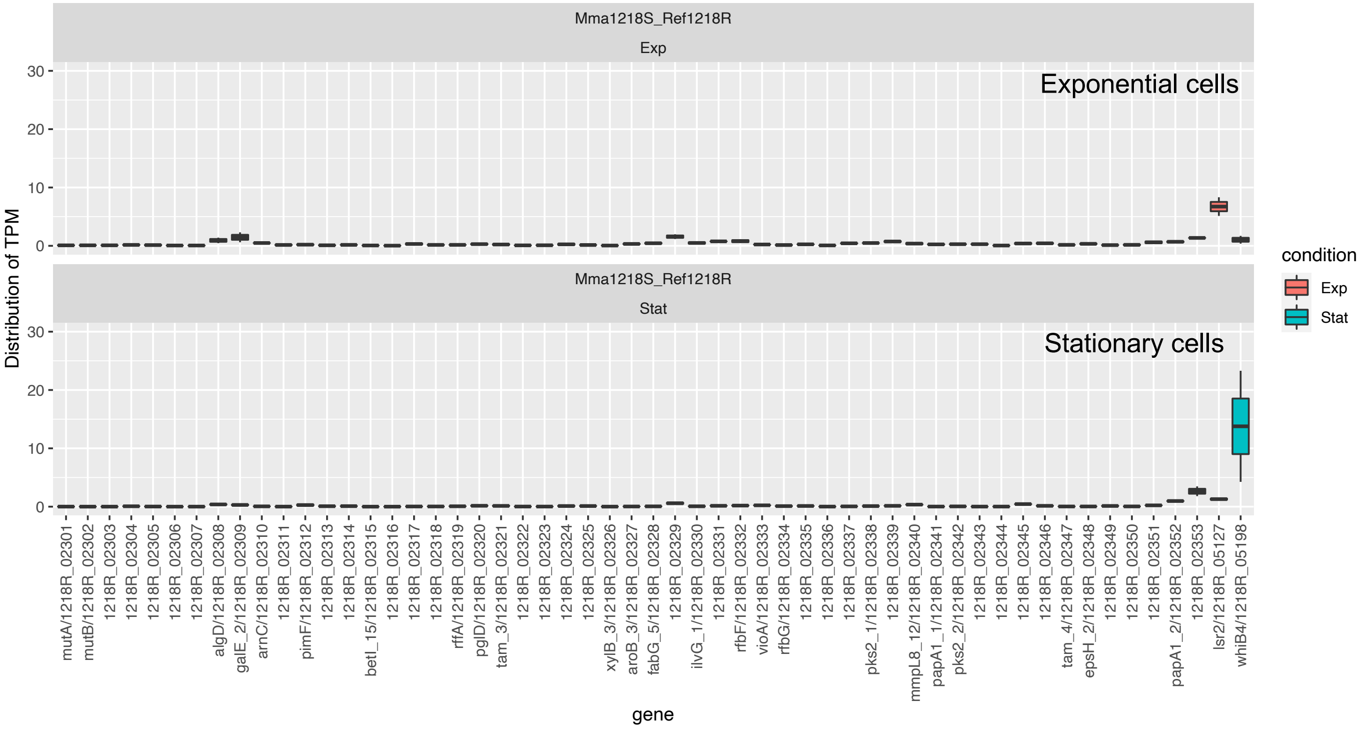


D 1218S

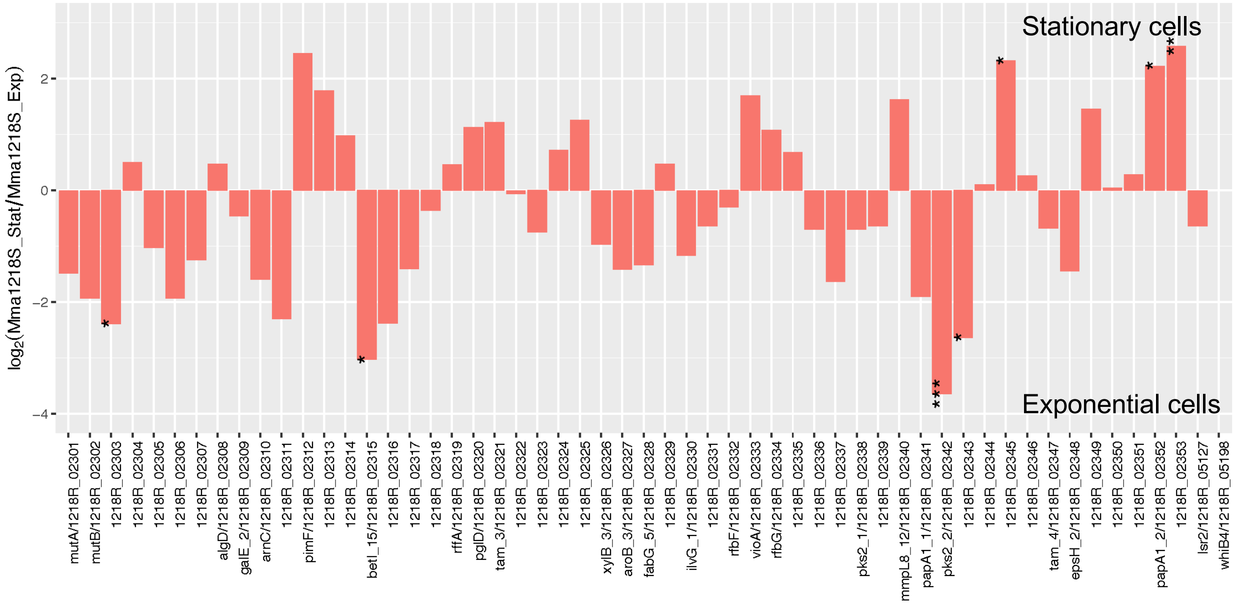

**Figure S10** Ms1 RNA gene transcript levels.

Ms1 RNA levels in exponentially growing and stationary 1218R and 1218S cells (left panel); *Mmar*$^{CCUG}$ and *Mmar*$^{M}$ (see also Figure 7a). Levels with postive values correspond to higher levels in stationary cells.

Statistical significance, see Materials and Methods; *p-value < 0.05; **p-value < 0.01; ***p-value < 0.001.

Fig S10

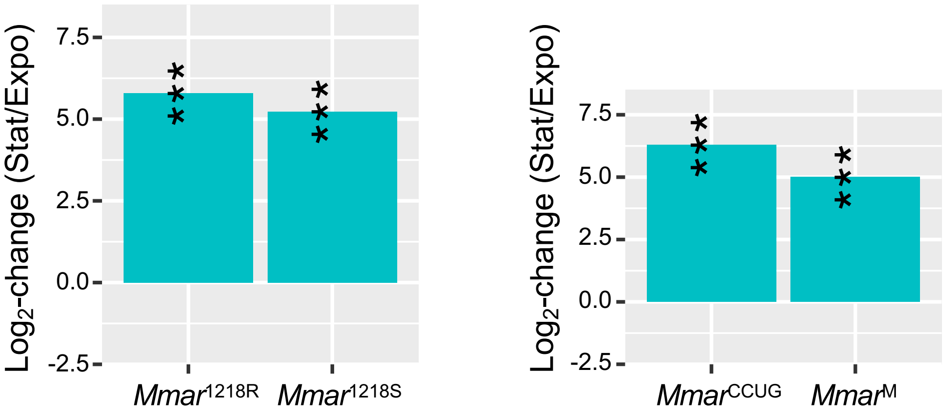

Log2-change (Stat/Expo)
7.5
5.0
2.5
0.0
-2.5
***
***
Mmar1218R
Mmar1218S
Log2-change (Stat/Expo)
7.5
5.0
2.5
0.0
-2.5
***
***
MmarCCUG
MmarM

**Figure S11** Growth on 7H10 media after transformation of 1218R and 1218S with the empty control plasmid (pBS401) or with pBS401$^{espF\text{-}H}$ as indicated (uninduced, top panels; induced with tetracycline, lower panels). For details see main text, Figure 8 and Materials and Methods.

### Fig S11

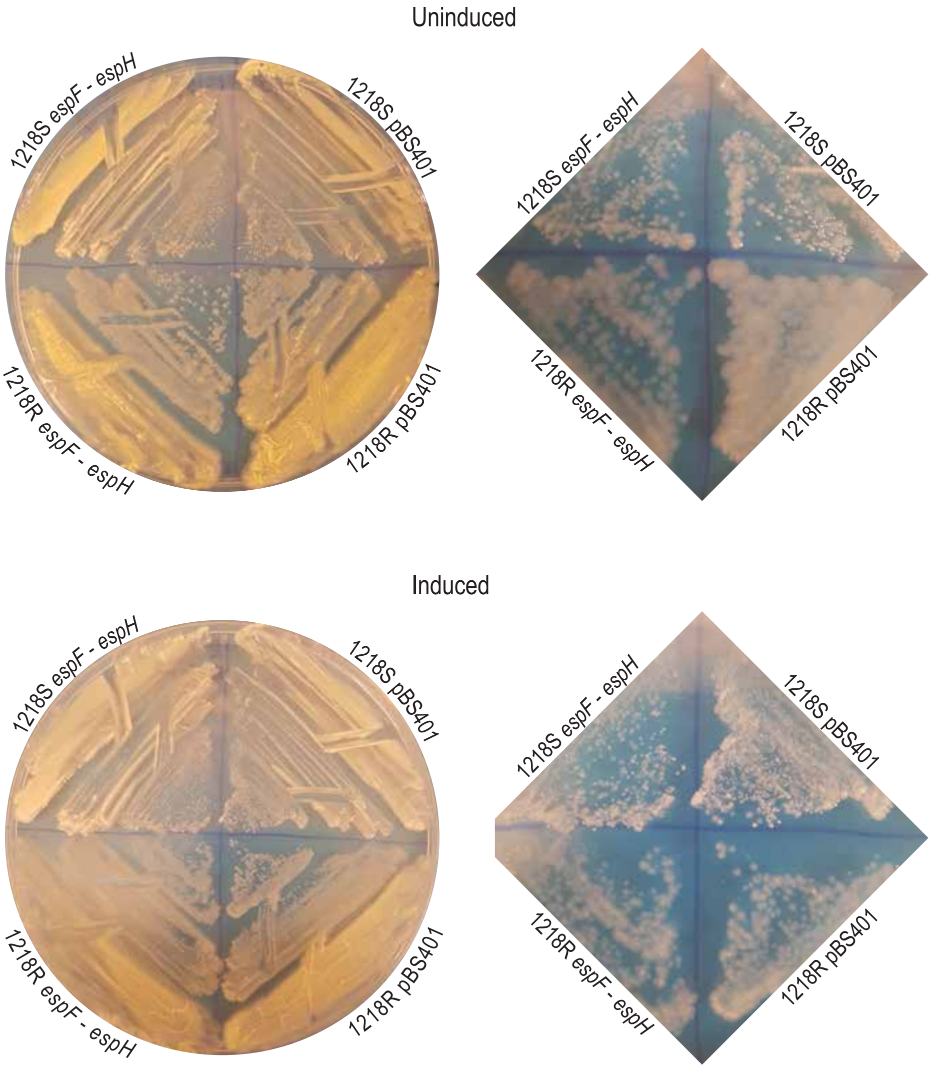

Uninduced
1218S espF - espH
1218S pBS401
1218R espF - espH
1218R pBS401
1218S espF - espH
1218S pBS401
1218R espF - espH
1218R pBS401
Induced
1218S espF - espH
1218S pBS401
1218R espF - espH
1218R pBS401
1218S espF - espH
1218S pBS401
1218R espF - espH
1218R pBS401
